# The mutational footprints of cancer therapies

**DOI:** 10.1101/683268

**Authors:** Oriol Pich, Ferran Muiños, Martijn Paul Lolkema, Neeltje Steeghs, Abel Gonzalez-Perez, Nuria Lopez-Bigas

## Abstract

Some cancer therapies damage DNA and cause mutations both in cancer and healthy cells of the patient^1^. These therapy-induced mutations may underlie some of the long-term and late side effects of the treatment, such as mental disabilities, organ toxicities and secondary neoplasms. Currently we ignore the mutation pattern and burden caused by different cancer treatments. Here we identify mutational signatures, or footprints of six widely-used anti-cancer therapies with the study of whole-genomes from more than 3500 metastatic tumors originated in different organs. These include previously known and new mutational signatures generated by platinum-based drugs, and a novel signature of treatment with nucleoside metabolic inhibitors. Exploiting these mutational footprints, we estimate the contribution of different treatments to the mutation burden of tumors and their risk of causing coding and likely driver mutations in the genome. In summary, the mutational footprints identified here open a window to precisely appraise the mutational risk of different cancer therapies to understand their late side effects.

## Introduction

Tumors start and evolve as a result of the interplay between somatic mutations and selective constraints faced along their development^1^. All cells of our body accumulate somatic variants contributed by both endogenous and external mutational processes. Each of these processes contribute preferentially certain types of nucleotide changes in specific sequence contexts. The repertoire of somatic mutations that a cell has received can thus be used to identify mutational signatures, which represent the mutational processes that have been active through its history^2–5^.

Many chemotherapies, which are still the workhorse in the treatment of primary tumors, cause DNA damage or change the pool of nucleotides and hence target both cancer and non-cancer cells of patients^6,7^. While many tumor and healthy cells affected by the DNA damage generated by these drugs will die, others can survive. In the offspring of the surviving cells, at least part of the original damage will be converted into mutations (Fig. 1a). Therefore, chemotherapies may contribute mutations to the tumor, and to healthy tissues of the patient’s organs, which likely underpin some of the long-term secondary effects caused by these treatments^8–10^. As with other mutational processes nucleotide changes caused by chemotherapy agents will leave an imprint in the genomes of treated cells, which can be detected as specific mutational signatures. Indeed, platinum-based drugs^6,7,11,12^, temozolomide^2,13^ and radiation treatments^14^ have already been associated to specific mutational signatures and the mutational footprints of some of them have been confirmed experimentally^6^. However, virtually nothing is known about the effects of other chemotherapeutic treatments on the mutational pattern of somatic and germ cells, since mutational signatures have been studied majoritarily across primary chemotherapy-naive tumors. As a result, we still ignore the specific mutational profile and burden caused by most chemotherapies to patient’s cells. This is of crucial importance to understand the resistance of tumors to chemotherapies and to explain and predict the long-term effects of these treatments to the patient. Here, using the somatic mutations present in 3506 metastatic tumors we identify the mutational footprints left by five chemotherapeutic agents and radiotherapy. Using these specific footprints we then estimate the contribution of these chemotherapies to the mutational burden of these tumors, in comparison to that of endogenous mutations contributed by the natural aging process. Finally, we assess the risk posed by each of these therapies to generate coding mutations and potential cancer driver mutations. We regard these two measures as the “mutational toxicity” of these four chemotherapeutic agents in different tissues.

**Figure 1.**
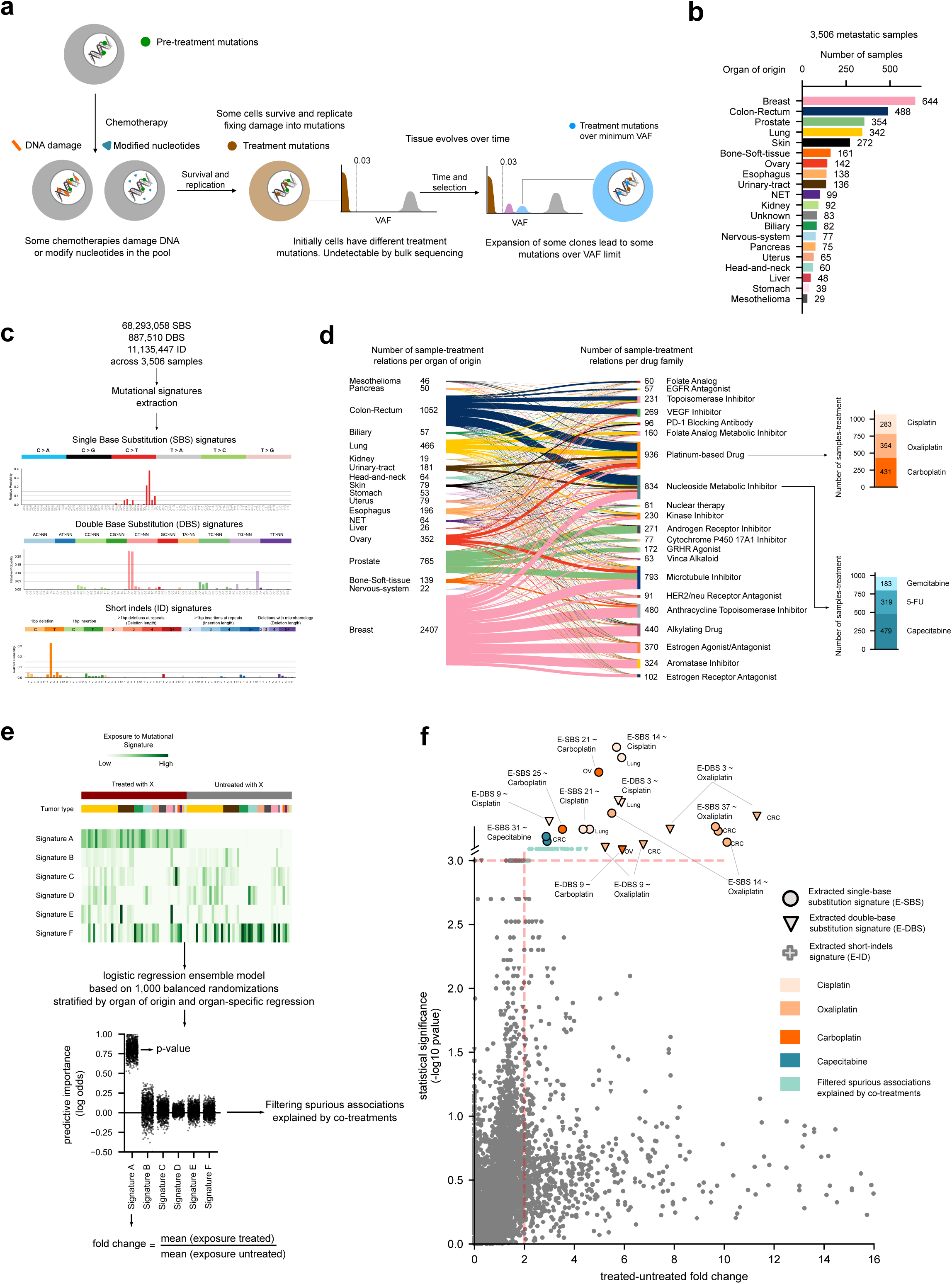
Uncovering mutational signatures associated to anti-cancer treatments. (a) Schematic representation of the evolution of a tumor upon treatment. Somatic cells bear mutations at the time of treatment contributed by different mutational processes (gray). Some treatments directly damage the DNA, while others alter the pool of nucleotides (red bars and hexagons, respectively), potentially causing the death of a large share of the cells. The surviving cells (after rounds of DNA replication) bear mutations caused by the unrepaired DNA damage, the consequences of misincorporated nucleotide analogs (green) or introduced by error-prone polymerases during repair. These treatment mutations are private of each surviving cell after the first round of replication, with low variant allele frequencies (VAF), so that bulk sequencing would be unable to detect them. However, pre-treatment somatic mutations will be present in larger fractions of the surviving cells (higher VAF). Some of the surviving cells with advantages over their neighbors (e.g., a set of driver mutations) will grow faster to replenish the niche opened by the massive death of tumor cells. Over time, the progeny of these faster-growing cells will represent a larger fraction of the population, effectively amplifying their genetic material within the tumor pool. At the time of biopsy of the metastasis, the VAF of treatment mutations present in the original surviving cells will raise above the threshold of detection of bulk sequencing. (b) Composition of the metastatic tumors cohort grouped by the organ/tissue of origin of the primary. The code to color tumors originated from different organs is used in subsequent figures. NET: Neuroendocrine tumors. (c) Three examples of SBS, DBS and ID signatures extracted from the cohort using the SignatureAnalyzer, represented as their mutational profiles. The profiles of all signatures identified with the SignatureAnalyzer and the SigProfiler appear in Supplementary Note 1. (d) Names of FDA families of anti-cancer treatments administered to patients with primary tumors from different origins and numbers of patients who received each of them. Stacked barplots at the right: number of metastatic tumors exposed to individual drugs belonging to two of the most widely employed families. (All numbers in this panel correspond to sample-treatment pairs, which due to complex regimens add up to more than the total number of tumors of each organ of origin in the cohort.) (e) Schematic representation of the logistic regression ensemble approach employed to identify treatment-associated mutational signatures (Methods). The toy heatmap in the top panel represents a matrix of activity of mutational signatures identified across tumors. Tumors are identified by the organ of their primary (colors immediately above the heatmap), and those exposed to an exemplary treatment (X) are labeled red above the organ-of-origin annotation, while tumors not treated with X are labeled gray. One thousand balanced subsets of tumors exposed and not exposed to X are randomly sampled from this matrix stratified by organ of origin. One regression model is applied for each treatment under analysis. The effect size given by each regression model for each signature is computed as the fold-change between the mean exposure of treated and untreated tumors. A p-value as a measure of statistical significance of the association is also computed. Finally, the results of these regressions are filtered to discard spurious associations potentially due to co-treatment regimens. Details of the methodology and validation of its performance using synthetic datasets are in Supplementary Note 2. (f) Treatment-associated mutational signatures (among the ones extracted with SignatureAnalyzer). Each dot represents the effect-size and the p-value of the association between one signature and one treatment detected through one regression model. Associations deemed significant (effect-size>2 and p-value<0.001, values that correspond to the vertical and horizontal dashed lines, respectively) are colored according to the treatment and shaped following the type of variant (SBS, DBS, ID) of the mutational signatures. (Since p-value=0.001 corresponds to the maximum resolution of the test, significantly associated signatures are placed above the truncated y-axis at arbitrary vertical positions.) Associations detected in organ-specific regressions are thus denoted; all other associations are detected across the entire pan-metastatic adult cohort. Detailed results appear in Table S2. SigProfiler-extracted signatures that appear associated to treatment appear in Figs. S3 and S4. CRC: colon-rectum, OV: ovary.

## Results

### Identification of mutational signatures associated to anti-cancer therapies

We reasoned that the analysis of available metastasis of patients who have undergone chemotherapy treatment regimens provide a good opportunity to identify the mutational footprint of anti-cancer agents. Driven by the clonal expansion experienced by these tumors, treatment-induced mutations, if present, would appear at values of variant allele frequency (VAF) that render them detectable through sequencing (Fig. 1a). We thus analyzed a cohort of 3506 metastatic tumor samples, sequenced at the whole-genome level^15^. These samples were taken from patients who previously suffered from primary tumors originated in at least 19 known different organs or tissues, ranging from 644 carcinomas of breast to 29 mesotheliomas (Fig. 1b, Table S1). We used SignatureAnalyzer^16,17^ and SigProfiler^2,18^, two widely-employed methods based on different principles that address the non-negative matrix factorization (NMF) problem (and a third non-NMF method across tumors of colorectal origin) to extract the mutational signatures active across these metastatic samples (Methods). Mutational signatures of single base substitutions (SBS), doublet base substitutions (DBS) and indels (ID) were extracted separately (Fig. 1c and Supp. Note 1). Some of the signatures discovered in the tumors of the cohort have been previously identified^2–4,6,18–20^, and thus to refer to them, we employ their known etiologies (e.g., aging signature).

We first manually curated the information of treatment exposure of the patients under study (Fig. 1d). In this cohort, 2124 tumor samples were taken from patients to whom treatments consisting of one or more of 206 drugs of 58 distinct FDA classes were administered. These drugs were given to the patients 2.33 years in median prior to the obtention of the biopsies of the metastases (Fig. S1). Platinum-based drugs (cisplatin, oxaliplatin and carboplatin) were the class most frequently employed to treat the patients in the cohort. The choice of chemotherapy was primarily guided by the organ of origin of the tumors, and most patients (1848) received more than one drug in the course of the treatment, either in a combined or sequential regimen (Fig. S2).

To discern the mutational signatures among those identified in this cohort that constitute the footprint of chemotherapies, we designed an *ad hoc* logistic ensemble regression model approach (hereinafter *regression model*). This model identifies associations between the exposure of metastatic tumors in the cohort to chemotherapeutic treatments and the activity of the identified mutational signatures (Fig. 1e). This approach controls for potential associations between treatments and organ-of-origin of the tumors, and reliably identifies signatures associated to the treatments, as demonstrated on mutations injected in samples of synthetic datasets (Fig. S3a, Table S2, Methods, Supp. Note 2). The approach also controls for potential spurious associations due to simultaneous treatments with several drugs –e.g., a signature that appears related to bevacizumab, but which was actually associated to concomitant oxaliplatin (Supp. Notes 1 and 2). We run pan-cancer and organ-specific regressions to gain sensitivity to identify potential associations missed across the entire cohort due to dilution effects. As a result (Fig. 1f), we identified seven mutational signatures (five SBS signatures and two DBS signatures) associated to 4 treatments (with overall or organ-specific effect size > 2 and p-value < 0.001). Equivalent sets of signatures were obtained with the two extraction methods (i.e. SigProfiler and SignatureAnalyzer - Figure S3 and S4) which shows that the chemotherapy mutational footprints detected are robust to the singularities of different signature extraction methods (Supp. Note 1).

### The mutational footprints of six anti-cancer therapies

Four SBS and two DBS signatures constituted the footprint of three platinum-based drugs (Fig. 2a, S3, and S4), with two SBS signatures associated to more than one drug and both DBS signatures associated to the three platinum-based drugs. One signature (with cosine similarity 0.954 to the SBS Carboplatin/Cisplatin signature) had been previously identified as the footprint of the treatment with cisplatin or carboplatin^5^. On the other hand, an oxaliplatin-related signature (SBS Oxaliplatin) is detected in this cohort for the first time. Platinum-based drugs-associated signatures exhibit transcriptional strand asymmetry (Methods), i.e., lower activity in the template strand of transcribed genes (Fig. 2b and S3c). These drugs generate DNA adducts that cause RNA polymerases to stall and recruit the transcription-coupled nucleotide excision repair^21,22^ machinery, yielding this asymmetric activity of its mutational footprint between strands.

**Figure 2.**
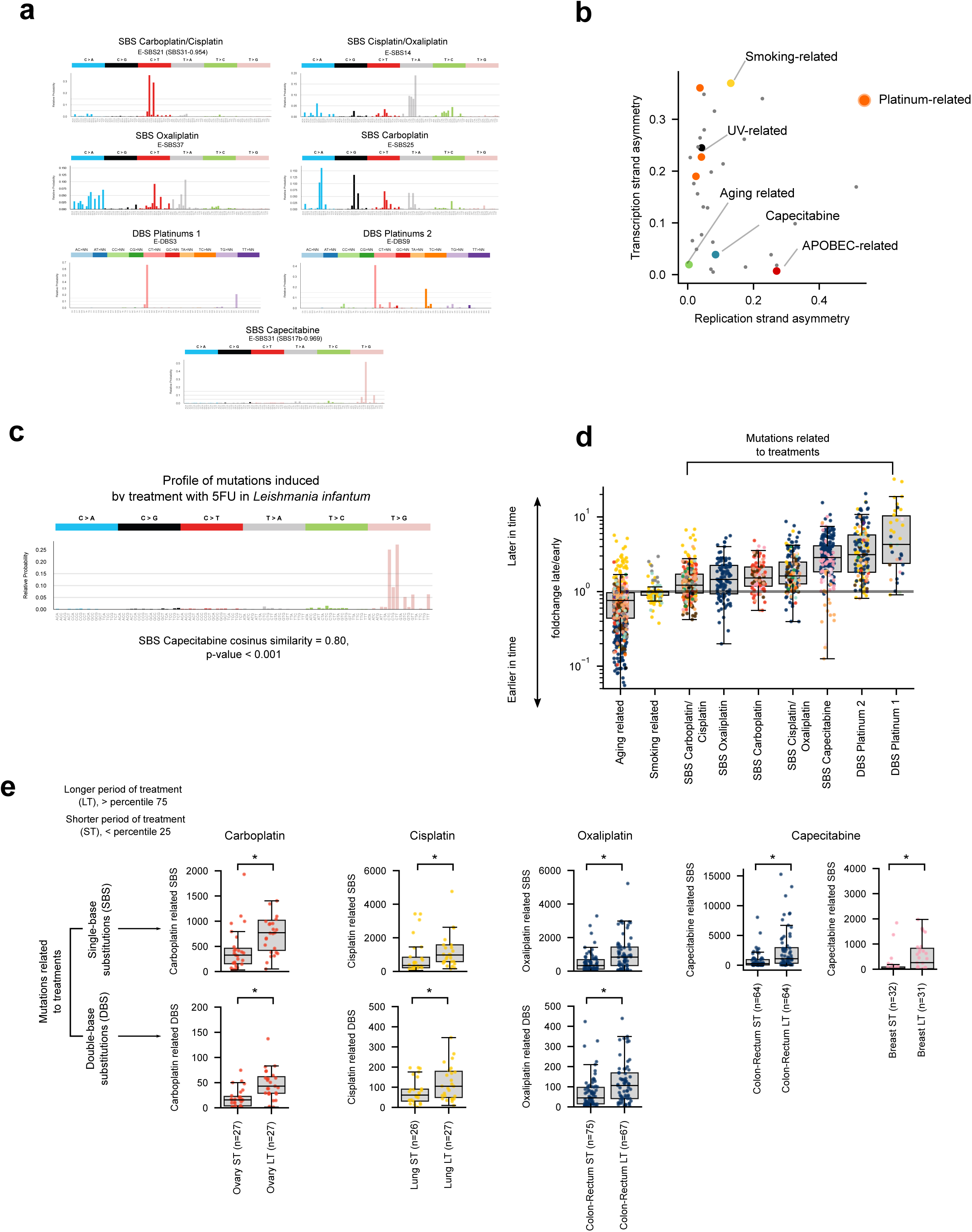
Characteristics of treatment-associated mutations. (a) Mutational profiles (frequency of each tri-nucleotide change) of the seven SBS and DBS signatures (in the SignatureAnalyzer extraction) associated to treatments through the logistic regression ensemble. *Ad hoc* names following their etiology given to each signature (above the plots) are used throughout the manuscript to refer to them. (b) Strand asymmetry of selected SignatureAnalyzer-extracted signatures and other signatures active across healthy and tumoral somatic cells. Each dot corresponds to a signature (colored following the code in Fig. 1e), with the abscisa representing its replication strand asymmetry (the unbalance of mutations between leading and lagging strand), and its ordinate, the transcriptional strand asymmetry (the unbalance of mutations between template and non-template strands). The strand asymmetry of SigProfiler-extracted signatures appears in Fig. S3c. (c) Profile (frequency of each tri-nucleotide change) of the private mutations (not present in parental cell) of 5 mutant *Leishmania infantum* strains treated with 5-FU. It is very similar to the SBS capecitabine signature shown in panel (a) (cosine similarity 0.8, p-value < 0.001, see Methods). (d) Mutations contributed by signatures associated to treatments (termed as in Fig. 2a) are enriched for later substitutions (higher late-to-early fold-change), in comparison to signatures active earlier in the lifetime of the patients (e.g., aging and smoking-related signatures). Each tumor is represented as a dot colored following the code of organ-of-origin presented in Figure 1a. SignatureAnalyzer-extracted signatures are represented in the Figure. Equivalent graph for SigProfiler-extracted signatures appear in Figure S7b. (e) The mutation load contributed by treatment-associated signatures correlates with the duration of the period of exposure to the treatment (extraction with SignatureAnalyzer). Comparison of the distribution of the number of SBS (upper row) and DBS (lower row) of signatures associated to each drug across ST and LT tumors of organ of origin with sufficient mutations to carry out the comparison. In every case, LT tumors possess significantly more mutations than ST tumors. (Equivalent graphs for SigProfiler-extracted signatures appear in Fig. S7d.)

One already known ID signature (ID11; Supplementary Note 1) associated to radiation treatment^14^ appeared close to significance (p-value < 0.01, effect size < 2). Their activity is higher in Homologous Recombination (HR)-defective than HR-proficient tumors (Fig. S5a). Among HR-proficient tumors, the irradiated ones exhibit significantly higher activity of the irradiation-signature than the non-irradiated ones. The regression model failed to detect a known SBS signature associated to treatment with temozolomide (TMZ)^2,13^. Searching specifically for this signature we found that it appears in 5 TMZ-exposed samples, but is lacking in other 17 equally TMZ-treated tumors, thus rendering the association given by the regression model non-significant. The difference is explained by protein-affecting mutations in genes of the MMR pathway in the three tumors of the first group (Fig. S5b). Four MMR-deficient tumors with no annotated TMZ treatment show a relatively high activity of the TMZ-associated signature.

We also discovered a previously unknown SBS signature significantly associated to the treatment with capecitabine (SBS Capecitabine). Capecitabine is a nucleoside metabolic inhibitor metabolized to 5-fluorouracil (5-FU), a chemotherapy to which 319 tumor samples in the cohort were exposed (Fig. 1d). Not surprisingly, thus this capecitabine-associated signature is also active in 5-FU-exposed tumors (Fig. S6c). A regression analysis that compares 5-FU-exposed and unexposed samples when capecitabine-exposed tumors are removed from the latter group identifies its exposure as associated to the SBS Capecitabine across the cohort and in tumors of breast and colorectal origin independently (Table S2). Furthermore, the association is stronger and reaches higher statistical significance when the activity of the SBS Capecitabine across samples exposed to either drug is compared to unexposed tumors (Table S2). To obtain experimental validation of the association of capecitabine/5-FU to this signature, we analyzed mutations in five resistant cultures of *Leishmania infantum* exposed to 5-FU^23^. This revealed a profile dominated by CTC>CGC and CTT>CGT mutations, as in the SBS Capecitabine (p-value < 0.001; Figs. 2c and S6), thus confirming the etiology of the signature identified in tumors. Inside cells, 5-FU is converted to 5-fluorodeoxyuridine monophosphate, an inhibitor of thymidylate synthase, and 5-fluorodeoxyuridine triphosphate (FdUTP). As a result, the pool of pyrimidines triphosphate becomes acutely depleted for thymines and enriched for FdUTPs, which polymerases can incorporate into the DNA^24,25^. The SBS Capecitabine (or, more accurately, SBS Capecitabine/5-FU) signature exhibits a mutational profile very similar to the known signature 17b (cosine similarity 0.97) –proposed to be caused by oxidative damage to DNA bases in certain tissues, such as esophagus and stomach^26^. Both, the SBS Capecitabine/5-FU and the 17b signatures co-exist in the tumors of the cohort according to the two (three for colorectal tumors) methods of signature extraction employed (Supp. Note 1). Nevertheless, while the previously reported 17b signature occurs ubiquitously across gastric and esophageal cancers, the SBS Capecitabine/5-FU signature is active in tumors exposed to the drug (Fig. S6c). In summary, we propose that this novel signature constitutes the mutational footprint of the treatment with 5-FU or capecitabine.

### Characteristics of therapy-associated mutations

We hypothesized that mutations contributed by therapy-associated signatures would exhibit certain specific properties that differ from those contributed by many endogenous mutational processes. For example, treatment-associated mutations appear only upon exposure to the chemotherapies (Fig. S7a), which is a late event in the evolution of tumors. Thus, we computed the relative time of appearance (between 0, or closest to the metastasis biopsy, and 1, earliest in tumor development) of SBS across the 3506 tumor samples^27^, and classified the SBS in each tumor as early or late. Then, for each tumor we computed the enrichment for late variants (late-to-early fold-change) among the SBSs contributed by each signature. As expected, SBS contributed by drug-associated signatures are enriched for late variants relative to others contributed by signatures that are active early or throughout the evolution of the tumors (Fig. 2d and S7b). Mutations contributed by drug-associated signatures also tend to be subclonal (Fig. S7c). This is consistent with treatment-associated mutations being late, and occurring randomly across tumor cells. Several tumor cells surviving the treatment with different mutations may subsequently give rise to different clones of the metastases (Figure 1a).

Furthermore, we reasoned that more mutations contributed by drug-associated signatures should appear in metastatic tumors from patients who have been under treatment for longer periods of time, or who have received more courses of the same treatment. We computed the duration of the overall period of exposure to a drug of tumor samples taken from patients exposed to platinum-based drugs or capecitabine/5-FU as the difference between the annotated end and beginning of the patients’ treatment with the drug (Fig. S1a). The 25% of tumors with longest period of exposure to therapies (LT) exhibit significantly higher burden of mutations (SBS and DBS) contributed by treatment-associated signatures than the 25% of tumors with shortest period of exposure (ST; Fig. 2d, S7d). These disparities contrast with the uniform burden of mutations contributed by the aging signature with respect to the period of exposure of the tumors to the drugs (Fig. S7e).

### The mutation burden of anti-cancer therapies in metastatic tumors

Chemotherapeutic agents that cause DNA damage, such as platinum-based drugs and capecitabine/5-FU have the potential to cause mutations in both tumor and healthy cells. We reasoned that the identification of their mutational footprint carried out in this work provides an opportunity to estimate their mutational toxicity across metastatic tumors of different origin. This constitutes a proxy of the mutational toxicity of chemotherapies in healthy tissues.

As a first estimate of the mutational toxicity of chemotherapies, we computed their contribution to the mutation burden of the tumors exposed to them (Figs. 3a, S8a and e). This contribution can be estimated through the activity of treatment-associated signatures recovered from the extraction with reasonable reliability (Supplementary Note 2). Next, adding the mutations contributed by different treatments to the same tumors we obtain the overall contribution of chemotherapies to the mutational burden of cells. Treatments administered to patients contribute between a few dozen and more than 10,000 SBS in tumors originated from different organs (Figs. 3b and c, S8b, c, f and g, Table S2). These contributions account for between 1% and more than 65% of the total tumor mutation burden. The median number of mutations of the cisplatin-associated mutations in pediatric metastatic tumor samples in a separate cohort^28^ is similar to that observed in adult tumors. However the median proportion of chemotherapy mutations is higher due to the lower activity of other mutational processes in pediatric tumors (Fig. 3b). A few tens of DBS are contributed by treatment associated signatures, which represent up to virtually half of the DBS burden in metastatic colorectal tumors, but as few as 13% in metastatic lung tumors, where tobacco carcinogens also contribute to the DBS burden. Platinum-based drugs contribute slightly more to the mutation burden of tumors than the aging signature, while capecitabine contributes slightly less (Figs. 4a, S8d, h). Nevertheless, while tumors are exposed to treatments during a comparatively short period of time, they are exposed to aging mutations during the entire lifespan of patients. Every month of the period of exposure (see definition above) to chemotherapies contributes two orders of magnitude more mutations to the burden of metastatic tumors than does exposure to the aging signature (Figs. 4a, S8d, h). Taking as reference a time closer to the guidelines of administration of chemotherapies (Table S4) instead of the annotated period of exposure of tumors, the number of chemotherapy-contributed mutations is of the same order (Figure S9).

**Figure 3.**
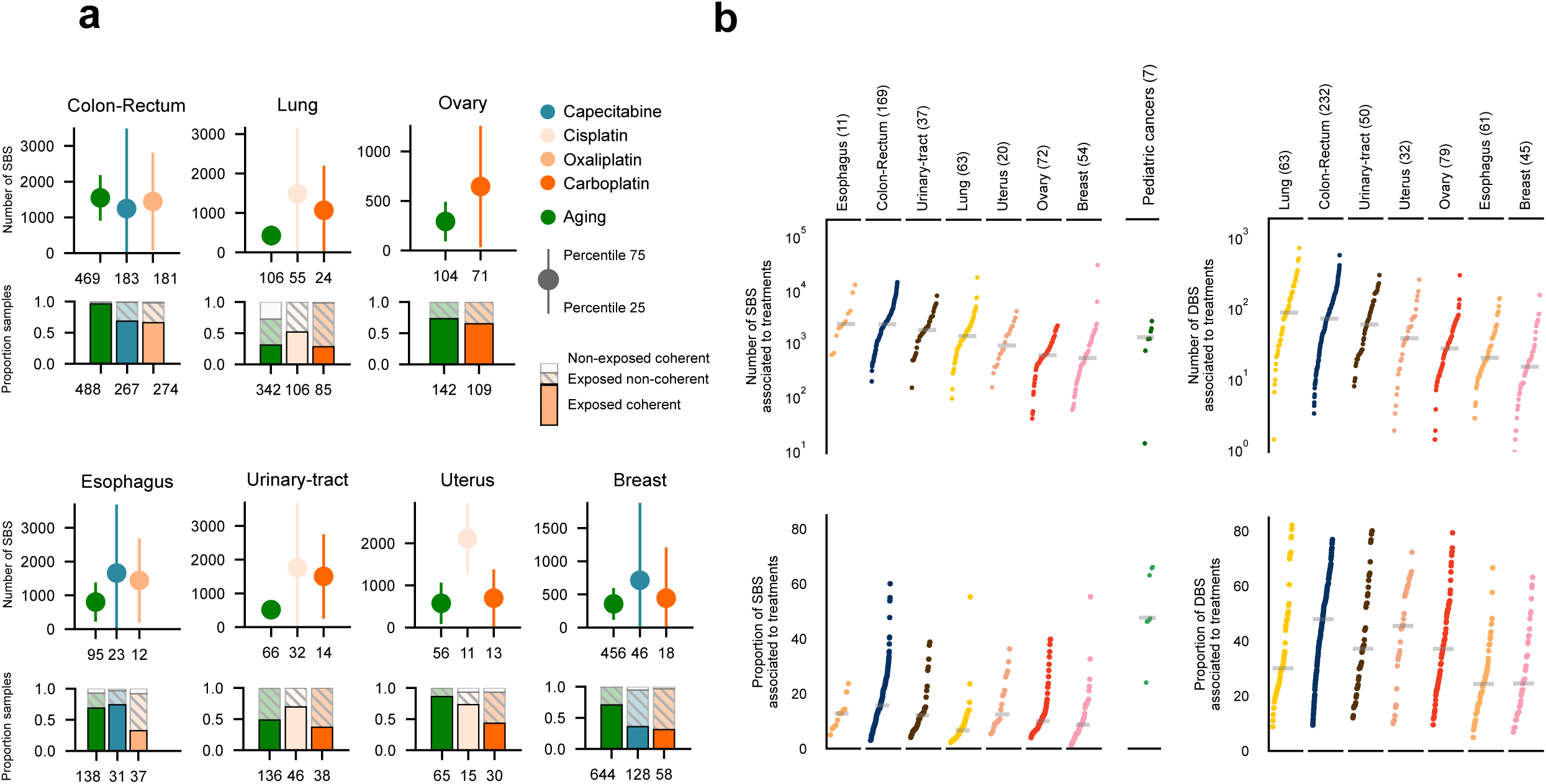
The contribution of anti-cancer treatments to the mutation burden of tumors. (a) Comparison of the contribution of different treatments and the aging signature to the mutation burden of tumors originated in different organs. Only tumors in which the activity of signatures according to SignatureAnalyzer and SigProfiler is coherent (less than 10% difference) are included in the contribution plots (coherent). Numbers in the x-axis represent the number of coherent tumors included. The contribution of signatures to the burden of coherent tumors is represented as a filled circle centered at the median of the distribution, and whiskers extending to its 25th and 75th percentiles. In the stacked barplots below each graph, the fraction of all tumors exposed to the treatment that are coherent are colored, and the numbers below each stacked bar correspond to the total of tumors with activity of the signature according to either method. Equivalent graphs with SignatureAnalyzer-extracted and SigProfiler-extracted signatures of all tumors are included in Figure S8. (b) Contribution in total number (upper) and proportion (lower) of all treatment-associated SBS (left) and DBS (right) to the mutation burden of metastatic tumors originated in different organs in the metastatic tumors cohort. A separate column in the left graph presents the activity of cisplatin-associated signatures (corresponding to the SBS Carboplatin/Cisplatin and SBS Cisplatin/Oxaliplatin obtained from the metastatic adults cohort) for cisplatin treated pediatric patients. Only coherent tumors are included in these plots (with numbers in parentheses).

**Figure 4.**
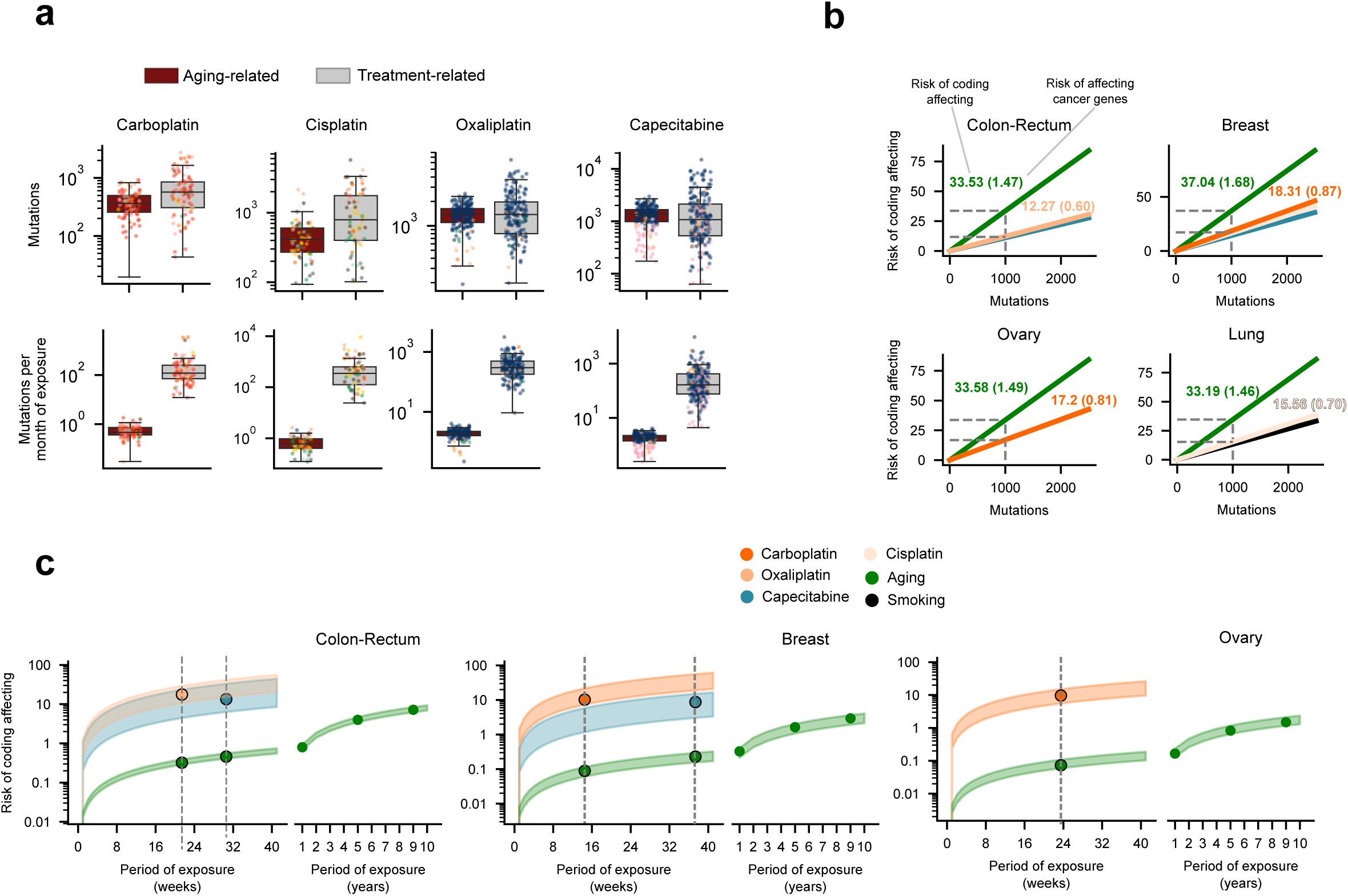
The mutational risk of anti-cancer treatments. (a) Contribution of treatment-associated signatures and aging signature to the mutational burden of metastatic tumors. First row: contribution of treatments (gray boxplots) and the aging signature (brown boxplots). Second row: distribution of the mutations contributed by treatments and the aging process to the mutation burden of tumors during one month of exposure to each. Each tumor is represented as a dot colored following the code of organ-of-origin presented in Figure 1b. The same analysis using as reference an estimate of the mode duration of the period of treatment with these drugs according to the clinical guidelines is presented in Figure S9. (b) Number of coding-affecting mutations expected to be contributed by several signatures (risk of coding affecting) estimated from their contribution to the mutation burden of tumors (Methods). The lines in the graphs corresponding to tumors originated in different organs represent the computed linear relationship between the total contribution of signatures and their risk of coding affecting. Broken lines mark the risk of coding affecting (spelled-out by numbers above the lines) that corresponds to a total contribution of 1000 mutations by the aging and treatment-associated signatures. In parentheses, number of known cancer genes sequence-affecting mutations expected to be contributed by several signatures (risk of affecting cancer genes) given their contribution to the mutation burden of tumors. (c) Risk of coding affecting mutations across tumors contributed by different signatures according to the duration of the exposure of tumors. Risk values are represented as a range spanning between the 25th and the 75th percentile of the distribution of contribution of signatures to the burden of tumors in one month of exposure (Fig. 3b). Vertical lines intersecting these risk value ranges are placed at the median of the distribution of times of exposure of all tumors of the given organ or origin to a given drug. The range of values of risk for the mutations contributed by the aging signature is extended several years to the right of the graph. The same analysis using as reference an estimate of the mode duration of the period of treatment with these drugs according to the clinical guidelines is presented in Figure S9. Details of the results shown in this figure are provided in Table S3.

### Chemotherapy treatments pose different risks of causing coding mutations

Another way to measure the toxicity of mutational processes is through their risk of causing coding mutations --or specifically mutations affecting cancer genes-- which could lead to malfunctioning cells or increase their likelihood to become malignant. Different mutational processes pose different risk of contributing coding mutations. The sequence determinants of the DNA damage underlying the signature (with respect to the nucleotide composition of coding sequences) and the mechanisms of DNA repair that correct it influence this risk. We reasoned that using the contribution of different therapies to the mutational burden of tumors we would be able to estimate their risk of introducing coding mutations (and mutations in cancer genes) in a patient’s cells. We devised a method to compute the expected load of coding-affecting mutations in one cell of the patient’s body (under neutrality) –across either all coding genes, or a set of known cancer genes^29^– contributed by a signature, given its overall contribution to the tumor mutation burden. The activities of the aging and treatment-associated signatures across the human genome are used to compute the probability of appearance of signature-associated coding mutations accounting for the mutational profile and the differential rate along the genome of each signature (Methods). We calculated that 33.53 of every 1000 mutations contributed by the aging signature across tumors of colorectal origin –given their tri-nucleotide preferences and distribution along the genome– affect the sequence of coding genes. More specifically, 1.47 mutations in a thousand are expected to affect the sequence of known cancer genes (Fig. 4b, S9). On the other hand, out of 1000 oxaliplatin-contributed mutations, only 12.27 are expected to affect the sequence of coding genes, and 0.60 that of known cancer genes. Finally, we computed the number of expected (or risk) coding-affecting that a patient receives in the real-life setting of chemotherapy treatments (Fig. 4c, S9). Tumors originated in the colon or rectum exposed during an overall period of 21 weeks to oxaliplatin (the median duration of the period of exposure observed for patients suffering from tumors with this origin, marked by the vertical broken line in the plots), receive some 20 coding-affecting mutations and one mutation affecting a cancer gene (Fig. S9). However, during the same period, less than one coding-affecting mutation and less than 0.01 mutations affecting cancer genes are contributed by the aging process. Across tumors of different organs of origin, the risk of appearance of coding-affecting mutations during a period of 21 weeks of exposure is more than ten times higher than that posed by the contribution of the aging signature (Fig. 4c, Table S4 and Fig. S9).

## Discussion

The short-term side-effects of some chemotherapies are mediated by the death of healthy cells triggered by toxic levels of damage to their DNA^30–34^. While this loss of healthy cells may also underlie some of their long-term side-effects, somatic mutations that result from this DNA damage across tissues probably also contributes to some of them, such as the emergence of secondary malignancies^35–37^. This is important to cancer survivors --children, in particular-- who could develop these long-term effects, such as secondary neoplasms even decades after their initial diagnosis and treatment.

Here, we estimated the mutational toxicity of three platinum-based drugs and capecitabine exploiting their identified mutational footprint across metastatic tumors. Most of these mutational footprints identified across tumors exposed to the drug have been validated by other studies^2,3,7,12–14^ or by us (in the case of capecitabine, despite an unsuccessful previous attempt to experimentally determine its footprint^38^). We use this mutational toxicity identified from samples of tumors exposed to these drugs as a proxy of their potential mutational effect across the patients’ healthy tissues. The availability of biopsies from patient’s metastasis together with the clonal expansion characteristic of tumor development provides a unique opportunity to identify drug-associated mutations (see Introduction and Fig. 1a). Although mutations would also accumulate in cells of healthy tissues, samples from these are harder to obtain and the lack of clonal expansion would render treatment-associated mutations much more difficult to detect.

The mutational risk computed here may thus be regarded as a bulk estimate of the mutagenic potential of chemotherapies across healthy cells. The mutational risk that chemotherapies pose for various types of healthy cells from different tissues may differ due to differences in the rate of division, hierarchy and proficiency of DNA repair. These reasons, and others, such as the pharmacodynamics and metabolization of drugs will likely also determine that they pose different risks between different tissues and individuals. This estimation will need to be refined through carefully planned prospective studies that periodically sample healthy cells (e.g. blood) from treated patients and survivors to monitor across the years the load of mutations introduced by chemotherapies.

Our estimate of the contribution of chemotherapies to the mutational burden of metastatic tumors per time of exposure are conditioned by the annotations collected regarding the duration of the period of exposure to each treatment. Since inaccuracies and omissions may appear amongst such annotations, we also made these calculations with average time of chemotherapy exposure taken from clinical guidelines, confirming overall similar mutation burden and toxicity. In any case, our estimate focuses on the order of magnitude --and it is meant to be understood as such-- of this contribution rather than on the actual number computed.

Although the tumors in the cohort were exposed to 206 different therapies (in complex treatment regimens), we only identified the mutational signatures of six widely-used treatments. On the one hand, therapies that don’t directly damage the DNA or alter the pool of nucleotides are not expected to leave a mutational footprint. On the other, in our analysis, we choose to be conservative, and true associations may lie under the stringent limit of significance set (Table S2). It is also possible that some associations are currently under the statistical power of this cohort. The approach developed here could be used to unravel novel drug-associated mutational signatures in larger cohorts or cohorts of specific treatments as they become available in the future.

In summary, in this study we present new mutational signatures associated to platinum-based treatments, confirm the role of defective DNA-repair pathways in certain treatment-associated signatures, and we discover the mutational footprint of capecitabine/5-FU. We use the contribution of treatment footprints to the mutational burden of tumors as a proxy of their contribution to mutations generated in healthy cells of patients undergoing chemotherapy. This study provides, for the first time, a window into the precise appraisal of the risk posed by chemotherapies to induce mutations in patients’ tissues –their mutational toxicity–, which may cause late side-effects, with special relevance for pediatric cancer survivors.

## Author contributions

O.P., A.G.-P. and N.L.-B. designed the project. O.P. carried out the analyses and built the figures. F.M. conceived the methodology to analyze the treatment-signature associations and the mutational risk, and carried out the simulation analysis in Supplementary Note 2. O.P. and F.M. developed and tested the framework to analyze the treatment-signature associations and mutational risk. O.P., F.M., A.G.-P. and N.L.-B. participated in the design of analyses and in the interpretation of the results. A.G.-P. and N.L.-B. drafted the manuscript. O.P., F.M., A.G.-P. And N.L.-B. edited the manuscript. A.G.-P. and N.L.-B. supervised the project. M.-P. L and M. S. contributed more than 5% of the samples in the adult metastatic dataset used in the analyses and provided feedback.

## Acknowledgments

N.L-B. acknowledges funding from the European Research Council (consolidator grant 682398) and Spanish Ministry of Economy and Competitiveness (SAF2015-66084-R, MINECO/FEDER, UE). IRB Barcelona is a recipient of a Severo Ochoa Centre of Excellence Award from the Spanish Ministry of Economy and Competitiveness (MINECO; Government of Spain) and is supported by CERCA (Generalitat de Catalunya). O.P. is the recipient of a BIST PhD fellowship supported by the Secretariat for Universities and Research of the Ministry of Business and Knowledge of the Government of Catalonia, and the Barcelona Institute of Science and Technology (BIST). A.G-P. is supported by a Ramón y Cajal contract (RYC-2013-14554). We acknowledge Santi Gonzalez for guidance in the analysis of mutations timing. This publication and the underlying study have been made possible partly on the basis of the data that Hartwig Medical Foundation has made available to the study. In particular, we want to acknowledge Neeltje Steeghs (NKI-AVL, Amsterdam), Martijn Lolkema (Erasmus MC, Rotterdam), Els Witteveen (UMC Utrecht, Utrecht), Haiko Bloemendal (Meander Medisch Centrum, Amersfoort), Henk Verheul (VUmc, Amsterdam), and Laurens V. Beerepoot, MD PhD (Elisabeth Tweesteden Ziekenhuis, Tilburg, the Netherlands), whose institutions contributed more than 5% of the samples in the adult metastatic dataset used in the analyses. Data from the Childhood Solid Tumor Network has also been used in the paper.

## Methods

### Genomics and clinical data of tumor samples

Single base substitutions (SBS), doublet base substitutions (DBS) and indels (ID), referred to collectively as mutations, detected in 3506 tumor samples (including relapses) were obtained from Hartwig Medical Foundation^15^; we call this the pan-metastatic adult cohort. We kept only mutations labeled as PASS by the calling pipeline and filtered out mutations in lowly mappable (Duke regions and CRG36mer) and fragile and low-complexity regions of the genome^39^. In parallel, clinical data of the donors of each sample was obtained from the same source. This data comprised the treatments administered to each patient in this cohort, and the date of beginning and end of each treatment round. We then converted treatment regimen acronyms to their unitary drugs and manually assigned drugs administered to patients to 58 different FDA drug categories (https://www.accessdata.fda.gov/cder/ndctext.zip), and the dates of beginning and end of treatments were used to compute the time of treatment.

The SBS of primary, metastatic and relapse tumor samples from 652 pediatric patients were obtained from the St. Jude Cloud (St. Jude cohort). Information regarding the treatment and duration of 12 samples taken from 4 of these patients with cisplatin was retrieved from the metadata of a related publication^28^. The exonic SBS and clinical data of one cohort of glioblastomas (treated with TMZ), as well as annotations of the tumors that had undergone hypermethylation of the MGMT promoter were obtained from a previous publication^13^. The SBS were processed as described for the pan-metastatic adult cohort cohort. Further details are presented in Supplementary Information.

### Extraction of mutational signatures active across tumor samples

The extraction of the mutational signatures active in the metastatic adult cohort tumor samples and the St. Jude pediatric cohort was carried out with the SignatureAnalyzer^16,17^ and the SigProfiler^2,18^. This decision was based on challenges observed by their authors in their effort to produce the catalog of mutational signatures in human cancers^3^ and to ensure that the conclusions obtained by this study do not depend on the signature extraction method but are robust to it. Briefly, to run the SignatureAnalyzer, we used the R implementation provided by the authors of the method (https://www.synapse.org/#!Synapse:syn11801488). Due to limitations in the obtention of a MATLAB license to run the signature extraction with the SigProfiler, we reimplemented this module in the Julia programming language and ran it in parallel in our computer cluster. We prepared the cohort of tumor samples for both methods as explained by their authors in the analysis of similar cohorts. All details on the execution of the methods and the comparison of their results are presented in Supplementary Information. We also extracted the signatures active across colorectal tumors using a non-NMF-based method^40^.

Throughout the paper, we present results based on analyses carried out using signatures and exposures obtained across samples using the SignatureAnalyzer. Equivalent results using the signatures extracted with the SigProfiler and their attribution to individual samples are presented as Supplementary Figures. To compute the number of mutations contributed by different signatures (presented in Figure 3 and 4) we selected tumor samples for which both methods differ no more than 15% in the number of mutations computed for treatment-associated signatures and the aging signature. Thus, the results of exposure and number of mutations contributed by each signature in the Figure constitute the mean of the values of both methods. In Supplementary Figures, the same results for all tumor samples based on each method are presented.

### Dependencies between individual treatments and signature exposures

To infer statistical dependencies between the treatments administered to the patients and the exposures to the mutational signatures uncovered, we required two levels of analysis. First, for each treatment label T we want to establish which signatures are strongly associated with T (step 1). Second, we must rule out spurious treatment-signature associations that could be explained with higher parsimony by another concomitantly administered treatment (step 2).

To address step 1, we devised a logistic regression approach with response variable Y representing whether T has been administered or not, and design matrix given by the relative exposures of each sample to each signature. Specifically, if N is the number of samples and s is the number of signatures, let X be the design matrix of size *N* × (*s* + 1) defined by the column vectors of normalized exposures (Z-scores) to each signature across all samples, also including an intercept column. We want to estimate β = (β_0_, β_1_, …, β_*s*_) such that *logit E*(*Y* … *X*) = *X* · β, i.e., the basal effect β_0_ (log-odds) and the log-odds ratios β_1_, …, β_*s*_.

A straightforward logistic regression approach would face an important challenge in our setting: the treatments being administered to the patients show dependencies on the tumor type and since the tumor type can also explain the exposure to tumor-type-specific signatures, tumor type is a clear confounder, hence we must correct for it. To this end, we fit an ensemble of logistic models to balanced, stratified random data samples. Specifically, we fit an ensemble of 1,000 L2-regularized logistic regression models with likelihood function of the form:

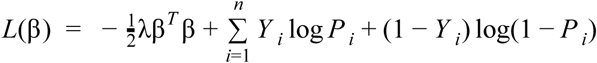

with 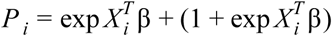 and regularization strength λ = 10.

Each logistic model was fitted with a randomized subset, balanced and stratified by tumor-type, i.e., for each tumor-type the same number of treated and untreated samples are drawn. Thus, we required the same number *n* = α · *min*(*t, u*) of treated and untreated samples to be drawn, where t (resp. u) are the number of treated (resp. untreated) samples for the tumor-type. The factor α was set to 1/3 as a compromise to prevent the same sample subgroups showing up in every randomization, while keeping each regression informative.

For each treatment and signature we obtained a vector (β_1_, …, β_*s*_) arising from each randomization that allowed us to compute an empirical p-value for each signature as the proportion of instances where the values are < 0 over the 1,000 randomizations. We also assessed the effect-size of each treatment-signature association as the average fold change of the exposures to the signature between treated and untreated samples. Finally, we deemed significant those treatment-signature associations with effect-size > 2 and p-value < 0.001.

In step 2 we aimed to assess the signature-specific mutation rate that can be allocated to each treatment when several concomitant treatments co-occur. The first step produced a collection of putative treatment-signature associations. However, we reasoned that some of these associations might be artifacts explained by the fact that several treatments are administered to similar sets of patients, in such a way that some treatment could “borrow” the association from the true causal treatment.

Given a treatment T and a signature S, we were bound to estimate the relative contribution of T to the exposure of S compared to other concomitant treatments associated with S. To this end we conducted a positive least-squares regression, as follows: let N be the number of samples, let X be the *N* × 2 design matrix with binary values with columns corresponding to T and a concomitant treatment C, and let Y be the N-dimensional vector of exposures of the target signature S. We want to estimate β = (β_*T*_, β_*C*_) with β_*i*_ ≥ 0 such that *E*(*Y* |*X*) = *X* · β. We can think of each β_*I*_ as an “average efficiency” to generate exposure of signature S; likewise, we can think of β_*T*_ /β_*C*_ as the “relative efficiency” of T with respect to C. Bearing in mind this set-up, we can now analyze all the concomitant treatments of T and check in each case whether the estimated efficiencies support that T is the most efficient generator of exposure of signature S: if the resulting efficiency of T is higher than all the other concomitant treatments associated to S, we conclude that T is the treatment most likely associated with S.

Finally, we run the above described steps with two treatment settings: coarse-grained and fine-grained. The coarse-grained setting considers groups of treatments by FDA category. The fine-grained setting considers specific treatment labels. For the sake of consistency, we deem a treatment-signature association significant if either of the following conditions hold: i) both the specific treatment and its FDA group raise significance in the fine-grained and coarse-grained setting, respectively; ii) the specific treatment raises significance in the fine-grained setting, but no FDA group raises any significance in the coarse-grained setting.

### Validation of the approach using synthetic datasets

We built synthetic datasets of mutations that are similar to the metastatic tumors analyzed with regard to the composition of mutational signatures. We then injected a known number of mutations drawn from the mutational profile of a foreign signature to a known number of samples of these synthetic datasets. We thus control the number of samples bearing the mutational footprint of the drug, the number of drug-induced mutations present in each sample, the signature of the drug-induced mutations and the number of samples known to have undergone treatment (allowing for discrepancies between these two parameters). Using these synthetic datasets, we tested i) the extraction of drug-associated signatures, ii) the detection of the mutational footprints of drugs through the regression ensemble, iii) the identification of the correct etiology of the signature in the case of tumors exposed to co-treatments, and iv) the accuracy of the estimation of the number of mutations contributed by drugs to the burden of tumors. In the analyses, we challenged our entire methodological setting with fluctuations in the synthetic data reflecting a variety of common scenarios. The analysis of these synthetic datasets demonstrates that the approach followed correctly identifies the foreign signatures as the molecular footprints of anti-cancer treatments within a wide range of numbers of exposed samples. The methodology is robust to systematic errors such as miss-annotation of treatments or lack of activity of the associated signatures in a subset of exposed samples. It is also able to estimate the mutational burden contributed by the treatment within acceptable confidence intervals. The results of these analyses have been useful to fine-tune the parameters of the methodologies developed to detect the mutational footprint of treatments. Details of the methodology and results of the analysis with synthetic datasets are in Supplementary Note 2.

### Identification of mutational signatures active across other metastatic tumors

Due to the low number of mutations in the glioblastoma cohort employed in the analyses, rather than extracting mutational signatures *de novo*, we fitted the catalog of identified mutational signatures^7^ to the mutational profile matrix of each sample of the cohort. We employed deconstructSigs^41^ using PCAWG SBS^3^ as a reference signatures.

### Strand asymmetry of treatment-associated signatures

To compute the signatures activity strand asymmetry we used a slight modification of an approach described elsewhere^42^. Briefly, using dypirimidines as a base reference, we classified each of the mutations as occurring in either transcribed and non-transcribed (or leading and lagging). We then retrieved the trinucleotide context, thus obtaining 96 channels for both transcribed and non-transcribed (leading and lagging) (192 in total). The identity of the signatures extracted across the 192 channels (averaged) is assessed through their cosine similarity to the signatures extracted from the adult metastatic cohort across the 96 channels. We pooled the tri-nucleotide counts corresponding to each of the six main base change channels (C>A through T>G) and selected the channel with the largest contribution to the signature profile to represent it. Then, the activity of these channels in the transcribed and non-transcribed (or leading and lagging) strands were computed. Letting the activity in the transcribed (or leading) strand be *S*_1_ and the activity in the non-transcribed (or lagging) strand be *S*_2_, we computed the asymmetry as |1 − 2*S*_1_/(*S*_1_ + *S*_2_) |. This way, a strand asymmetry of 1 means the activity of the signature is restricted to either the transcribed (or leading) or the non-transcribed (or lagging) strand. This is the value plotted in Figure 2b.

### Relationship between the activity of treatment-associated signatures and the duration of treatment

We sorted metastatic tumor samples originated from each organ following the duration of their exposure to different treatments. Then, for cohorts with more than 40 tumor samples with mutations associated with each treatment, we made two groups of samples, LT y ST containing the 25% tumor samples with longer and shorter treatment duration, respectively. We obtained the number of mutations associated to treatment i in a tumor as:

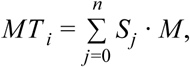

where *S*_*j*_ is the exposure of the tumor to one of the mutational signatures associated to treatment i and M is the total mutation burden of the tumor. Finally, we compared the distribution of the burden of treatment-associated mutations of ST and LT tumor samples using the Mann-Whitney U test.

### The timing and clonality of treatment associated mutations

We used the MutationTime. R package developed elsewhere^24^ and tested across 2658 primary tumor samples, which exploits large chromosomal amplifications and/or whole-genome duplication of a tumor, to classify all its SBS as early, late or subclonal. The method classifies mutations in a tumor as clonal early, clonal late, or subclonal. Then, we associated each mutation uniquely with a mutational signature using a maximum likelihood approach. We computed a fold-change between the relative proportions of late and early clonal mutations associated to platinum-based drugs and capecitabine/5-FU, as well as other selected mutational signatures. This provided the relative activity of each signature in early and late stages of tumor development. Similarly, we computed a fold-change between the relative proportions of clonal (grouping early and late clonal mutations) and subclonal mutations associated to platinum-based drugs and capecitabine/5-FU, as well as other selected mutational signatures. This provided the relative activity of each signature among clonal and subclonal tumor mutations.

### Risk of acquiring coding-affecting mutations through treatments

For each cohort of tumor samples we inferred the proportion of neutral mutations hitting coding non-synonymous sites that can be explained by a group of etiologies. The attribution of the observed mutations to etiologies was carried out resorting to the signatures for which we could establish an association with the etiology. The etiologies –alongside their corresponding SigProfiler signatures– are the following:

**capecitabine:** E-SBS19;

**carboplatin:** E-SBS1;

**cisplatin:** E-SBS1;

**oxaliplatin:** E-SBS20;

**tobacco-smoking:** E-SBS17;

**aging:** E-SBS23;

To conduct this analysis, we partitioned the sequence of the human genome into 1Mb chunks. Non-mappable and repetitive positions were discarded. For the etiology and cohort of samples of interest, we considered all the mutations observed in each chunk, excluding those mutations in Cancer Gene Census (CGC) genes^29^ to avoid positive selection bias.

To model the local mutation rate explained by an etiology S across 1Mb chunks, we rely on a generative probabilistic model whereby: i) the probability that a new mutation occurs in a 1Mb chunk is proportional to the average number of mutations in this chunk explained by S across samples; ii) the probability that a new mutation reaches a specific site in the 1Mb chunk is proportional to the normalized relative frequency (i.e., assuming same abundance for all trinucleotides) of its context according to the signature S.

From the signature deconstruction analysis we inferred the function *P* (*c, i*) encoding the conditional probability that a mutation in context c and sample i has been generated by signature S. Given a chunk, say k, let *n*_*ci*_ be the number of mutations in context c and sample i in this chunk. Then the average number of mutations explained by signature S across samples in chunk k is:

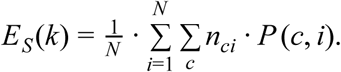

If *f*_*c*_ stands for the normalized relative frequency for channel c in signature, we assigned all the per-mappable-site mutation probabilities of the chunk as follows: letting *n*_*c*_ be the count of mappable sites in context c, then all the sites of the chunk in context c are given the same probability *p*_*c*_ determined by the following two conditions:

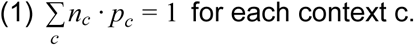

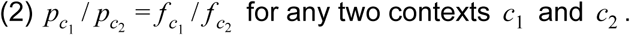

Finally, using VEP 88^32^ we annotated the consequence types for every genic (coding) mappable site of the chunk. We then counted all nucleotide changes yielding mutations that potentially affected the sequence of coding genes (i.e., non-synonymous and truncating) for every context c in the chunk: let *m*_*c*_ be this count for context c. Finally, the proportion of non-synonymous mutations among neutral mutations explained by S can be quantified as the following quotient:

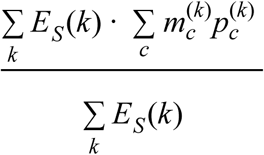

where we denote the specific counts and probabilities for each chunk with the (k) superscript.

In summary, we got a site-specific neutral mutation rate estimate by linearly spreading a unit of exposure, first by using the observed mutations to define local mutation rates in 1Mb chunks, then by spreading probabilities in accordance with the operative signature. And we used this neutral mutation rate model to derive an expected overlap of the unit exposure with the coding, non-synonymous region.

### 5-fluorouracil mutations in mutant strains of *Leishmania infantum*

Sequencing reads of five mutant strains of *Leishmania infantum* resistant to the treatment with 5-fluorouracil, and the parental sensitive strain^23^ were obtained from the EGA database (ERP001815 and ERP002415). The five mutant strains had been treated with 5-fluorouracil previous to sequencing, while the parental strain was cultivated under the same conditions (with exception to the drug) and for the same duration. We downloaded the *Leishmania infantum* reference genome from the Ensembl genomes database, and aligned the reads of both the resistants and the parental strains to its sequence, using bowtie2. In the resulting sam files, the aligned reads were sorted and processed with samtools, and mutations were called for the parental and resistant strains. High quality mutations (above 20) were used to build the mutational profile (tri-nucleotide context changes) of each sequenced strain.

### Significance of cosine similarity with respect to a signature

Given a mutational signature *S* (i.e. SBS capecitabine) and a cosine similarity *C* (i.e. 0.8) we can associate a p-value to *C* relative to the signature *S* by randomly drawing vectors σ from the signature simplex and computing the frequency with which cos(*S*, σ) ≥ *C*. We carried out this computation with a random generator that produces signatures with same expected sparsity as found in the COSMIC catalogue: signatures are chosen uniformly from COSMIC catalogue, then a random permutation is applied on the channels.

