## Supplementary Figures for "The mutational footprints of cancer therapies"

### Supplementary Figure 1

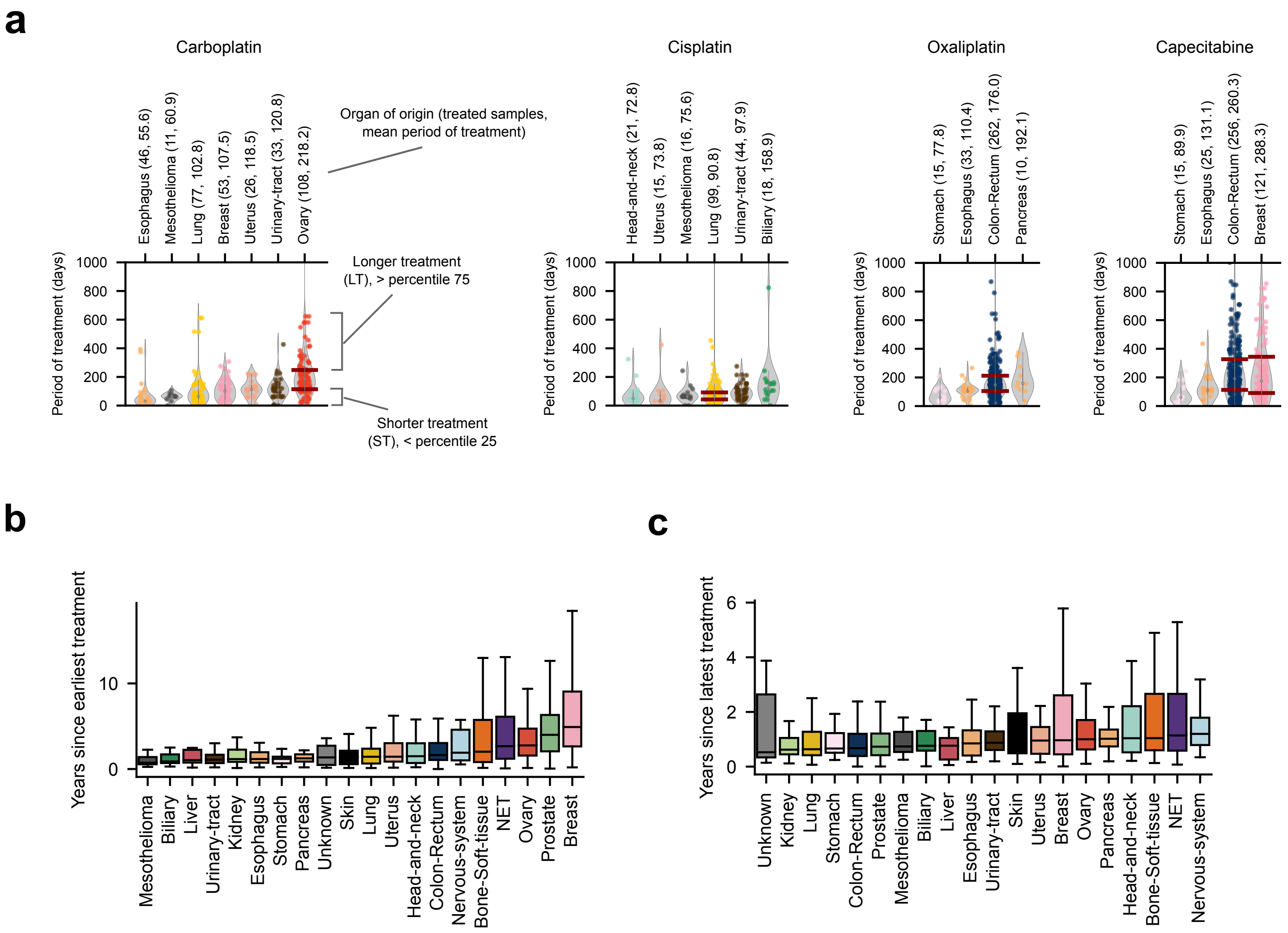

**Figure S1. Time between treatment and metastasis biopsy of metastatic adult tumors**

(a) Distribution of the time of treatment of patients (with tumors originated in organs with at least 10 samples in the cohort) with carboplatin, cisplatin, oxaliplatin and capecitabine. Red bars perpendicular to the violin-plots representing the distribution of tumors correspond to the upper boundary of their first quartile (short-treatment, or ST tumors) and the lower boundary of their fourth quartile (long-treatment, or LT tumors). This nomenclature is used in Figure 2.

(b) Distribution of time elapsed since earliest treatment administered to patients in the pan-metastatic adult cohort. Metastatic tumors are grouped according to the organ of origin of the primary.

(c) Distribution of time elapsed since latest treatment administered to patients in the pan-metastatic adult cohort. Metastatic tumors are grouped according to the organ of origin of the primary.

### Supplementary Figure 2

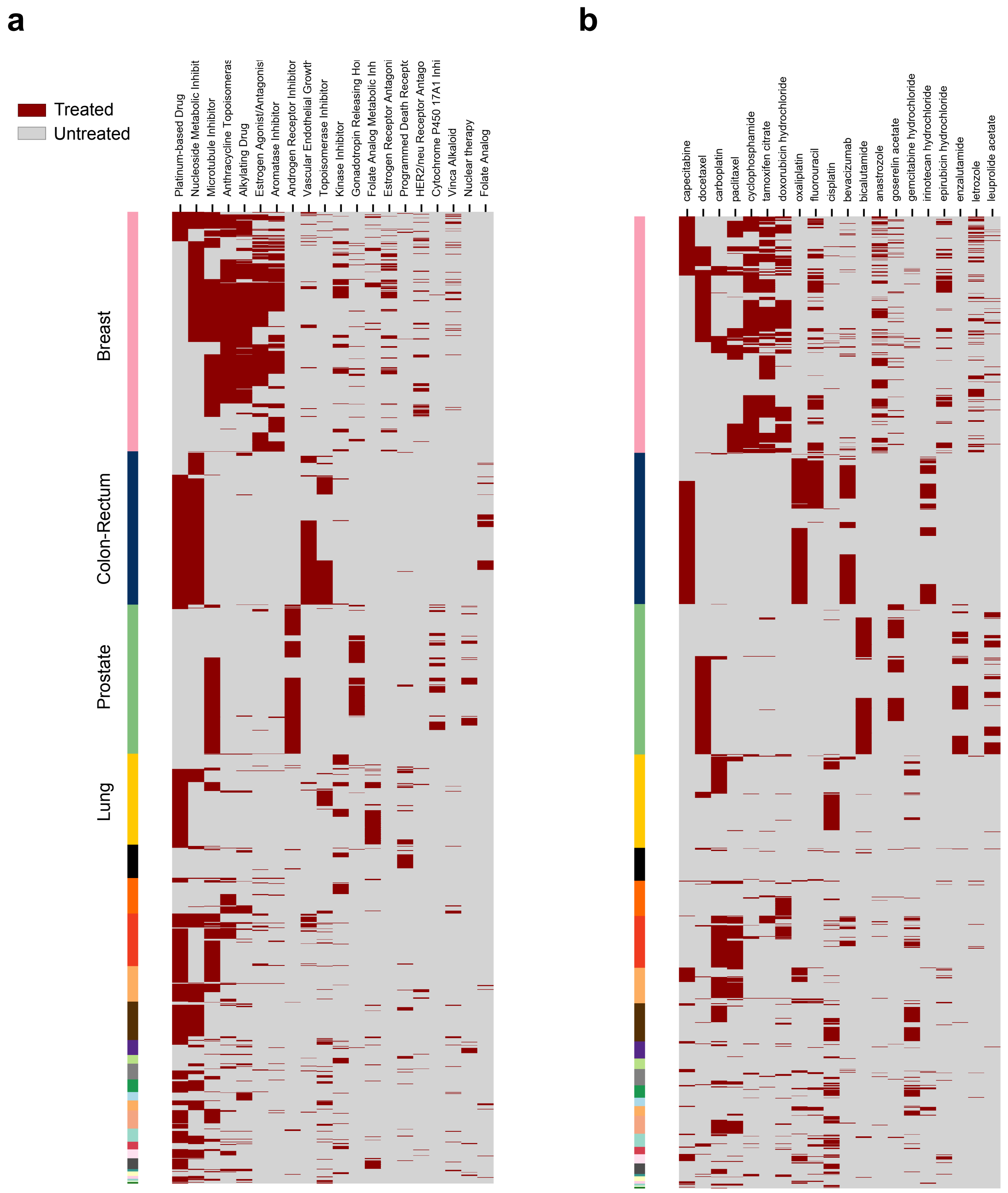

**Figure S2. Landscape of treatments administered to patients in the metastatic adult cohort**

(a) Heatmap representing the exposure of tumors originated in different organs (rows, represented by the color annotation by the left border that follows the code introduced in Fig. 1 of the main paper) to groups of drugs of different FDA clases (columns).

(b) Heatmap representing the exposure of tumors originated in different organs (rows) to selected chemotherapies.

### Supplementary Figure 3

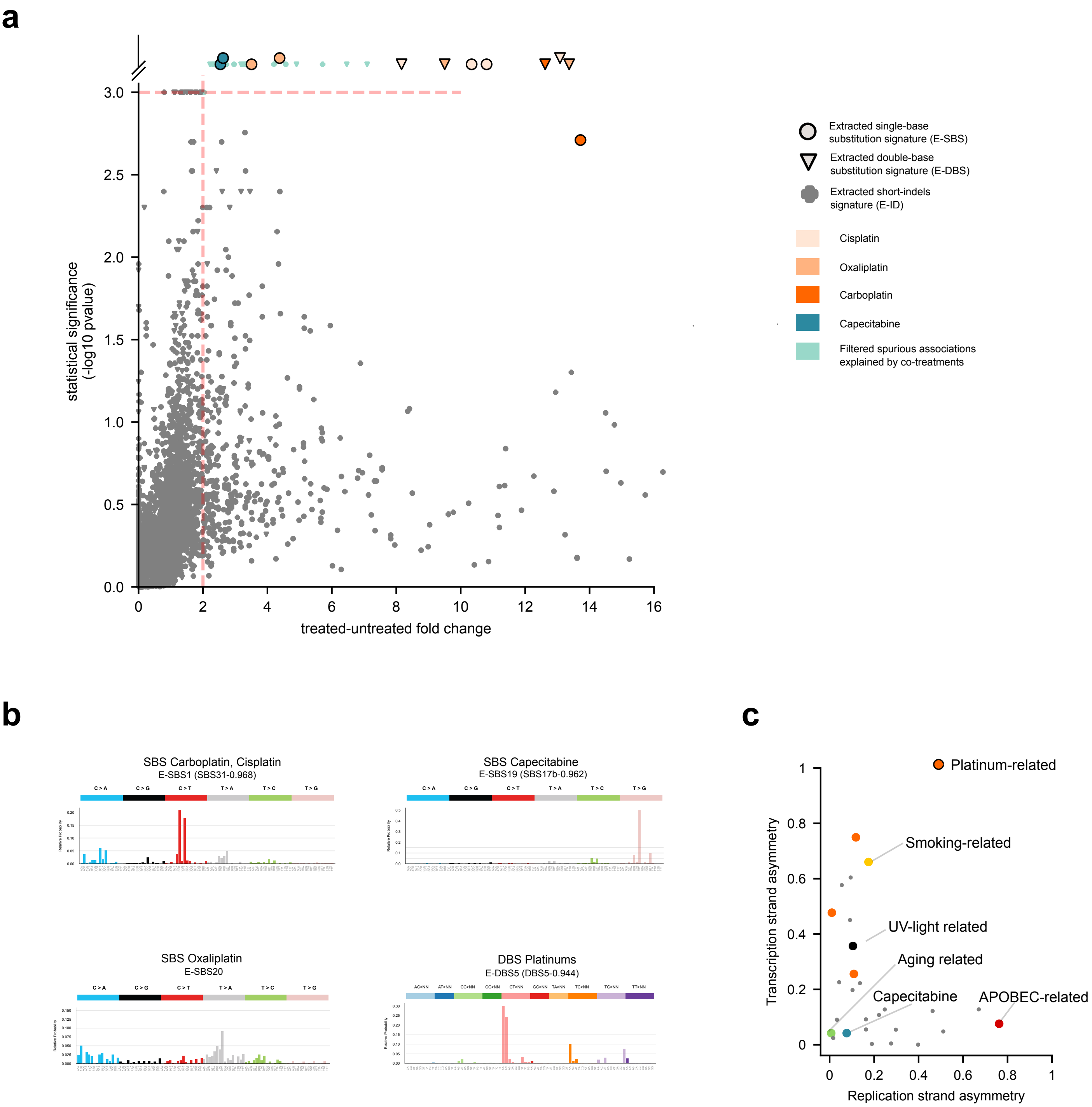

**Figure S3. Treatment-associated signatures extracted using SigProfiler**

(a) Treatment-associated mutational signatures (extracted with SigProfiler). Each dot represents the effect-size and the p-value of the association between one signature and one treatment detected through one regression model. Associations deemed significant (effect-size>2 and p-value<0.001) are colored according to the treatment and shaped following the type of variant (SBS, DBS, ID) of the mutational signature. The signature analogous to the SBS Carboplatin/Cisplatin extracted by SignatureAnalyzer (Fig. 2a) appears very close to significance (p-value=0.002; highlighted in the panel) and has thus been “rescued” as associated to the treatment in this figure. This panel is equivalent to Figure 1e of the main paper.

(b) Strand asymmetry of selected SigProfiler-extracted signatures active across healthy and tumoral somatic cells. Each dot corresponds to a signature associated to a therapy (colored following the code in Fig. 1e of the main paper), with the abscisa representing its replication strand bias (the unbalance of its mutations between leading and lagging strand), and its ordinate, the transcriptional strand bias (the unbalance of its mutations between template and non-template genic strands). Detailed results in Table S2. This panel is equivalent to Figure 2b of the main paper.

### Supplementary Figure 4

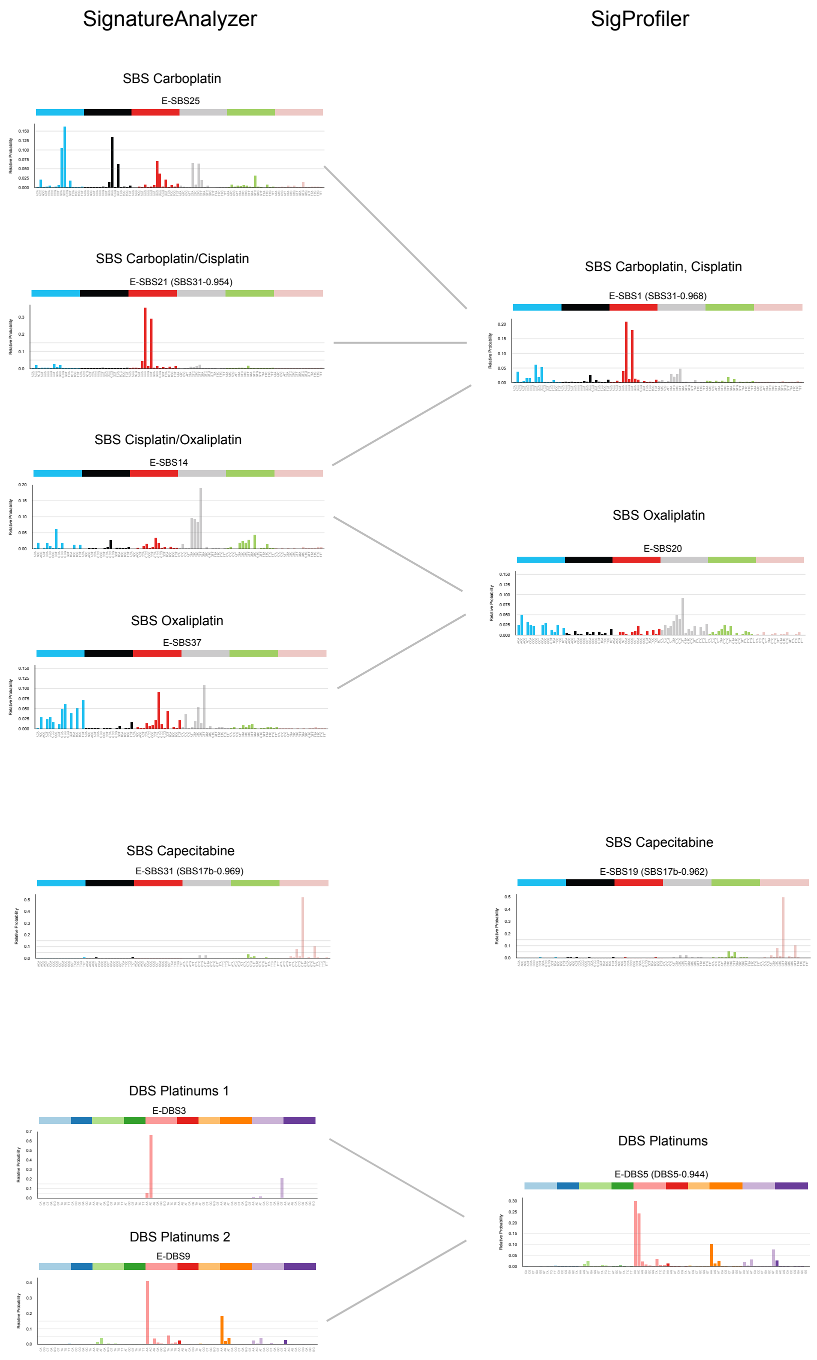

**Figure S4. Comparison of treatment-associated signatures extracted with SigProfiler and SignatureAnalyzer**

### Supplementary Figure 5

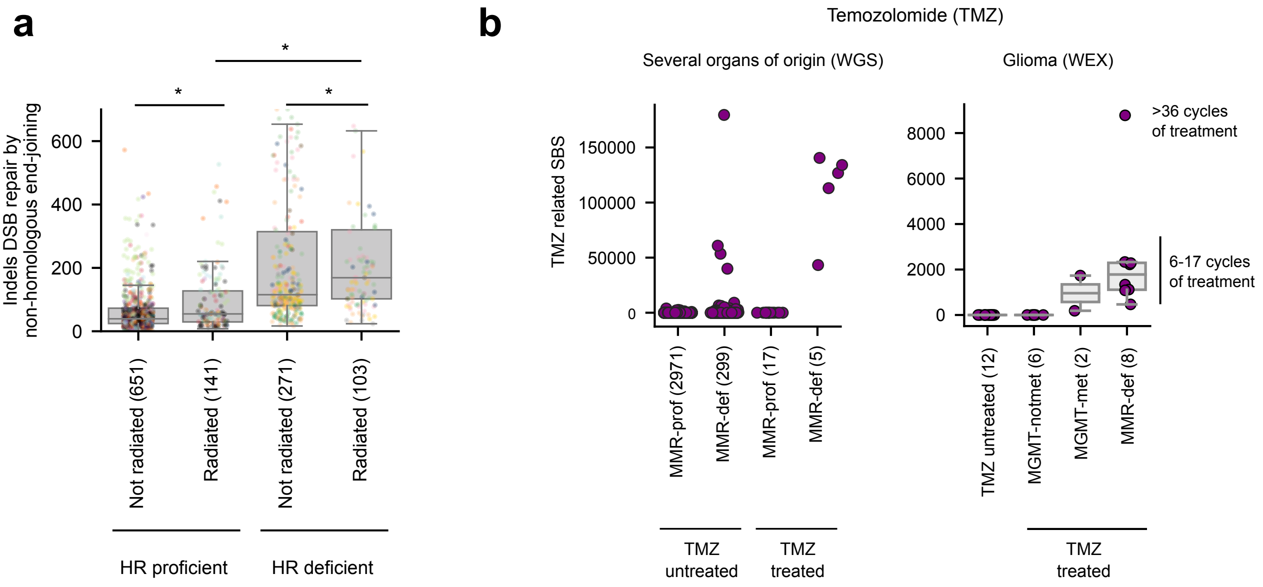

**Figure S5. Uncovering mutational signatures associated to radiation and temozolomide**

(a) HR-deficiency plays a key role in the appearance of an ID signature previously associated to radiation. Tumors in the fourth quartile of activity of HR signature (BRCAness signature) are considered HR-deficient, while tumors in the first quartile are considered HR-proficient. Boxplots compare the distribution of the number of IDs of this signature across HR-deficient and HR-proficient tumors either exposed or unexposed to radiation. SignatureAnalyzer-extracted signatures and their activities in tumors are represented in the figure.

(b) MMR or MGMT-deficiency plays a key role in the generation of a TMZ-associated SBS signature. Left panel represents the load of TMZ-associated SBS in tumors of the cohort exposed or unexposed to TMZ separated by their MMR status (considered defective if at least one protein-affecting mutation was detected among a list of MMR-related genes). Right panel is the same for TMZ-related exonic mutations across glioblastomas of an independent cohort.

### Supplementary Figure 6

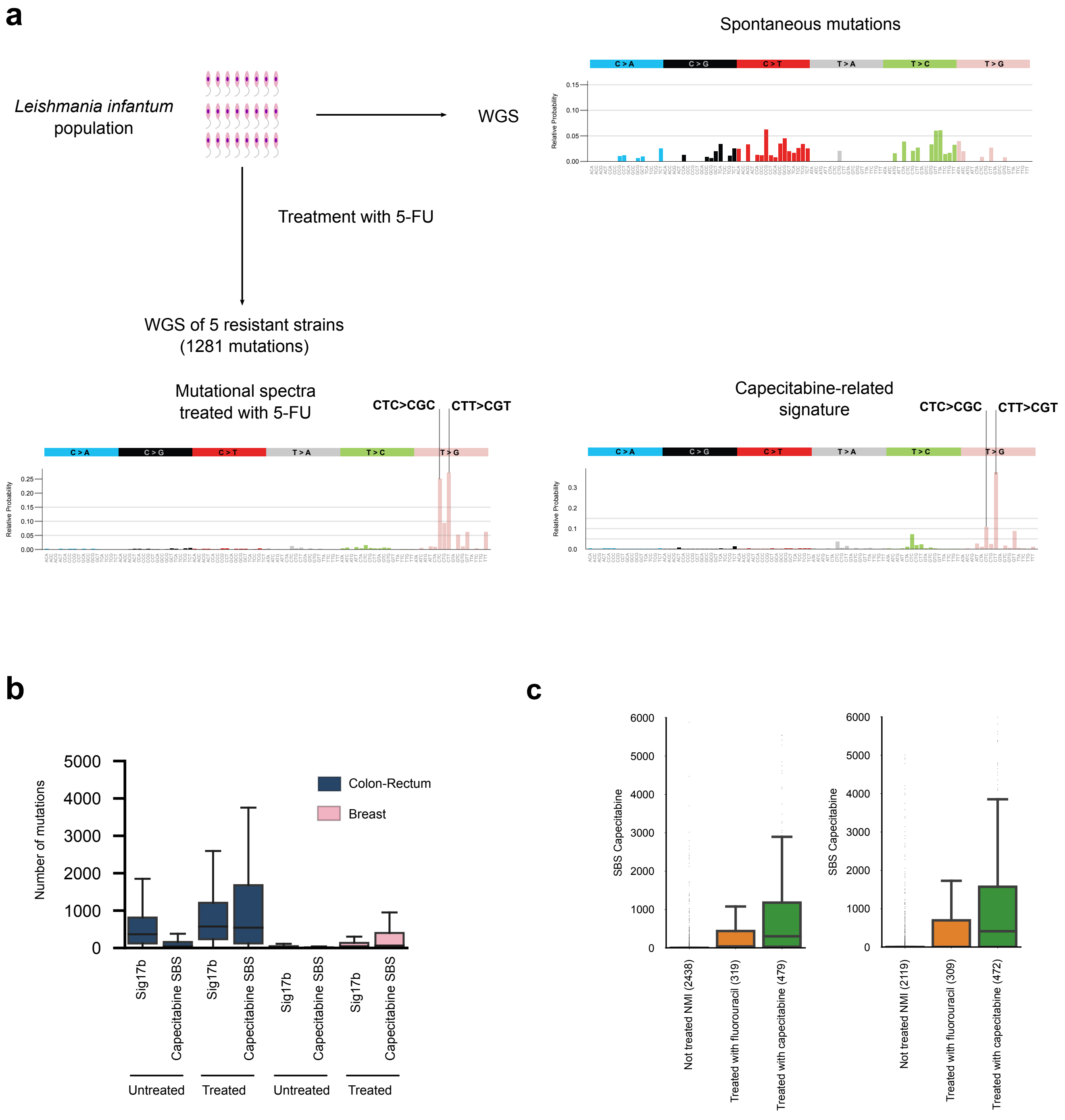

**Figure S6. The capecitabine/5-FU mutational footprint**

(a) Mutational profile of 5-FU-induced mutations in five resistant strains of *Leishmania infantum*. The profile was built with the mutations private to the strains (that is, after subtraction of the mutations found in the parental strain).

(b) Contribution of Capecitabine SBS (Capecitabine/5-FU SBS) and the previously reported 17b signature (Sig17b) to the mutation burden of tumors that have not been exposed (left boxplots) or exposed (right boxplots) to capecitabine/5-FU. The left graph corresponds to the activity of both signatures in colorectal tumors, and the right graph to their activities in breast tumors.

(c) Contribution of capecitabine (green) and fluorouracil (orange) to the mutation burden of tumors exposed to either drug, compared to tumors not treated with nucleoside metabolic inhibitors. The left graph corresponds to the Capecitabine SBS extracted with SignatureAnalyzer and the right graph to the Capecitabine SBS extracted using SigProfiler.

### Supplementary Figure 7

**a**

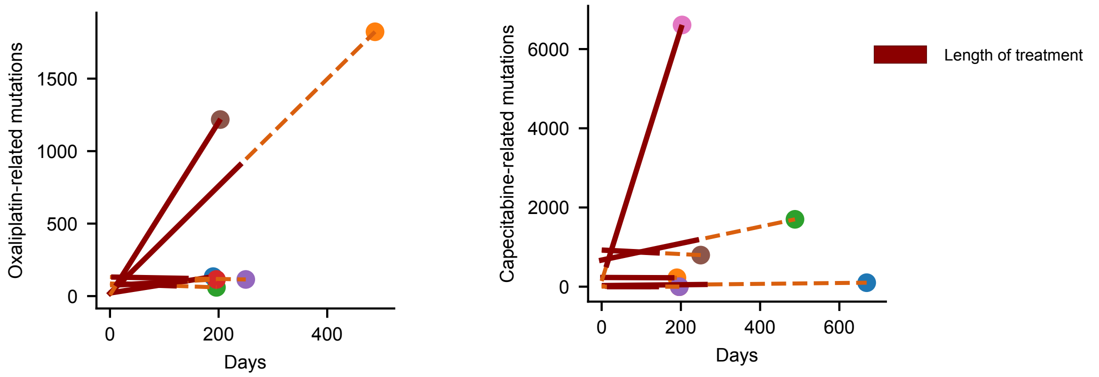


**b**

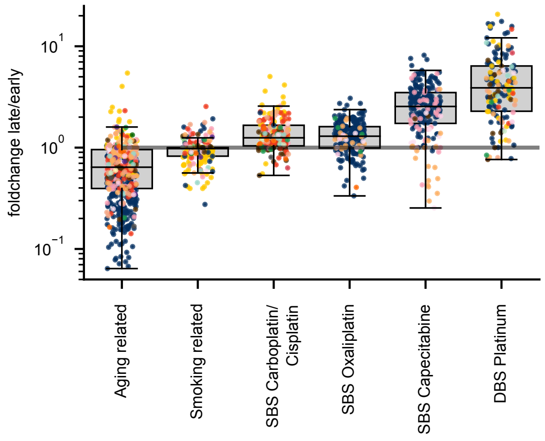


**c**

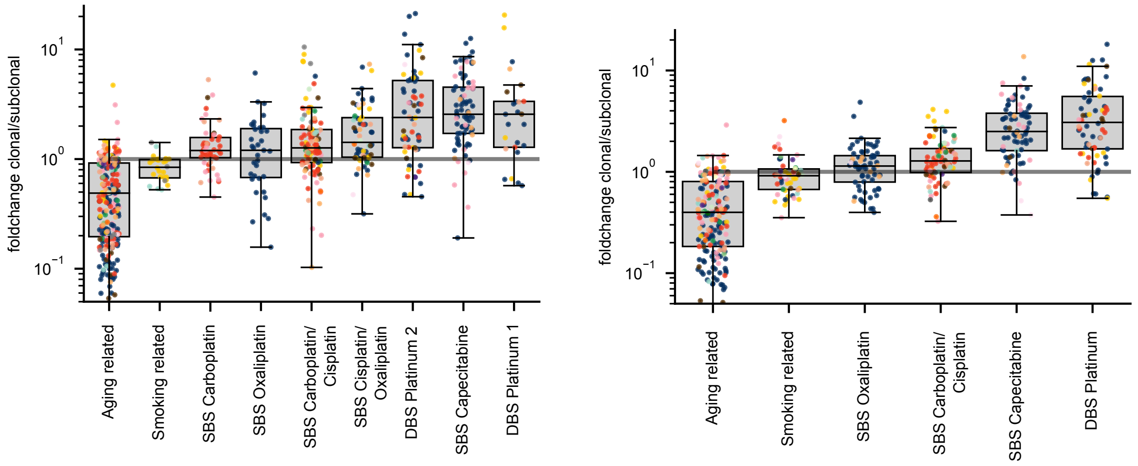

**d**

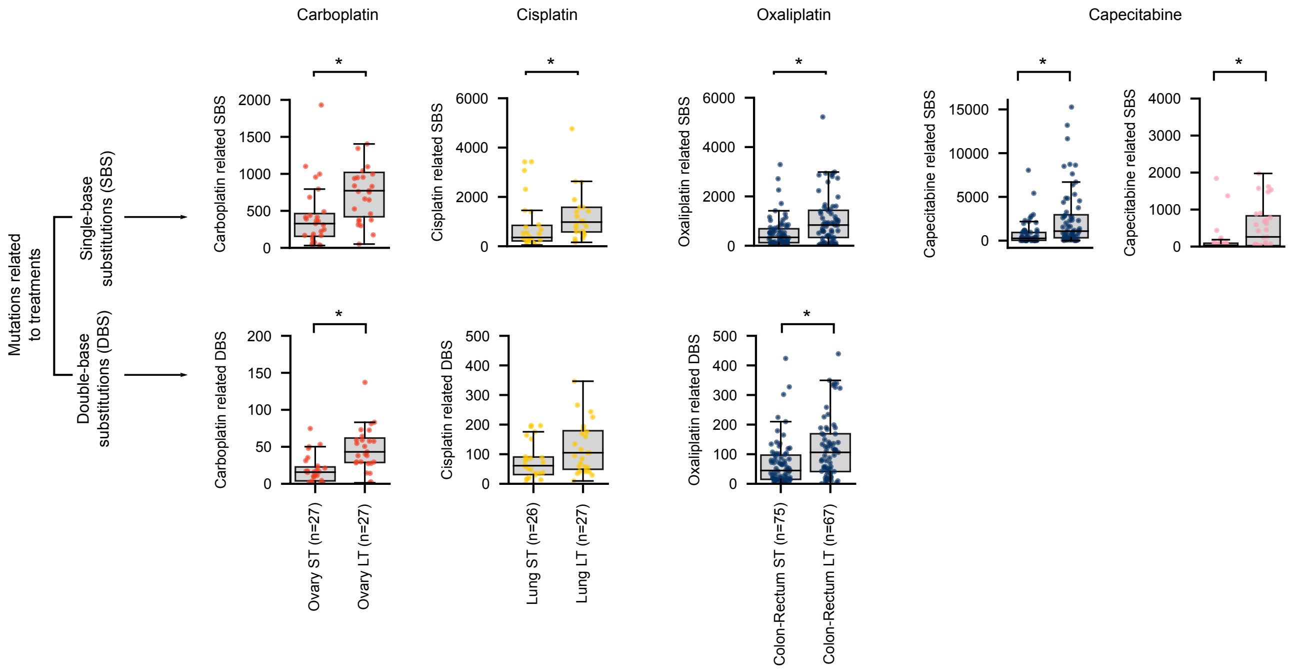
Carboplatin
Cisplatin
Oxaliplatin
Capecitabine
Mutations related to treatments
Single-base substitutions (SBS)
Double-base substitutions (DBS)
Carboplatin related SBS
Cisplatin related SBS
Oxaliplatin related SBS
Capecitabine related SBS
Carboplatin related DBS
Cisplatin related DBS
Oxaliplatin related DBS
Ovary ST (n=27)
Ovary LT (n=27)
Lung ST (n=26)
Lung LT (n=27)
Colon-Rectum ST (n=75)
Colon-Rectum LT (n=67)

**e**

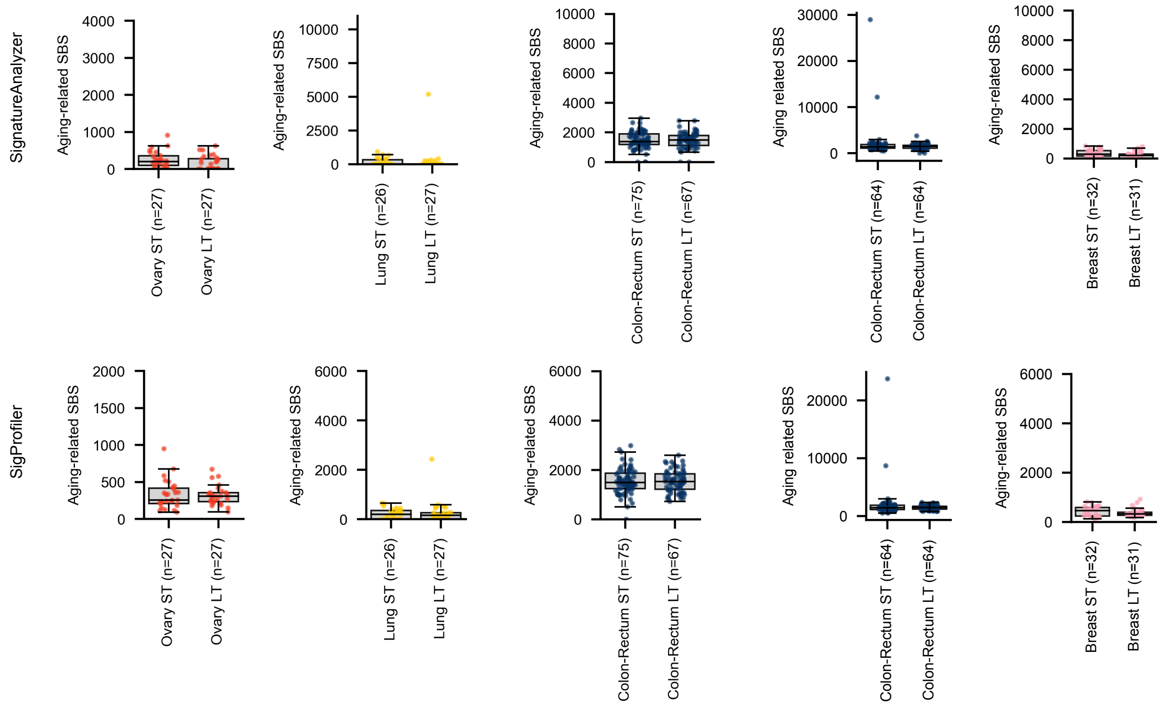
SignatureAnalyzer
SigProfiler
Aging-related SBS
Aging related SBS
Ovary ST (n=27)
Ovary LT (n=27)
Lung ST (n=26)
Lung LT (n=27)
Colon-Rectum ST (n=75)
Colon-Rectum LT (n=67)
Colon-Rectum ST (n=64)
Colon-Rectum LT (n=64)
Breast ST (n=32)
Breast LT (n=31)

**Figure S7. Treatment-associated mutations are recent and subclonal because they occur late in tumor development.**

(a) Treatment-associated mutational signatures appear active in post-treatment, but not pre-treatment samples. Pairs of samples corresponding to biopsies of the same patient taken before the start and during or after treatment are represented as pairs of dots connected by a dashed line in the figure. The first biopsy (before treatment) appears at the left and the second (during or after treatment), at the right. The vertical position of each dot represents the activity of the treatment-associated signatures (with the contribution of different signatures associated to the same therapy summed). The dots of the first biopsies are aligned (time 0), while the dots of the second biopsies are dispersed, so that the distance between the two is proportional to the time elapsed between both biopsies (x-axis). Solid segments in the lines connecting the dots that correspond to the biopsies of tumors from the same patients represent the duration of the treatment administered to each of them. The upward trajectory of the lines corresponding to patients treated longer (more likely to carry tumors with a detectable footprint) supports the conclusion that the signatures associated to treatments through the regression are indeed the mutational footprint of the therapies.

(b) SBS of signatures associated to treatments (following the nomenclature of Fig. 2a) are enriched for later substitutions (higher late-to-early fold-change), in comparison to signatures active since earlier in the lifetime of the patients (e.g., aging sig. 1 and tobacco-related sig. 4). SigProfiler-extracted signatures are represented in the Figure. This panel is equivalent to Figure 2e of the main paper.

(c) SBS of signatures associated to treatments (following the nomenclature of Fig. 2a) are enriched for subclonal substitutions in comparison to signatures active since earlier in the lifetime of the patients (e.g., aging signature and tobacco-related signature). The left graph corresponds to signatures extracted with SignatureAnalyzer and the right graph to signatures extracted using SigProfiler.

(d) The mutation load contributed by treatment-associated signatures correlates with the duration of the period of exposure to the treatment (SigProfiler-extracted signatures). The panels compare the distribution of the number of SBS (upper row) and DBS (lower row) of signatures associated to each drug across ST and LT tumors (see definition in main text) of organ of origin with sufficient mutations to carry out the comparison. In every case, LT tumors possess significantly more mutations than ST tumors. This panel is equivalent to the lower panels in Figure 2c of the main paper.

(e) The mutation load contributed by the aging signature does not correlate with the time of exposure to treatments. Upper panels correspond to signatures extracted with SignatureAnalyzer and lower panels to signatures extracted using SigProfiler.

### Supplementary Figure 8

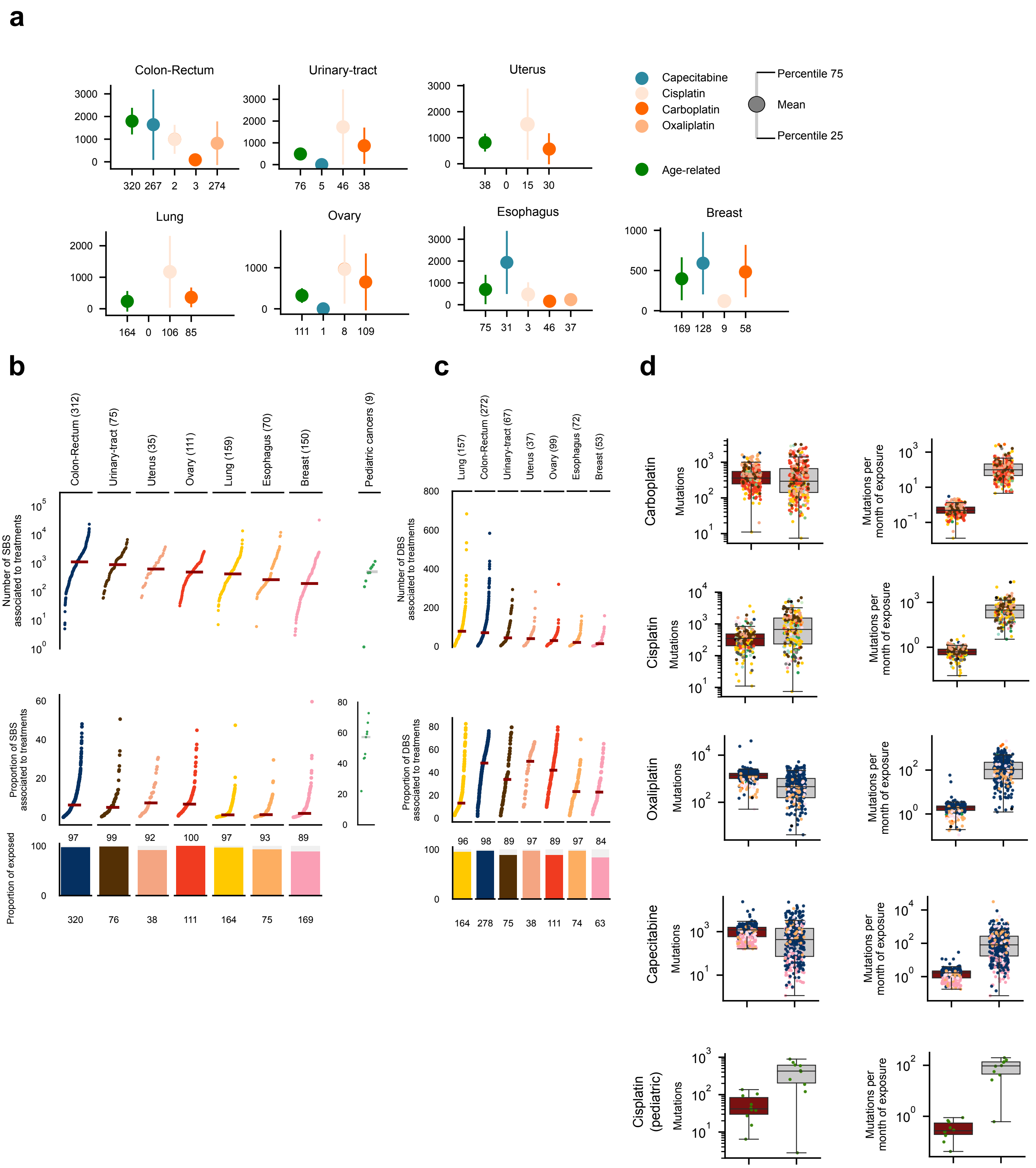

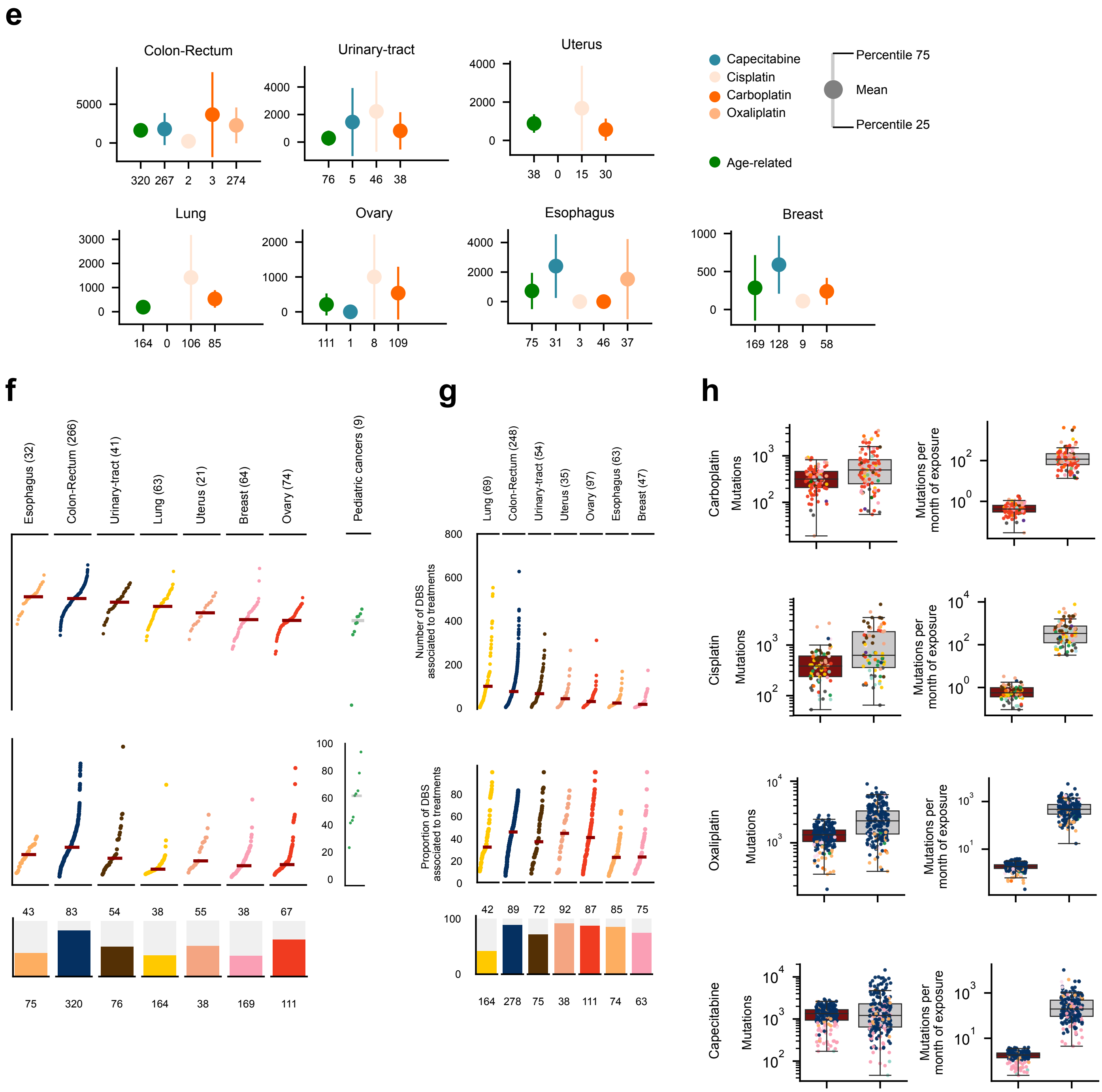
e
Colon-Rectum
Urinary-tract
Uterus
Lung
Ovary
Esophagus
Breast
Capecitabine
Cisplatin
Carboplatin
Oxaliplatin
Age-related
Percentile 75
Mean
Percentile 25
320 267 2 3 274
76 5 46 38
38 0 15 30
164 0 106 85
111 1 8 109
75 31 3 46 37
169 128 9 58
f
Esophagus (32)
Colon-Rectum (266)
Urinary-tract (41)
Lung (63)
Uterus (21)
Breast (64)
Ovary (74)
Pediatric cancers (9)
Number of SBS associated to treatments
Proportion of SBS associated to treatments
Proportion exposed
43 83 54 38 55 38 67
75 320 76 164 38 169 111
g
Lung (69)
Colon-Rectum (248)
Urinary-tract (54)
Uterus (35)
Ovary (97)
Esophagus (63)
Breast (47)
Number of DBS associated to treatments
Proportion of DBS associated to treatments
42 89 72 92 87 85 75
164 278 75 38 111 74 63
h
Carboplatin
Cisplatin
Oxaliplatin
Capecitabine
Mutations
Mutations per month of exposure

**Figure S8. The contribution of anti-cancer treatments to the mutation burden of tumors**
(a) Comparison of the contribution of different treatments and the aging signature to the mutation burden of tumors originated in different organs. This figure is constructed with SignatureAnalyzer-extracted signatures. Numbers in the x-axis represent the number of tumors represented. The contribution of signatures to the burden of each set of is represented as a filled circle centered at the mean of the distribution, and whiskers extending to its 25th and 75th percentiles. In the stacked barplots below each graph, the fraction of all tumors exposed to the treatment that are coherent are colored, and the numbers below each stacked bar correspond to the total of tumors with activity of the signature according to either method. This panel is equivalent to Figure 3a of the main paper.
(b, c) Contribution in total number (upper) and proportion (lower) of all treatment-associated SBS (b) and DBS (c) to the mutation burden of metastatic tumors originated in different organs in the metastatic tumors cohort. The lower barplot panel represents the percentage of tumors of each organ of origin that are represented in the previous panels, with the absolute numbers below the bars. Numbers in parentheses represent the number of tumors with exposure to the signature included in each distribution.
(d) First column: distribution of the contribution of treatments (and the aging signature) to the mutation burden of tumors exposed to them. Brown boxplots represent the distribution of the contribution by the aging signature, and gray boxplots, that of all treatment-associated signatures. The contribution to the burden of individual tumors is represented as circles colored following the code in Figure 1b. Second column: distribution of the contribution of treatments (and the aging signature) to the mutation burden of tumors during one month of exposed to them.
(e) Same as (a), using the mutational signatures extracted with SigProfiler.
(f, g) Same as (b,c), using the mutational signatures extracted with SigProfiler.
(h) Same as (d), using the mutational signatures extracted with SigProfiler.
Details of the results shown in this figure are provided in Table S2.

### Figure Supplementary 9

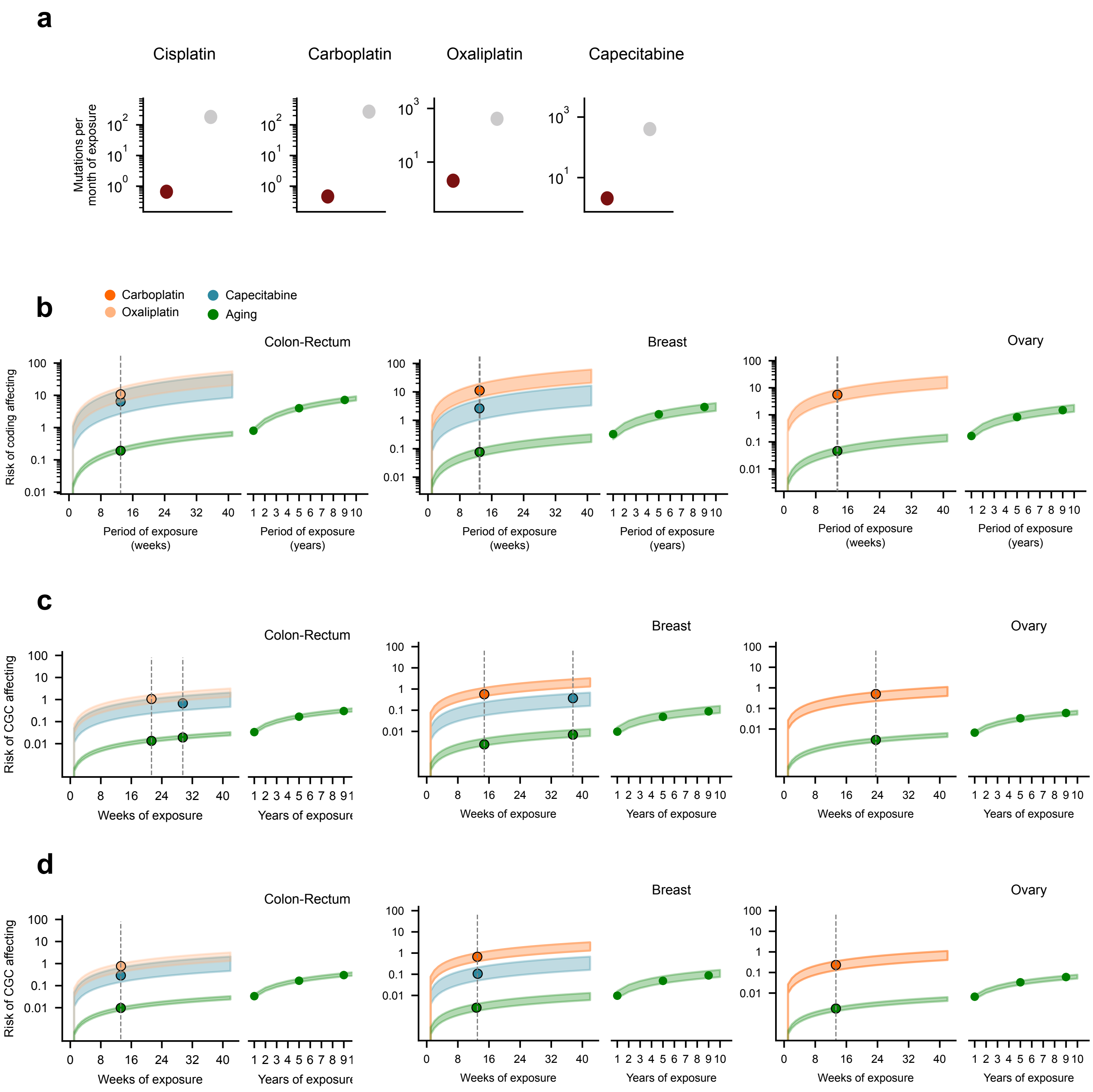

**Figure S9. Risk of coding affecting mutations in cancer genes.**

(a) Contribution of treatment-associated signatures and aging signature to the mutational burden of metastatic tumors. The composition is the same as in Figure 4a, but the duration of the period of exposure to the drugs of all tumors has been taken from the average duration of courses of treatment indicated in clinical guidelines (Table S4). This is why the contribution of each treatment to all tumors is now indicated by a single dot corresponding to the division of the median number of mutations contributed by each treatment by the average duration of courses of treatment indicated in clinical guidelines.

(b) The risk of coding affecting mutations contributed by treatment-associated and aging signatures. Risk values are represented as a range spanning between the 25th and the 75th percentile of the distribution of contribution of signatures to the burden of tumors in one month of exposure (represented in panel d). This panel is similar to Figure 4b. However, here, vertical lines intersecting these risk value ranges are placed at the average duration of courses of treatment indicated in clinical guidelines (Table S4). The range of values of risk for the mutations contributed by the aging signature is extended several years to the right of the graph. Details of the results shown in this figure are provided in Table S3.

(c) Risk of coding mutations affecting cancer genes across tumors contributed by different signatures according to the duration of the exposure of tumors. Computed as in Figure 4c, but restricted to genes in the Cancer Gene Census.

(d) Same as (c), but with the vertical lines intersecting the risk value ranges placed at the average duration of courses of treatment indicated in clinical guidelines (Table S4).
