## Supplementary Note 1 for "The mutational footprints of cancer therapies"

### Supplementary Note 1. Mutational signatures extracted from the cohort of metastatic tumors

#### Supplementary Information

##### 1. Genomic data summary

Table 1 of this document presents a summary of the cohorts of tumor samples employed in the analyses presented in the paper. SBS, DBS and ID detected across the samples in each cohort were obtained from the sources mentioned in the Table. Depending on the number of samples exposed to treatments, and the number of mutations identified across samples, we either carried out a *de novo* mutational signatures extraction, or a fitting of the mutational profile of the cohort on known mutational signatures (see Table 1 at the end of this document).

##### 2. Extraction of mutational signatures

Several methods have been developed in recent years to address the extraction of mutational signatures active across tumor samples[1–6]. The majority of these methods are based on a non-negative matrix factorization (NMF) approach, although they differ in the precise implementation of this approach. As explained in the Methods section, we employed two state-of-the-art NMF-based methods (SignatureAnalyzer and SigProfiler) to extract the mutational signatures active across the tumor samples of the pan-metastatic adult cohort. We decided to use these two methods because a recent comparison of their results in the extraction of the catalog of mutational signatures active across human cancers yielded high degree of agreement, but also produced important differences[2]. The authors of this catalog noted differences in i) the number of signatures extracted by either method --in particular across hypermutated tumors--, ii) the profiles of certain signatures (in particular flat ones), and iii) the attribution of the activity of different signatures across individual tumors.

We run the SignatureAnalyzer using the R implementation provided by the authors[5,6] (https://www.synapse.org/#!Synapse:syn11801488) and following the guidelines they recently presented for the analysis of similar cohorts[2]. On detail, we prepared matrices of counts across 1536 rows (channels) of the penta-nucleotide contexts of SBS, 78 channels for DBS and 83 channels for ID. For the extraction of SBS signatures, we then used a composite matrix integrating the 1697 channels of the three types of variants. On the other hand, for the extraction of DBS and ID signatures, we started with count matrices including only the channels describing each of them. A first step of signatures extraction was carried out for tumor samples with low mutation burden. We considered a sample as highly mutated (and therefore excluded from this step) if it contained a number of variants higher than the median plus 2.5 times the interquartile range of the distribution of mutational burden across the cohort. In a second step, we carried out the extraction of signatures characteristic of highly-mutated tumors, giving the opportunity to signatures already discovered across lowly-mutated tumors to explain part of their mutational profile. As explained by the authors of the method, this two-step process minimizes the possibility of “signature bleeding” or biases introduced by hypermutated samples. Similarly, the attribution of activities to individual samples was carried out separately for lowly-mutated and highly-mutated tumors. Only signatures inferred in the step 1 were available for exposure attribution to lowly mutated samples.

In particular, signature SBS4 (smoking signature) was only available to the metastatic samples of tumors originated in the lung or head and neck, as in a recent paper by the authors of the method[2]. We used in the analysis presented in the paper the exposure attributed to tumors of composite SBS signatures and the signatures extracted separately for DBS and ID.

We were unable to run the SigProfiler signatures extraction using the MATLAB implementation provided by its authors, due to the limitation of the student's license we hold to run in a high-performance computer environment. Therefore, we reimplemented the extraction module in the Julia programming language. To extract the mutational signatures, we constructed matrices of counts across 96 channels of the tri-nucleotide contexts of SBS, 78 channels for DBS and 83 channels for ID. We then carried out the approach to the direct identification of signatures and their attribution to the individual samples in the cohort as described in the original SigProfiler paper[1]. We excluded from the extraction and attribution processes the metastatic tumors originated in the skin and those of unknown origin to avoid the "bleeding" and biases mentioned above. A separate extraction and attribution process is carried out for these samples, but no treatment-associated mutational signatures are identified across them.

**3. Comparison of signatures extracted with both methods**

Using the SignatureAnalyzer, we extracted 72 SBS signatures, 16 DBS signatures and 36 ID mutational signatures across tumor samples in the pan-metastatic adult cohort. On the other hand, the application of SigProfiler to the cohort yielded 26 SBS signatures, 5 DBS signatures and 11 ID signatures. The mutational profiles of all signatures extracted are presented in Figure 1 at the end of this document.

The first step to characterize the SBS, DBS and ID mutational signatures identified by both, SignatureAnalyzer and SigProfiler across the tumor samples of the pan-metastatic adult cohort was their comparison to the catalog of mutational signatures active in human tumors[2] (Fig. 2 at the end of this document). SBS, DBS and ID mutational signatures extracted in this cohort were compared to said catalog. For each signature we indicate the name of the signature in the catalog with highest cosine similarity (above 0.85). Signatures extracted in this cohort for which no comparison yielded a cosine similarity of 0.85 or above were considered new discoveries.

**4. Concordance of treatments being administered to the same patients**

To understand the limitations of our method to disentangle associations between specific treatments and signature activities --with a view towards informed vetting-- we assessed the concordance between treatments across samples. We computed this concordance as the Matthews Correlation Coefficient (MCC) of all the pairs of binary vectors representing treatments, in which we encoded whether a treatment was administered to each patient or not. We carried out this computation across all samples in two settings: i) comparing individual treatments; ii) comparing bins of treatments of same kind, employing the FDA categories described in the Methods section (https://www.accessdata.fda.gov/cder/ndctext.zip).

In Figure 3 of this document we show the concordance for the top 100 most concordant treatment pairs in both settings. We limited our analysis to treatments or groups of treatments with which at least 50 patients were treated. The figure reflects the known fact that some treatment pairs --considering both settings-- are highly co-occurring in standard treatment regimens, which poses a limitation to our analysis. However, for some co-occurring treatments belonging to different treatment bins, the concordance of the respective bins was significantly lower than that of the individual treatments, and overall the concordances between bins were lower. We exploited this feature of the data to further refine our signature-treatment association analysis as described in the Methods section.

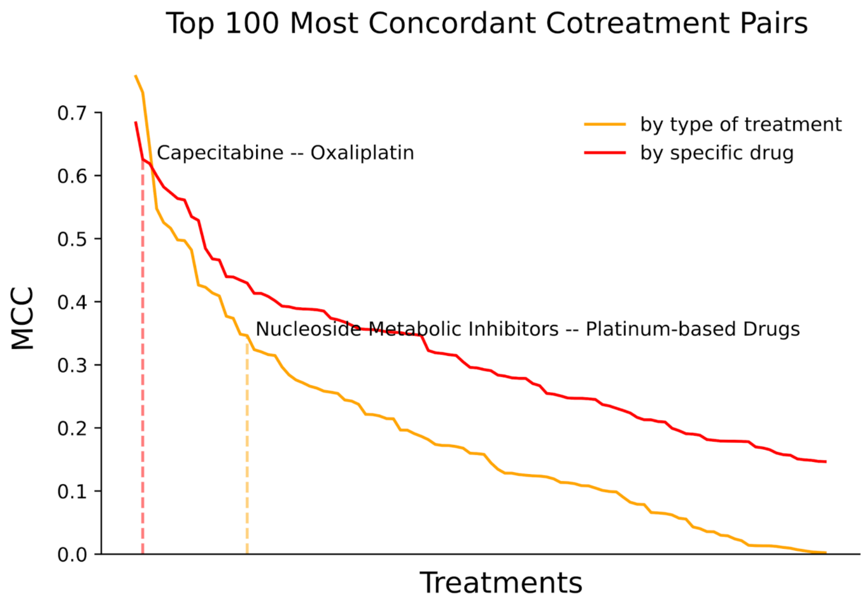


**Figure 3. Concordance in explanation of signature exposure of the 100 most concordant pairs of treatments.**
The yellow line represents the concordance between pairs computed by groups of drugs, while the red line tracks the concordance computed for pairs of individual drugs. The difference between the two methods for the capecitabine-oxaliplatin (or the corresponding nucleoside metabolic inhibitors-platinum-base drugs) are represented by vertical broken lines.

**5. Identification of tumors with coherent exposure to treatment-associated signatures**
We aimed to produce a robust estimate of the individual and collective contribution of different chemotherapies and aging to the mutation burden of tumors (presented in Fig. 4 of the main paper). Therefore, we explored the number of mutations contributed by platinum-based drugs, capecitabine, TMZ and aging (the summation of the products of exposure to associated signatures times the mutation burden; Methods) according to both methods across all tumor samples. We found that the number of mutations contributed by different chemotherapies according to both methods showed a strong correlation (Fig. 4 at the end of this document). However, several samples clearly deviated from this general trend, and the attribution of activity

made by SigProfiler (red) was consistently higher than that of SignatureAnalyzer (blue).

Therefore, to carry out the analysis at hand, we decided to focus on tumor samples with high agreement between both methods. Samples with 15% or less of difference in the number of mutations attributed by each method were considered “coherent”. As expected, coherent samples showed higher overall correlation between the exposures attributed by both methods. However, this reduced the number of tumor samples available to estimate the contribution of different treatments to the mutation burden.

Graphs presented in Figure 4 of the main paper focus on coherent tumors, and the exposure of each tumor to each signature used to compute the contribution of treatments to the mutation burden is represented by the mean of those computed by SignatureAnalyzer and SigProfiler. Similar figures for the contribution of treatment-associated signatures computed from the activities attributed by both methods to all tumors.

**Table 1. Description of cohorts of tumors included in the study**

| Cohort ID | Number of tumor samples | Number of samples exposed to treatments | Type of sequencing | Source | Method used to identify signatures |
|---|---|---|---|---|---|
| Pan-metastatic adult | 3131 | At least 1962 | WGX | HMF | Extraction |
| St. Jude cohort | 635 | At least 12 | WGX | St. Jude Cloud | Extraction |
| Glioblastomas | 28 | 16 | WEX | Wang J et al (2016) | Fitting |

### Figure Legends

**Figure 1. Profile of all SBS, DBS and ID signatures identified in the pan-metastatic adult cohort using SignatureAnalyzer and SigProfiler**

**Figure 2. Mutational signatures extracted from metastatic adult tumors**
(a) Mutational signatures extracted using the SignatureAnalyzer method. SBS, DBS and ID signatures are presented in separate heatmaps. Each row of the heatmap represents a signature active in the cohort across tumors originated in different organs (columns), which activity is represented by a circle with diameter proportional to the fraction of tumors with signature activity and the color representing the contribution of the signature to the mutation burden of tumors.
(b) Same as panel (a) for mutational signatures extracted using the SigProfiler.
(c) Mutational signatures extracted from the colorectal cohort using a not-NMF approach. Similar repertoire of mutational signatures is extracted from the colorectal cohort as with the SignatureAnalyzer and the SigProfiler.

**Figure 4. Selection of coherent tumors (associated to figure 4 of the main paper) according to the activity of signatures attributed by both methods**
Left panels show the agreement of both methods in the attribution of the exposure of tumors to treatment-associated signatures. Each pair of circles connected by a line represents the exposure

attributed by both methods to a tumor. Red circles represent the exposure attributed by SigProfiler, while blue circles represent the exposure attributed by SignatureAnalyzer. Middle panels show the correlation between the exposure attributed by both methods to all tumors, while right panels present the correlation of the exposure attributed by both methods to coherent tumors.

**Figure 1**

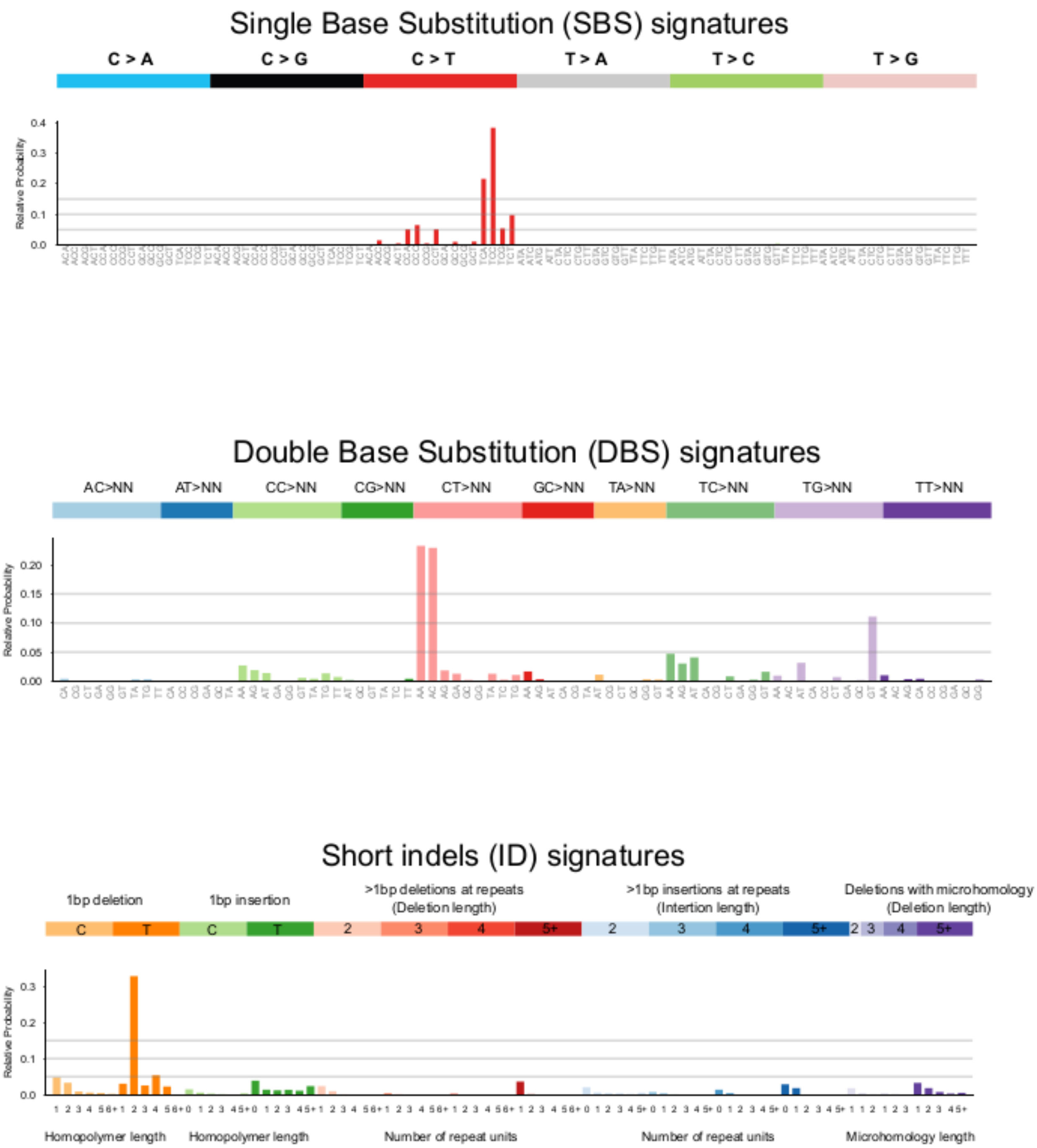
Single Base Substitution (SBS) signatures
C > A
C > G
C > T
T > A
T > C
T > G
Relative Probability
Double Base Substitution (DBS) signatures
AC>NN
AT>NN
CC>NN
CG>NN
CT>NN
GC>NN
TA>NN
TC>NN
TG>NN
TT>NN
Relative Probability
Short indels (ID) signatures
1bp deletion
1bp insertion
>1bp deletions at repeats
(Deletion length)
>1bp insertions at repeats
(Intertion length)
Deletions with microhomology
(Deletion length)
Relative Probability
Homopolymer length
Homopolymer length
Number of repeat units
Number of repeat units
Microhomology length

### SignatureAnalyzer - SBS

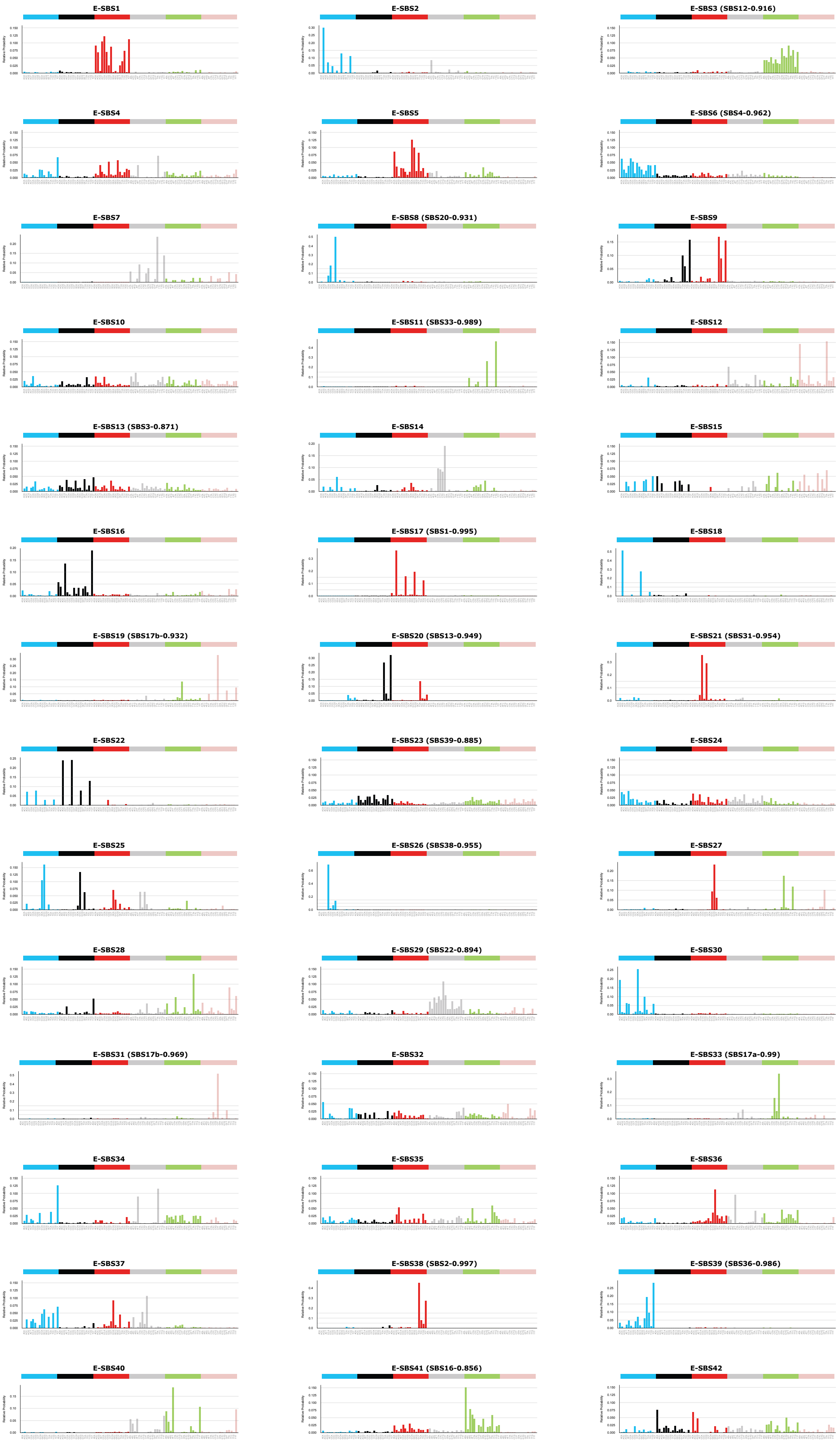

### SignatureAnalyzer - SBS (continuation)

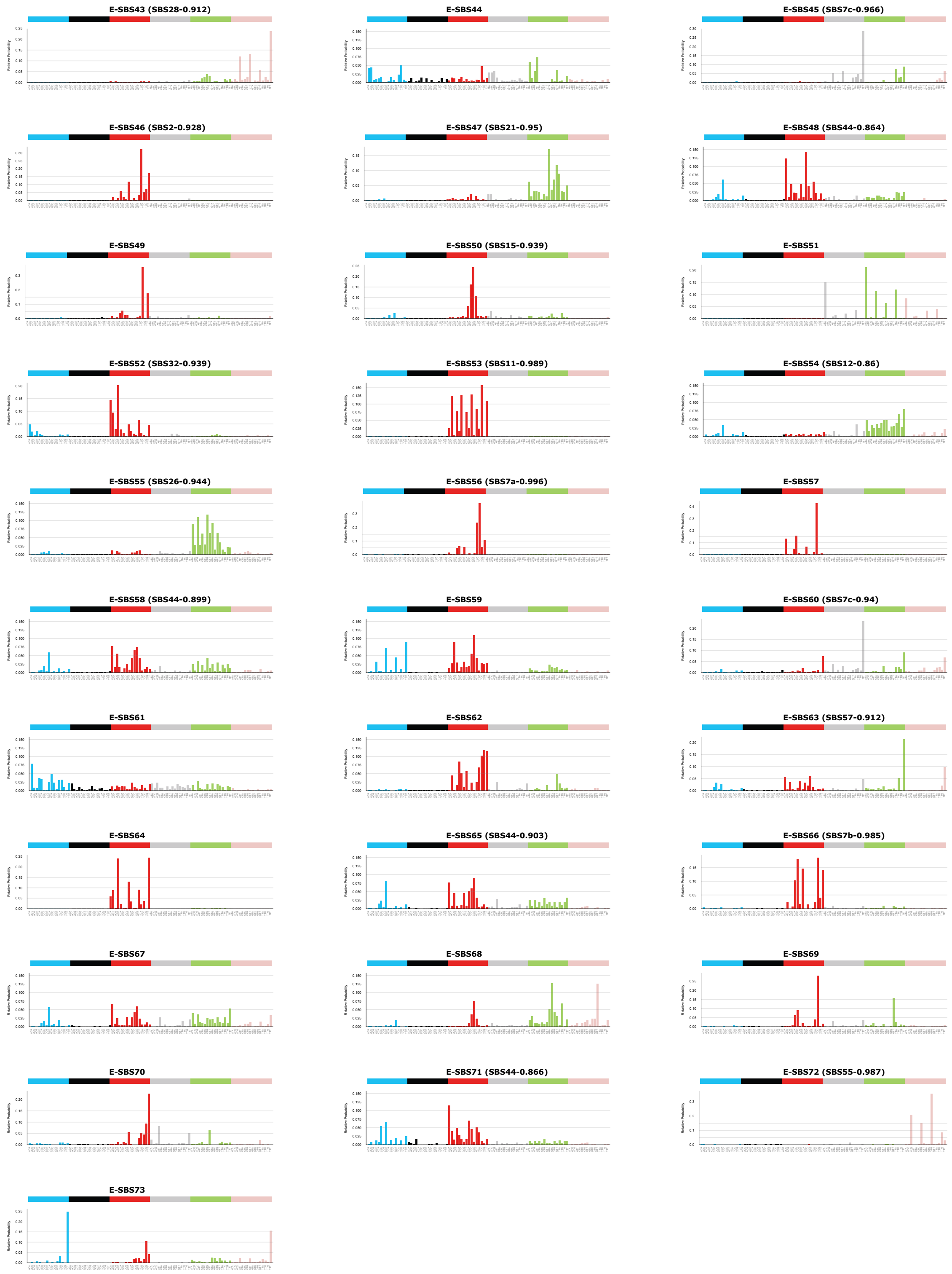

### SignatureAnalyzer-DBS

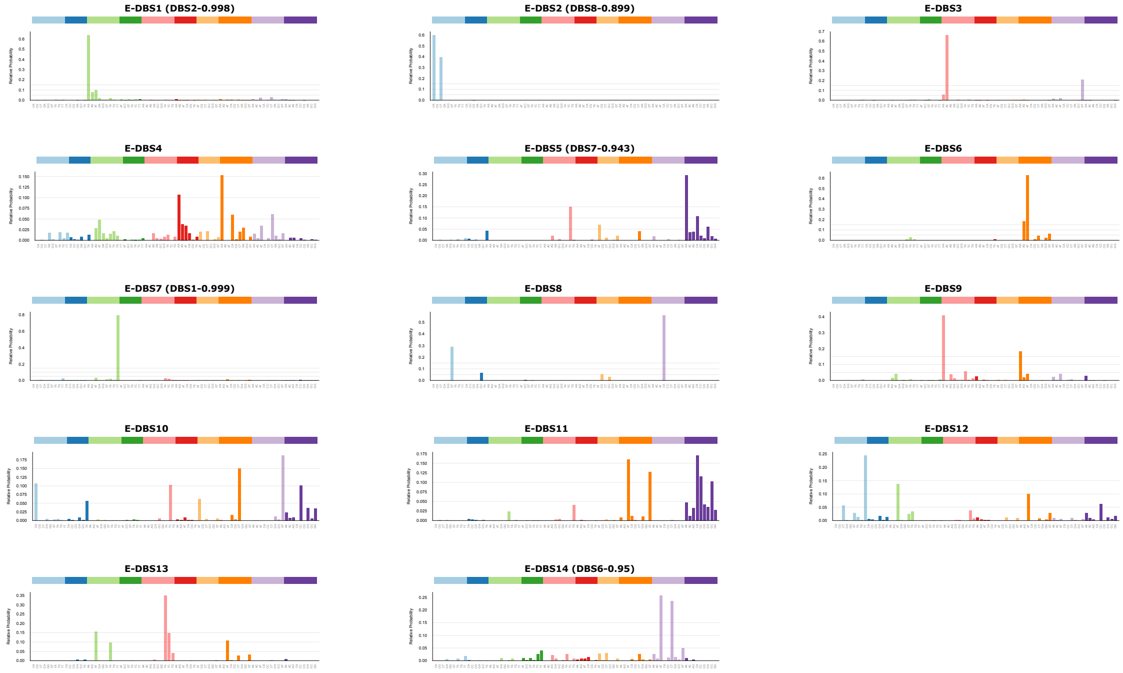

### SignatureAnalyzer-ID

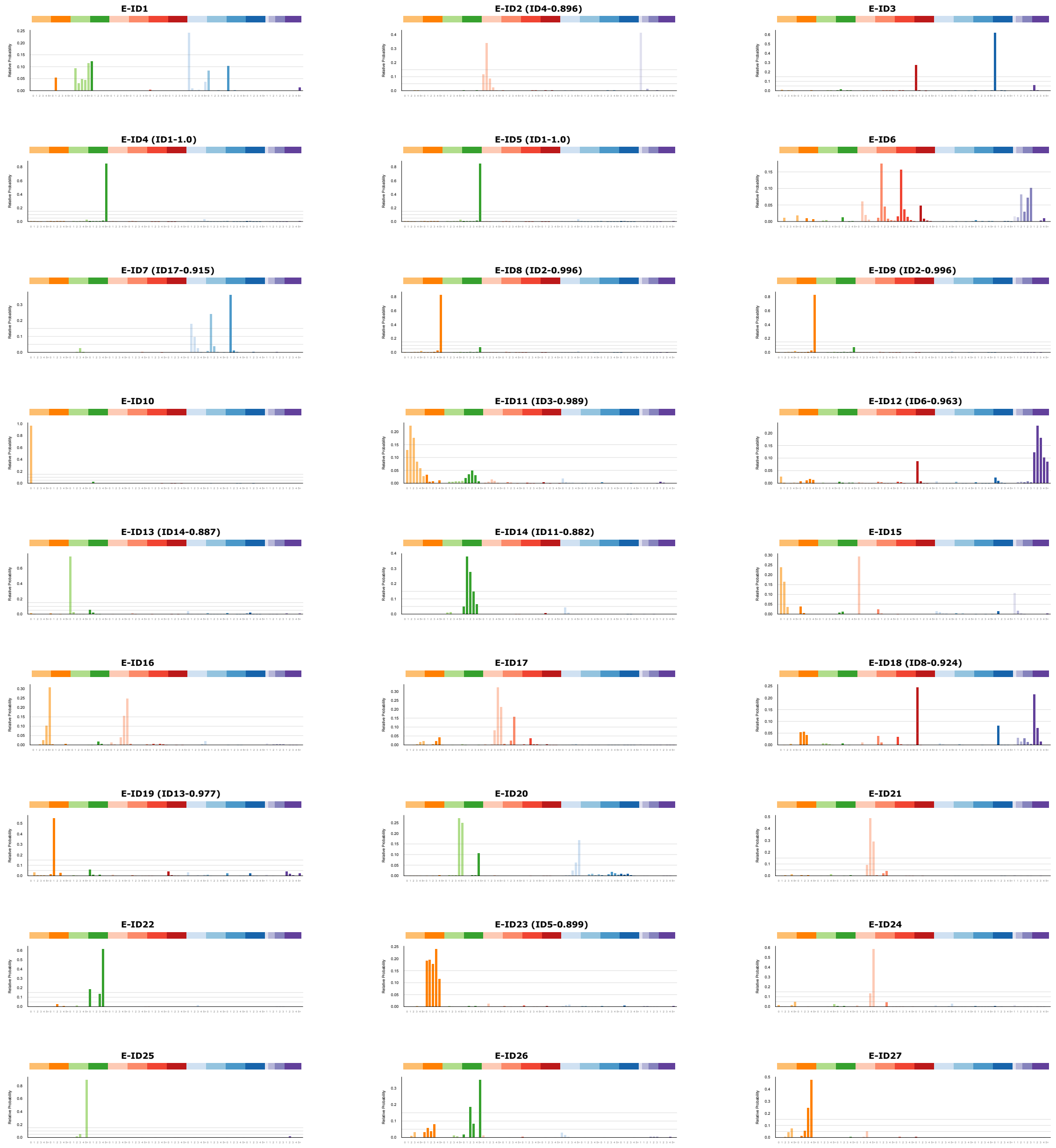

### SigProfiler-SBS

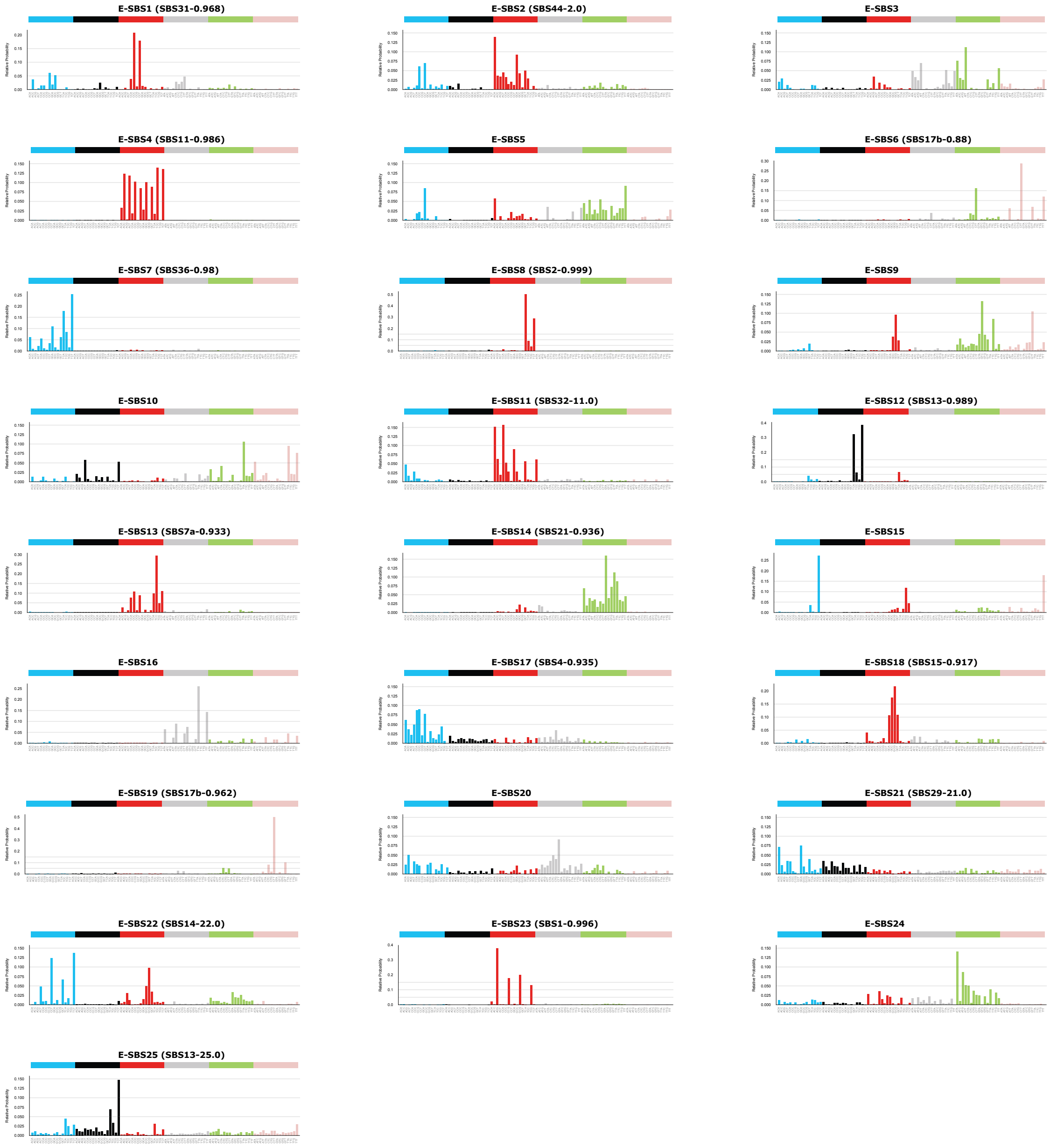

### SigProfiler-DBS

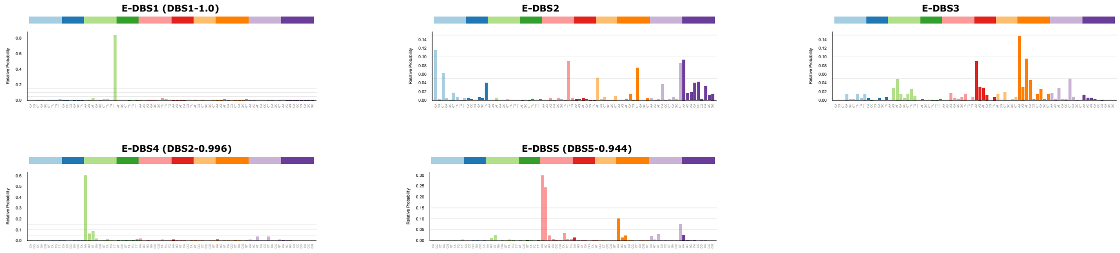

### SigProfiler-ID

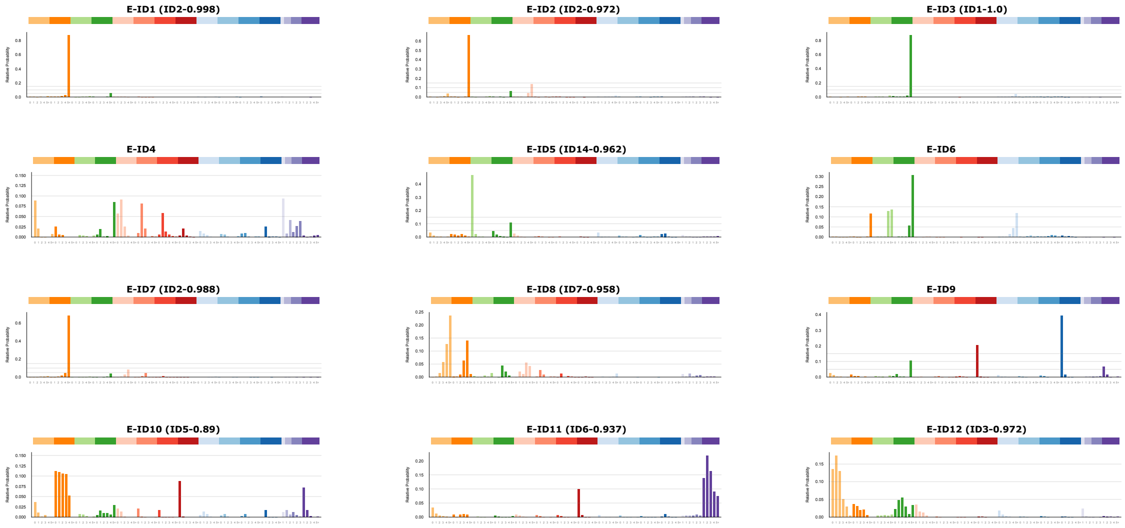

### Figure 2

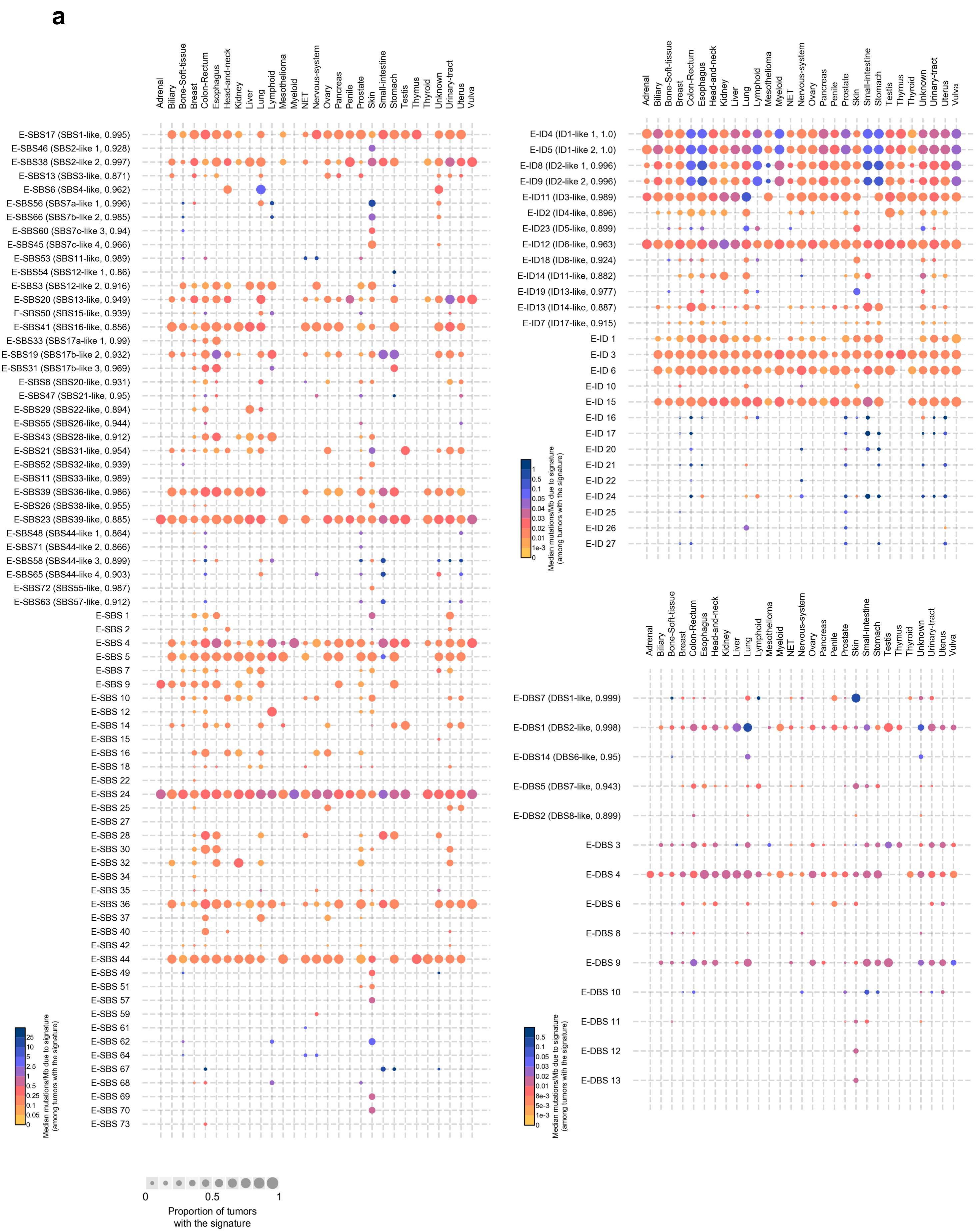
a
Adrenal
Biliary
Bone-Soft-tissue
Breast
Colon-Rectum
Esophagus
Head-and-neck
Kidney
Liver
Lung
Lymphoid
Mesothelioma
Myeloid
NET
Nervous-system
Ovary
Pancreas
Penile
Prostate
Skin
Small-intestine
Stomach
Testis
Thymus
Thyroid
Unknown
Urinary-tract
Uterus
Vulva
E-SBS17 (SBS1-like, 0.995)
E-SBS46 (SBS2-like 1, 0.928)
E-SBS38 (SBS2-like 2, 0.997)
E-SBS13 (SBS3-like, 0.871)
E-SBS6 (SBS4-like, 0.962)
E-SBS56 (SBS7a-like 1, 0.996)
E-SBS66 (SBS7b-like 2, 0.985)
E-SBS60 (SBS7c-like 3, 0.94)
E-SBS45 (SBS7c-like 4, 0.966)
E-SBS53 (SBS11-like, 0.989)
E-SBS54 (SBS12-like 1, 0.86)
E-SBS3 (SBS12-like 2, 0.916)
E-SBS20 (SBS13-like, 0.949)
E-SBS50 (SBS15-like, 0.939)
E-SBS41 (SBS16-like, 0.856)
E-SBS33 (SBS17a-like 1, 0.99)
E-SBS19 (SBS17b-like 2, 0.932)
E-SBS31 (SBS17b-like 3, 0.969)
E-SBS8 (SBS20-like, 0.931)
E-SBS47 (SBS21-like, 0.95)
E-SBS29 (SBS22-like, 0.894)
E-SBS55 (SBS26-like, 0.944)
E-SBS43 (SBS28-like, 0.912)
E-SBS21 (SBS31-like, 0.954)
E-SBS52 (SBS32-like, 0.939)
E-SBS11 (SBS33-like, 0.989)
E-SBS39 (SBS36-like, 0.986)
E-SBS26 (SBS38-like, 0.955)
E-SBS23 (SBS39-like, 0.885)
E-SBS48 (SBS44-like 1, 0.864)
E-SBS71 (SBS44-like 2, 0.866)
E-SBS58 (SBS44-like 3, 0.899)
E-SBS65 (SBS44-like 4, 0.903)
E-SBS72 (SBS55-like, 0.987)
E-SBS63 (SBS57-like, 0.912)
E-SBS 1
E-SBS 2
E-SBS 4
E-SBS 5
E-SBS 7
E-SBS 9
E-SBS 10
E-SBS 12
E-SBS 14
E-SBS 15
E-SBS 16
E-SBS 18
E-SBS 22
E-SBS 24
E-SBS 25
E-SBS 27
E-SBS 28
E-SBS 30
E-SBS 32
E-SBS 34
E-SBS 35
E-SBS 36
E-SBS 37
E-SBS 40
E-SBS 42
E-SBS 44
E-SBS 49
E-SBS 51
E-SBS 57
E-SBS 59
E-SBS 61
E-SBS 62
E-SBS 64
E-SBS 67
E-SBS 68
E-SBS 69
E-SBS 70
E-SBS 73
Median mutations/Mb due to signature (among tumors with the signature)
25
10
5
2.5
1
0.5
0.25
0.1
0.05
0
0
0.5
1
Proportion of tumors with the signature
E-ID4 (ID1-like 1, 1.0)
E-ID5 (ID1-like 2, 1.0)
E-ID8 (ID2-like 1, 0.996)
E-ID9 (ID2-like 2, 0.996)
E-ID11 (ID3-like, 0.989)
E-ID2 (ID4-like, 0.896)
E-ID23 (ID5-like, 0.899)
E-ID12 (ID6-like, 0.963)
E-ID18 (ID8-like, 0.924)
E-ID14 (ID11-like, 0.882)
E-ID19 (ID13-like, 0.977)
E-ID13 (ID14-like, 0.887)
E-ID7 (ID17-like, 0.915)
E-ID 1
E-ID 3
E-ID 6
E-ID 10
E-ID 15
E-ID 16
E-ID 17
E-ID 20
E-ID 21
E-ID 22
E-ID 24
E-ID 25
E-ID 26
E-ID 27
Median mutations/Mb due to signature (among tumors with the signature)
1
0.5
0.1
0.05
0.04
0.03
0.02
0.01
1e-3
0
E-DBS7 (DBS1-like, 0.999)
E-DBS1 (DBS2-like, 0.998)
E-DBS14 (DBS6-like, 0.95)
E-DBS5 (DBS7-like, 0.943)
E-DBS2 (DBS8-like, 0.899)
E-DBS 3
E-DBS 4
E-DBS 6
E-DBS 8
E-DBS 9
E-DBS 10
E-DBS 11
E-DBS 12
E-DBS 13
Median mutations/Mb due to signature (among tumors with the signature)
0.5
0.1
0.05
0.03
0.02
0.01
8e-3
5e-3
1e-3
0

**b**

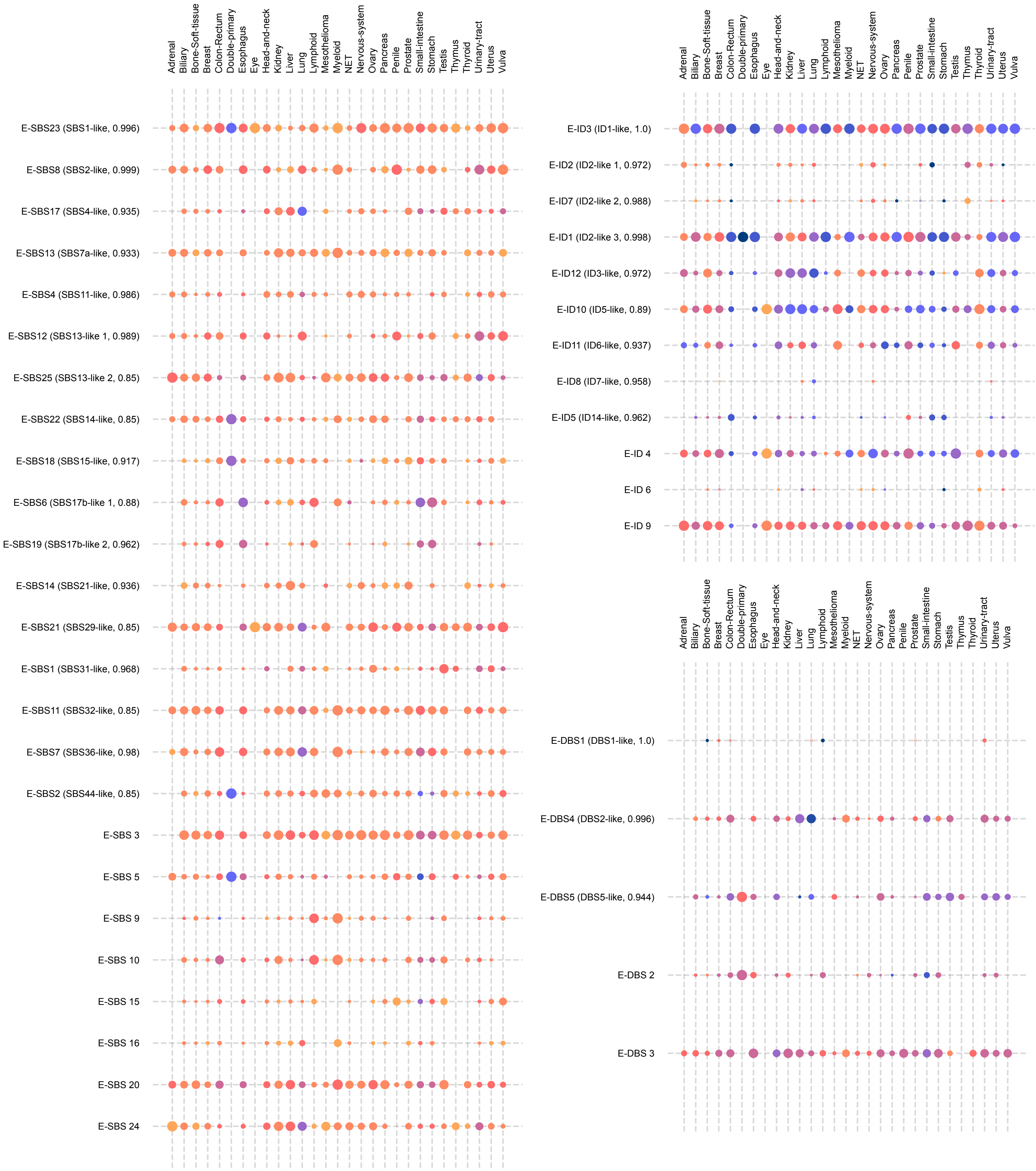

Adrenal
Biliary
Bone-Soft-tissue
Breast
Colon-Rectum
Double-primary
Esophagus
Eye
Head-and-neck
Kidney
Liver
Lung
Lymphoid
Mesothelioma
Myeloid
NET
Nervous-system
Ovary
Pancreas
Penile
Prostate
Small-intestine
Stomach
Testis
Thymus
Thyroid
Urinary-tract
Uterus
Vulva
E-SBS23 (SBS1-like, 0.996)
E-SBS8 (SBS2-like, 0.999)
E-SBS17 (SBS4-like, 0.935)
E-SBS13 (SBS7a-like, 0.933)
E-SBS4 (SBS11-like, 0.986)
E-SBS12 (SBS13-like 1, 0.989)
E-SBS25 (SBS13-like 2, 0.85)
E-SBS22 (SBS14-like, 0.85)
E-SBS18 (SBS15-like, 0.917)
E-SBS6 (SBS17b-like 1, 0.88)
E-SBS19 (SBS17b-like 2, 0.962)
E-SBS14 (SBS21-like, 0.936)
E-SBS21 (SBS29-like, 0.85)
E-SBS1 (SBS31-like, 0.968)
E-SBS11 (SBS32-like, 0.85)
E-SBS7 (SBS36-like, 0.98)
E-SBS2 (SBS44-like, 0.85)
E-SBS 3
E-SBS 5
E-SBS 9
E-SBS 10
E-SBS 15
E-SBS 16
E-SBS 20
E-SBS 24
E-ID3 (ID1-like, 1.0)
E-ID2 (ID2-like 1, 0.972)
E-ID7 (ID2-like 2, 0.988)
E-ID1 (ID2-like 3, 0.998)
E-ID12 (ID3-like, 0.972)
E-ID10 (ID5-like, 0.89)
E-ID11 (ID6-like, 0.937)
E-ID8 (ID7-like, 0.958)
E-ID5 (ID14-like, 0.962)
E-ID 4
E-ID 6
E-ID 9
E-DBS1 (DBS1-like, 1.0)
E-DBS4 (DBS2-like, 0.996)
E-DBS5 (DBS5-like, 0.944)
E-DBS 2
E-DBS 3

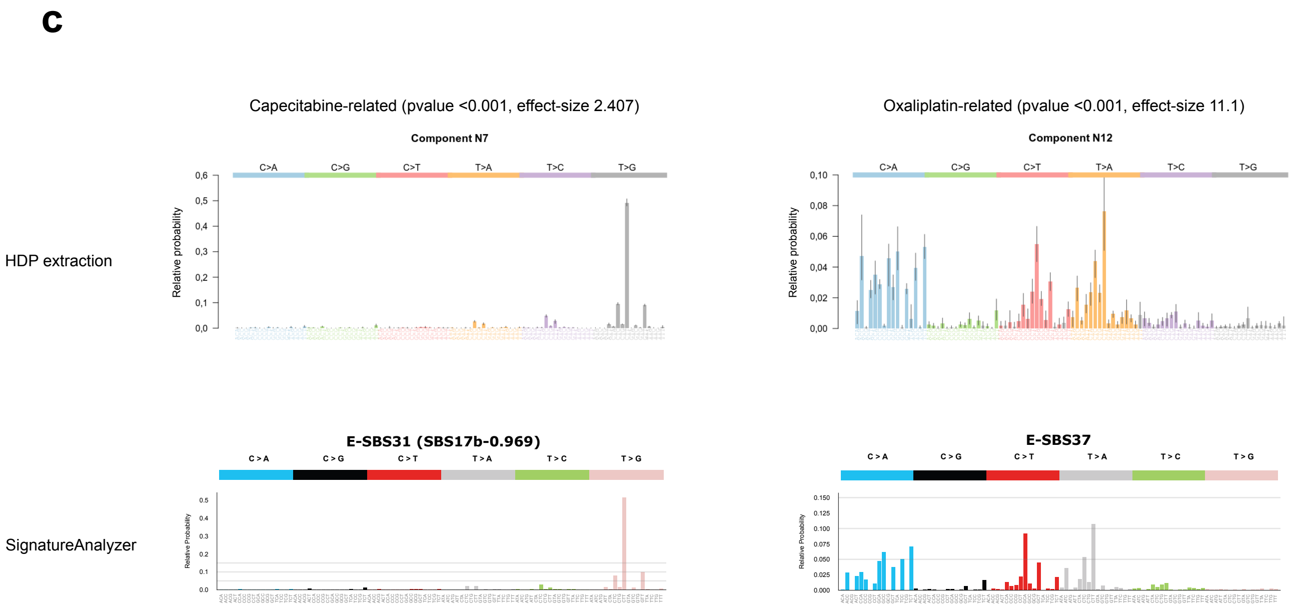
c
Capecitabine-related (pvalue <0.001, effect-size 2.407)
Oxaliplatin-related (pvalue <0.001, effect-size 11.1)
HDP extraction
Component N7
Component N12
C>A
C>G
C>T
T>A
T>C
T>G
Relative probability
SignatureAnalyzer
E-SBS31 (SBS17b-0.969)
E-SBS37
Relative Probability

### Figure 4

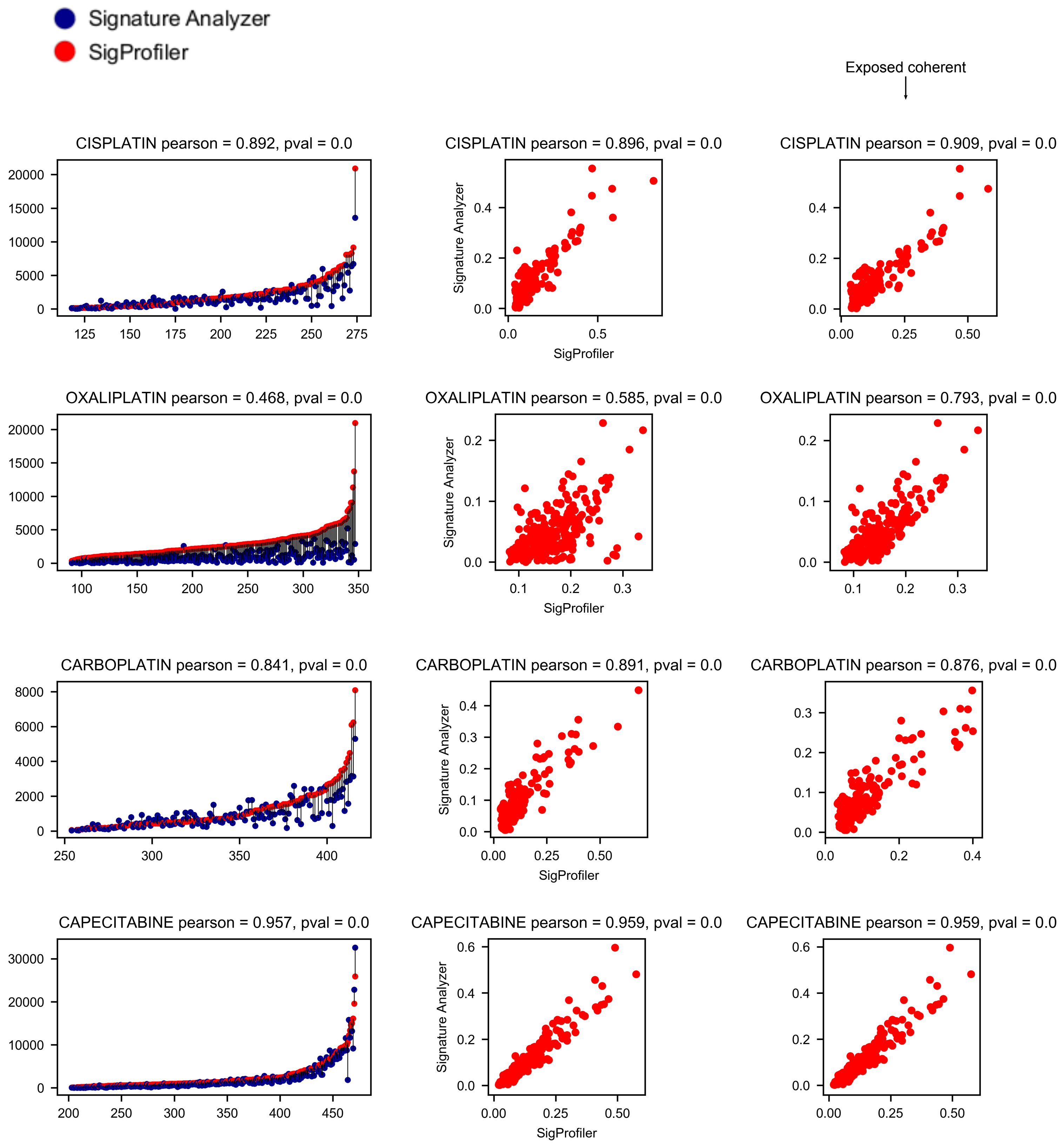
Signature Analyzer
SigProfiler
Exposed coherent
CISPLATIN pearson = 0.892, pval = 0.0
CISPLATIN pearson = 0.896, pval = 0.0
CISPLATIN pearson = 0.909, pval = 0.0
OXALIPLATIN pearson = 0.468, pval = 0.0
OXALIPLATIN pearson = 0.585, pval = 0.0
OXALIPLATIN pearson = 0.793, pval = 0.0
CARBOPLATIN pearson = 0.841, pval = 0.0
CARBOPLATIN pearson = 0.891, pval = 0.0
CARBOPLATIN pearson = 0.876, pval = 0.0
CAPECITABINE pearson = 0.957, pval = 0.0
CAPECITABINE pearson = 0.959, pval = 0.0
CAPECITABINE pearson = 0.959, pval = 0.0
Signature Analyzer
SigProfiler
