## Supplementary Note 2 for "The mutational footprints of cancer therapies"

### Supplementary Note 2: Validation of signature extraction and treatment-signature association with synthetic datasets

The mutational footprint of cancer therapies

### Table of Contents


**Abstract** **2**

**1. Aim and conceptual framework** **3**

**2. Generation of synthetic datasets** **4**
- Mutational catalogues 4
- Procedure to simulate mutational data 6

**3. Signatures extraction** **6**
- Conclusions 9

**4. Identification of treatment-signature associations** **9**
- Significance of the detected associations 9
- Effect-size and significance 9
- Introducing errors in treatment labels 13
- Conclusions 13

**5. Co-treatment analysis: injected signal against placebo** **17**
- Symmetric one-to-one setting 18
- Symmetric many-to-one setting 18
- Asymmetric one-to-one setting 18
- Asymmetric many-to-one setting 18
- Conclusions 18

**6. Recovery of the activity of injected signatures** **24**
- Conclusions 24

**7. Signature activity in non-treated samples** **30**
- Conclusions 30

**References** **32**

#### Abstract

We built synthetic datasets of mutations that are similar to the metastatic tumors analyzed with regard to the composition of mutational signatures. We then injected a known number of mutations drawn from the mutational profile of a foreign signature to a known number of samples of these synthetic datasets. We thus control the number of samples bearing the mutational footprint of the drug, the number of drug-induced mutations present in each sample, the signature of the drug-induced mutations and the number of samples known to have undergone treatment (allowing for discrepancies between these two parameters). Using these synthetic datasets, we tested i) the extraction of drug-associated signatures, ii) the detection of the mutational footprints of drugs through the regression method, iii) the identification of the correct etiology of the signature in the case of tumors exposed to co-treatments, and iv) the accuracy of the estimation of the number of mutations contributed by drugs to the burden of tumors. In the analyses, we challenged our entire methodological setting with fluctuations in the synthetic data reflecting a variety of common scenarios. The analysis of these synthetic datasets demonstrates that the modelling approach correctly identifies the foreign signatures as the molecular footprints of anti-cancer treatments within a wide range of numbers of exposed samples. The methodology is robust to systematic errors such as miss-annotation of treatments or lack of activity of the associated signatures in a subset of exposed samples. It is also able to estimate the mutational burden contributed by the treatment within acceptable confidence intervals. The results of these analyses have been useful to fine-tune the parameters of the methodologies developed to detect the mutational footprint and mutation burden of treatments.

### 1. Aim and conceptual framework

To demonstrate the validity of the approach developed to identify the mutational footprints of anti-cancer treatments, we designed several synthetic datasets similar to the analyzed cohort of metastatic tumors. The mutations in all synthetic datasets are drawn following the distribution of mutational profiles (thus respecting underlying mutational signatures) and mutation burden observed across primary tumors (two datasets with different genetic makeup were selected; see Section 2). To simulate the footprint of a treatment, we then injected to a subset of the samples in the synthetic datasets a number of mutations drawn from the profile of a signature foreign to the dataset (two signatures were chosen to this end; see Section 2). A number of synthetic datasets including one of two foreign signatures in varying number of samples were constructed following this rationale.

The synthetic datasets were thus used to test key points of the approach. We first tested the capability of one of the signatures extraction methods used in the paper to recover the injected signature (see Section 3). Secondly, we generated labels distinguishing the samples to which the foreign signature had been injected from the rest, and tested the ability of the regression method to correctly recover the relationship between the foreign signature and the sample labels (see Section 4). Bringing this test closer to the real-life scenario, we re-shuffled part of the labels identifying treated signatures and verified how this added noise affected the detection of the association between the signature and the treatment. We also checked the performance of the method developed to discern the true etiology of the footprint in cases of samples exposed to two overlapping treatments (with variable degrees of overlap; see Section 5). Finally, we transformed the activity of the foreign signature detected across exposed samples into a number of treatment-associated mutations (as in the paper) and compared it to the number of mutations that had been actually injected to each sample. This was done to test the accuracy of our estimates of the contribution of treatments to the mutational burden of tumors (see Section 6). We also explored how much “bleeding” --i.e., activity of injected signatures recovered from non-injected samples is introduced in the extraction (see Section 7).

#### 2. Generation of synthetic datasets

We built a collection of mutational catalogues, i.e., a tabular dataset of mutation counts across trinucleotide-contexts of a set of tumor samples in a cohort. Each catalogue represents the mutational profile of samples of a typical primary tumor type. In a subset of the tumors of this cohort where, for a subset of the samples, a known mutational signal is added, i.e., each catalogue of our collection would come about as a mixture of two mutational signals: i) a typical mutational signal of a primary tumor; ii) a known profile --reminiscent of the mutagenic effect of some etiology that is injected to only a subset of the samples.

##### Mutational catalogues

All catalogues were created with 300 samples. A baseline mutational signal for each catalogue was generated using PCAWG mutational catalogues as a reference [1]. Two sample sets were used to generate these baseline mutational signals: i) the breast adenocarcinoma --hereinafter Breast-- cohort with 198 samples, and ii) the combination of the breast, lung and colorectal adenocarcinoma cohorts --hereinafter Breast-Lung-Colorectal-- with 296 samples (Fig. 1).

As foreign single base substitutions (SBS) signatures, we selected SBS9 (related to AID-driven somatic hypermutation), and SBS31 (related to exposure to platinum-based drugs; see Fig. 2) from the collection of mutational signatures found in PCAWG [1]. For short, in this document we will call this additional exposure the “injected” exposure or signal.

Among the 300 samples of each synthetic dataset, only a subset of samples have injected exposure. We considered 5 sizes of this subset of samples with injected exposure: 10, 25, 50, 100, 150. In other words, we built different synthetic datasets with between 3% and 50% of the samples bearing mutations of the injected signature.

For each possible configuration defined by the mutational catalog of the primary tumors (2: Breast and Breast-Lung-Colorectal), injected signals (2), and number of samples with injected exposure (5), we generated several random draws to cope with stochasticity. For the Breast model we drew 10 replicates of each configuration. For the Breast-Lung-Colorectal model we drew 25 replicates for each configuration. Thus we analyzed a total of 700 mutational catalogues.

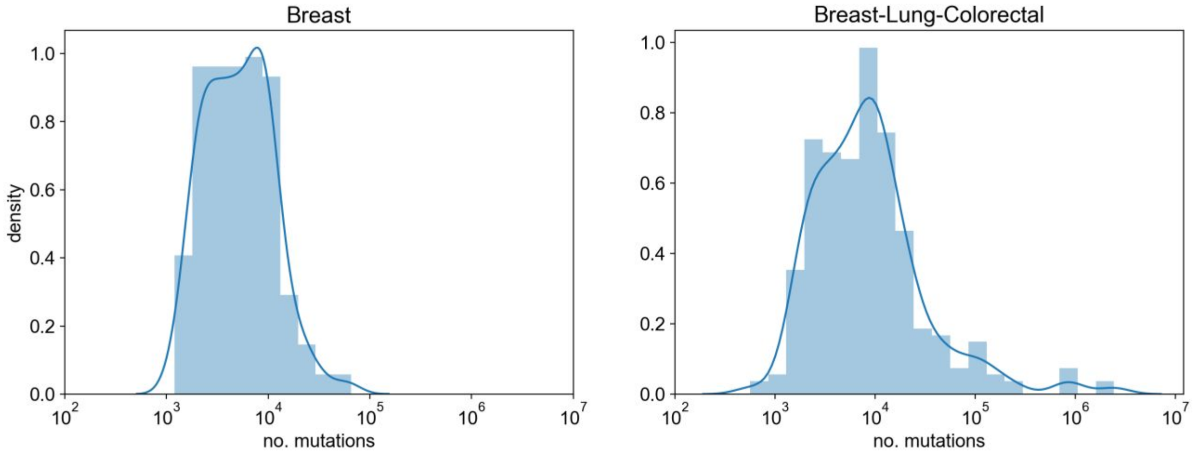


**Fig. 1.** Burden distributions of two reference cohorts used as templates to generate the simulated datasets: PCAWG Breast and Breast-Lung-Colorectal.

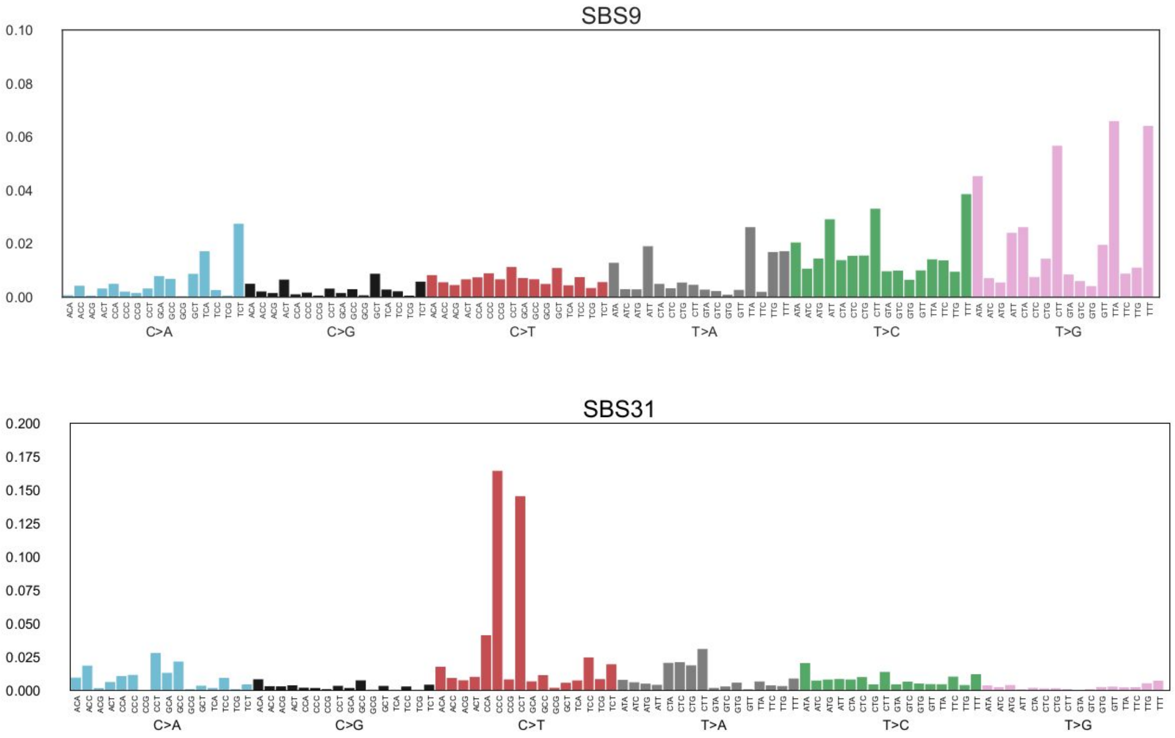


**Fig. 2.** Profile of 96-channel signatures SBS9 and SBS31 from the PCAWG catalogue [1].

#### Procedure to simulate mutational data

Given the parameters of the catalogue --primary tumor-type (P), injected signal (S) and number of samples with activity of the injected signal (T)-- each mutational catalogue is drawn according to the following procedure:

- From the PCAWG cohort matching P we randomly chose 300 samples with replacement from a uniform distribution. These samples --and mutation count thereof-- will be taken as templates for generating our own synthetic catalogue.
- For each sample $s$ drawn from P in the previous step a new 96-channel catalogue was randomly generated by drawing as many mutations as $s$ has --say $n(s)$-- from the 96-class multinomial distribution corresponding to the mixture of 96-channel signatures corresponding to $s$ according to the analysis of mutational processes conducted in PCAWG [1].
- Then we randomly selected T synthetic samples, to inject activity of S.
- Finally, for each sample selected in the previous step, we randomly drew $p$ from a beta distribution fit with the platinum-based exposures that were estimated as part of our analysis (Supplementary Table S2). The value $p$ is meant to represent a conservative typical proportion of mutations of a sample's catalogue that would be attributable to the signal S (see Fig. 3). Accordingly, for each synthetic sample we drew the corresponding number of mutations from the 96-class multinomial distribution corresponding to signature S and we added them to the catalogue.

### 3. Signatures extraction

We extracted the signatures active across the 700 synthetic mutational catalogues described above using SignatureAnalyzer [2]. Thus for each catalogue we obtained a collection of mutational signatures and a mapping of the samples to the activities --number of mutations-- contributed by each of the signatures.

We first analyzed whether the extraction method was able to recover the signals injected in subsets of samples. We proceeded by comparing all the signatures obtained by the deconstruction method against each of the injected profiles via cosine similarity. We deemed the one with highest similarity the reconstructed signature that represented the injected profile. In some cases, however, more than one signature yielded comparable cosine similarity values with the injected signal (Figs. 4a and b).

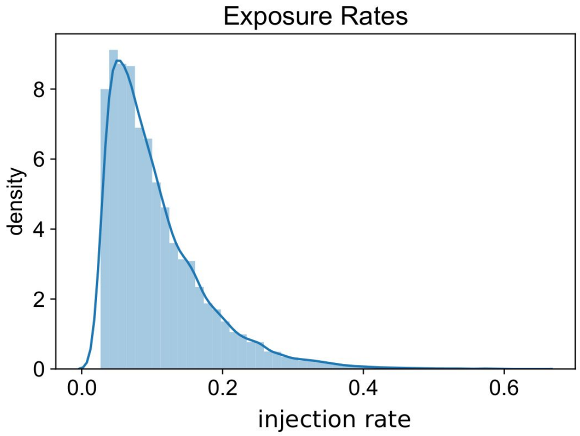


**Fig. 3.** Distribution of exposure rates used to generate the stochastic burden of the injected signatures. The histogram and density represent 1000 random variates of the beta distribution with parameters a~1.28, b~833471, loc~0.027, scale~51675. The injection rate --representing the proportion of mutations per sample that can be explained by the injected signature--distribution was fit with the observed exposures to platinum-based associated signatures in the main paper.

**a) Breast**

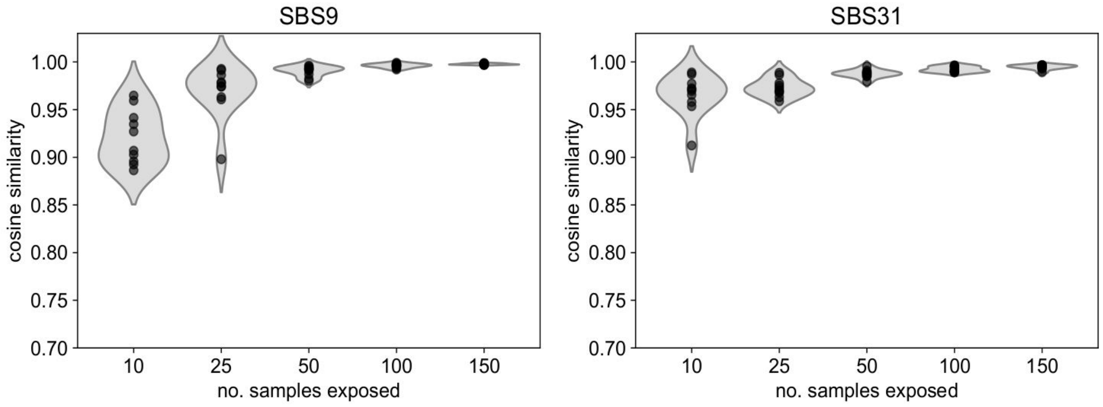


**b) Breast-Lung-Colorectal**

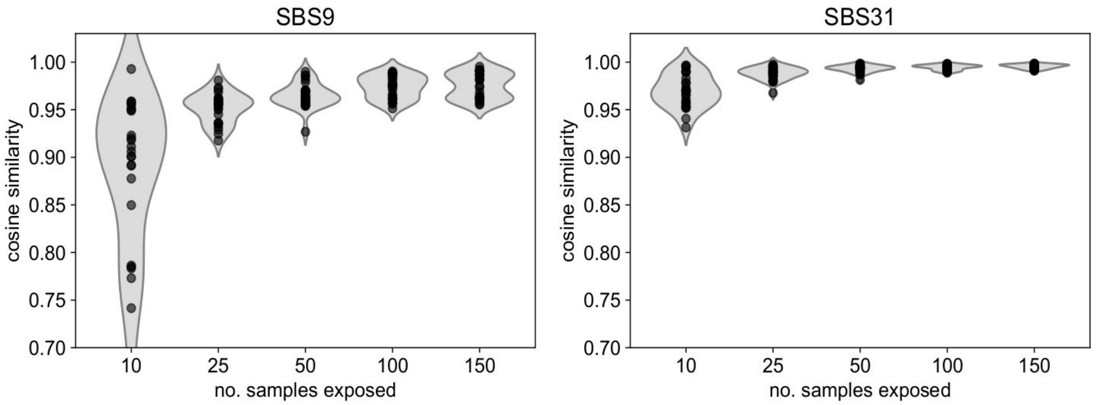


**Fig. 4.** Cosine similarities of the reconstructed signature to the real injected profiles (SBS9 and SBS31).

#### Conclusions

Unsupervised mutational signature extraction is able to recover the injected signature (mean cosine similarity across replicates above 0.9) even in cases in which the number of samples subject to injected exposure is below 10% of the total number of samples (Figs. 5a and b). As expected, the higher the fraction of samples bearing the injected signal, the higher the accuracy.

### 4. Identification of treatment-signature associations

As stated above, we also wanted to evaluate the performance of the regression method employed to identify signatures that are associated to treatments administered to the patients in the cohort. This is what we refer to as “Step 1” in the Methods section of the main paper. To this end, we added labels distinguishing samples that have been subject to a treatment (injected signature activity) from those that have not. We therefore wanted to ascertain how well we could predict these labels using only the activity of signatures extracted from the synthetic datasets. To render the exercise more similar to the real-life scenario, we introduced a certain degree of error (see below) in the assignment of labels to samples with injected signatures activity.

#### Significance of the detected associations

Our logistic regression ensemble approach starts by 1000 randomly drawn sets of tumors which are balanced and stratified by tumor type. For each set, regularized logistic fitting yields parameter estimates for all the active signatures, thus a collection of 1000 parameters --abusing notation we refer to them as log-odds-ratio or LOR-- are obtained for each active signature. If the values are positive, this is indicative of a positive association between the signature activity and the label. For a signature, thus, the proportion of LORs below zero is used as an empirical P-value for the signature-treatment association. In the following plots (Figs. 5a and b), those at the left show the mean LOR of the deconstructed injected signal (named SBSX in the plots) vs the mean LOR from the rest (other), and those at the right, the resulting empirical P-values.

#### Effect-size and significance

For the analysis described above, we also display the effect-size --as the fold change between the mean exposure values of those samples labelled injected and those labelled non-injected-- alongside the empirical P-values (Figs. 6a and b) in a volcano plot like the one supplied in Fig. 1e of the main paper. Dots colored in red correspond to those tests expected to raise a significant signal-to-noise ratio --i.e., how well the activity of the reconstructed signature that is most similar to the true injected signal predicts the injected labels.

#### a) Breast

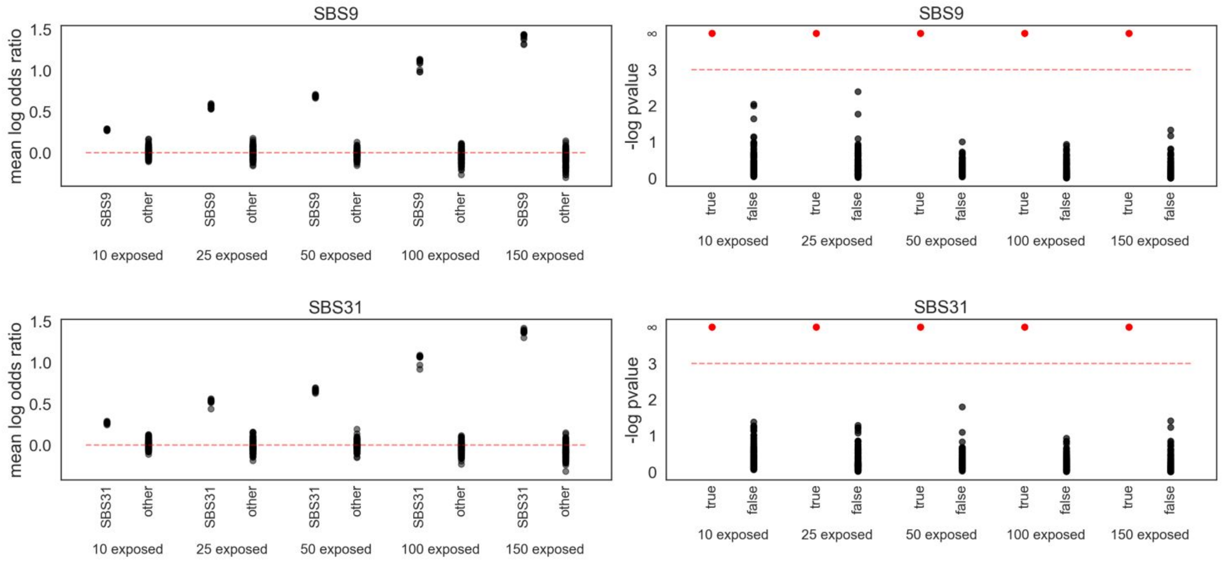


#### b) Breast-Lung-Colorectal

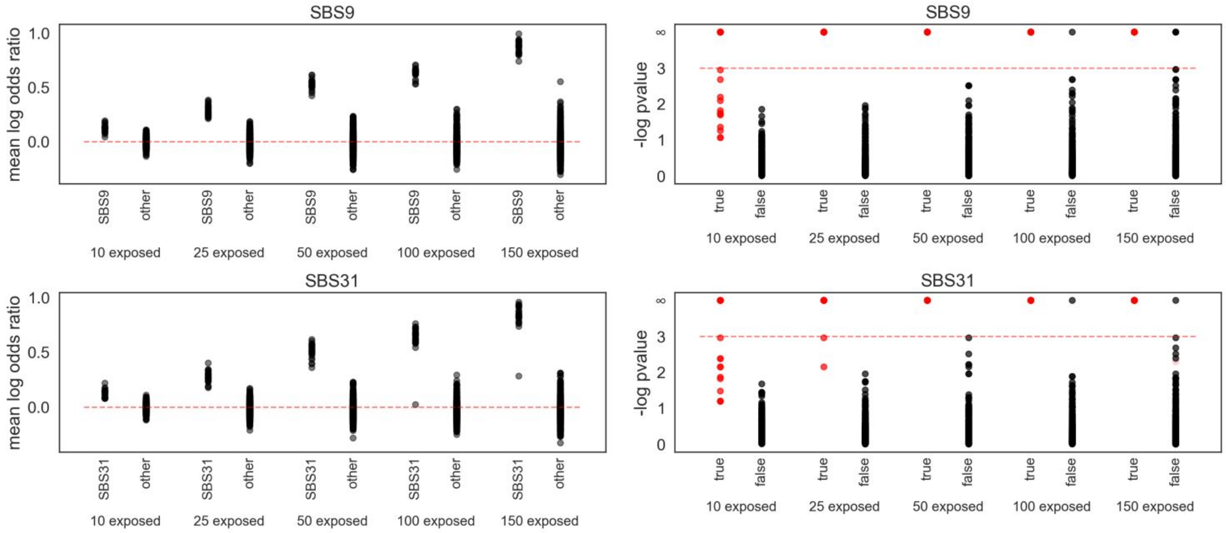


**Fig. 5.** Mean log-odds-ratios and significance values upon the regression method described in Methods as “step 1”. Red points (right plots) correspond to the associations of the injected labels to the reconstructed signature of the injection, thus true associations.

#### a) Breast

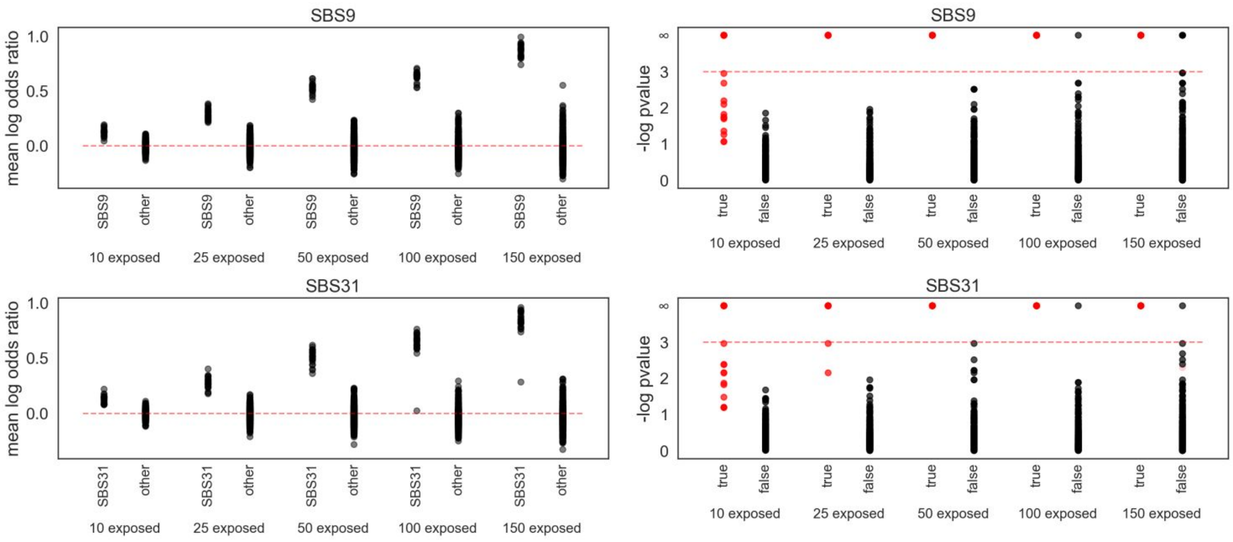


#### b) Breast-Lung-Colorectal

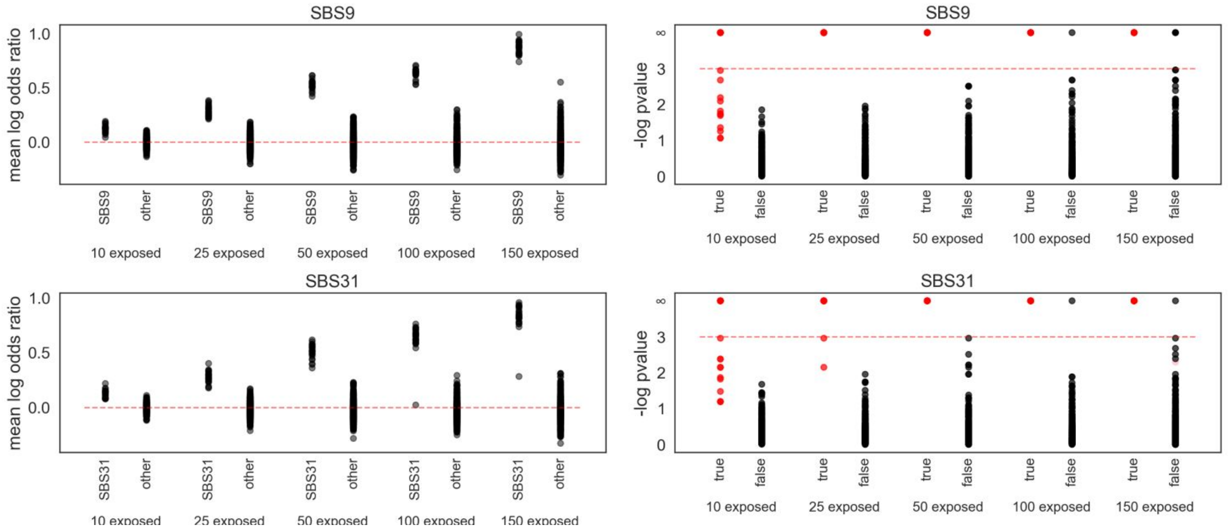


**Fig. 6.** Volcano plots that recapitulate the relationship between significance and effect-size (as in the main Figure 1e). All tests for all number of exposed samples and replicates are represented in the same plot. Red points (right plots) correspond to the associations of the injected labels to the reconstructed signature of the injection, thus true associations.

#### Introducing errors in treatment labels

As mentioned above, the regression method operates in a scenario that could contain some errors in the labels assigned to samples. Some patients in the cohort analyzed may be false positives --i.e., they are labeled as treated, but the treatment may have left no footprint due to low doses or short administration times-- or false negatives --i.e., no mention of treatment in the clinical data, but they have actually received it, resulting in mutational footprint-- hence complicating the fitting. Regarding false positives, due to the fact that our method to generate synthetic datasets draws the mutation burden for each sample randomly, our analysis already included some treated samples with a relatively small injected contribution. To test the impact of false negative errors, we relabelled so that all the treated samples had some chance (5%, 10% and 20%) to miss the label, then we run the method with the new labels (Figs. 7a and b).

#### Conclusions

When mutations of the foreign signature are injected to 25 or more samples of the synthetic dataset the signature reconstructed explains the labels of treated samples with high significance ($p < 0.001$). Furthermore, this association is robustly recovered even when an important fraction (up to 20%) of samples is mislabelled.

**a) Breast**

### 5% shuffle

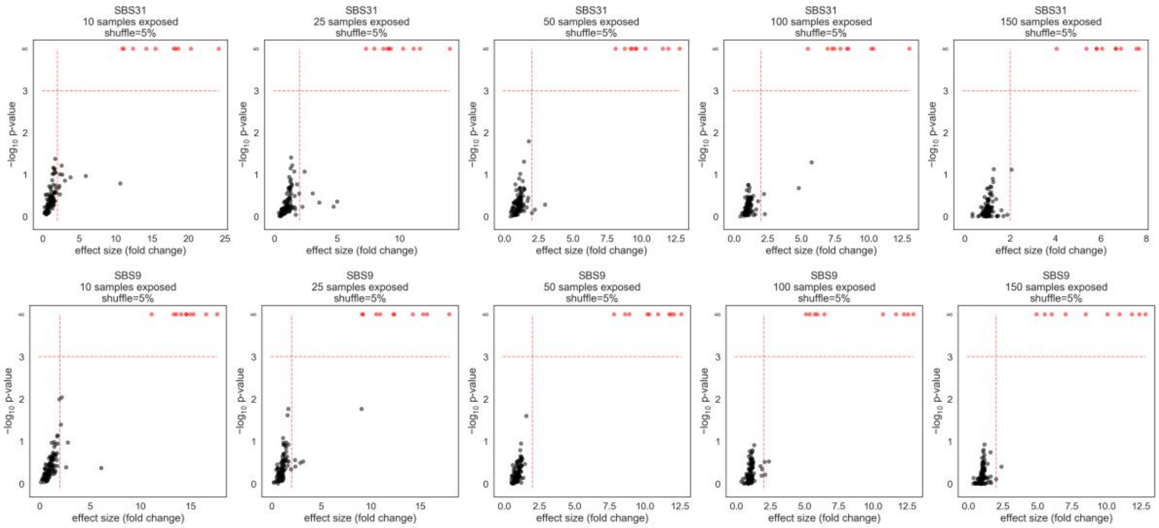


### 10% shuffle

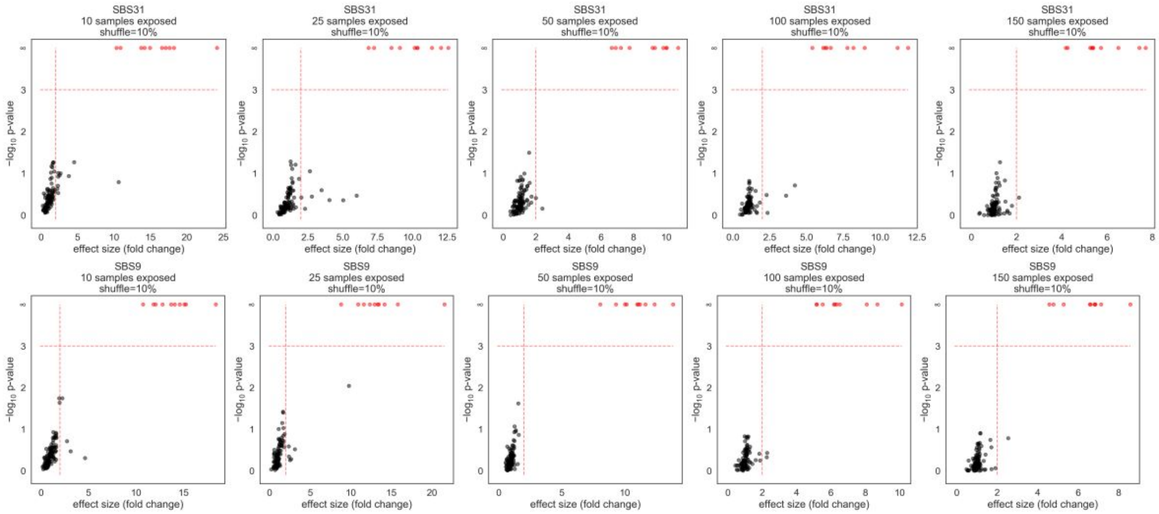

### 20% shuffle

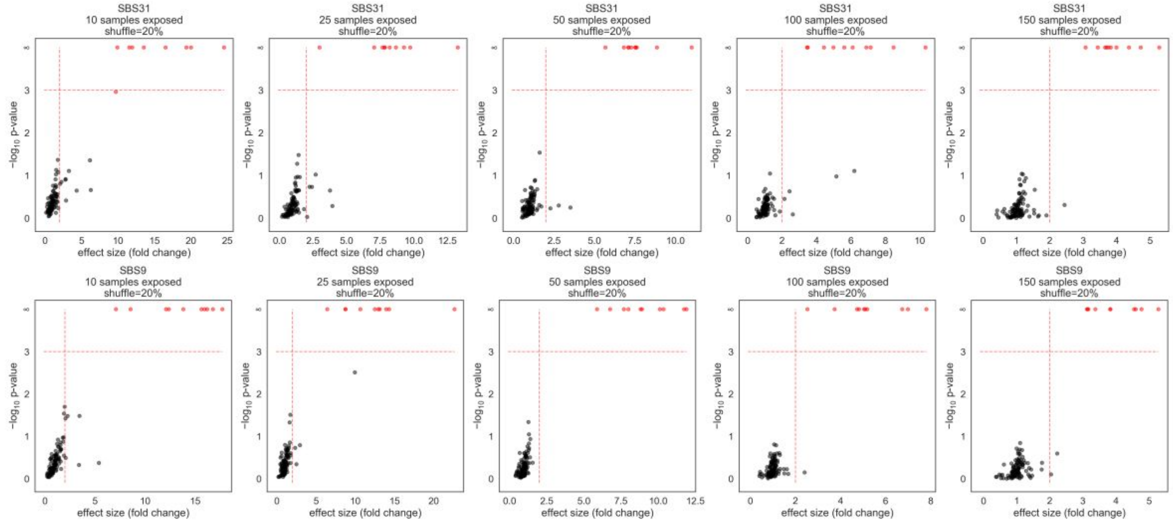


**b) Breast-Lung-Colorectal**

### 5% shuffle

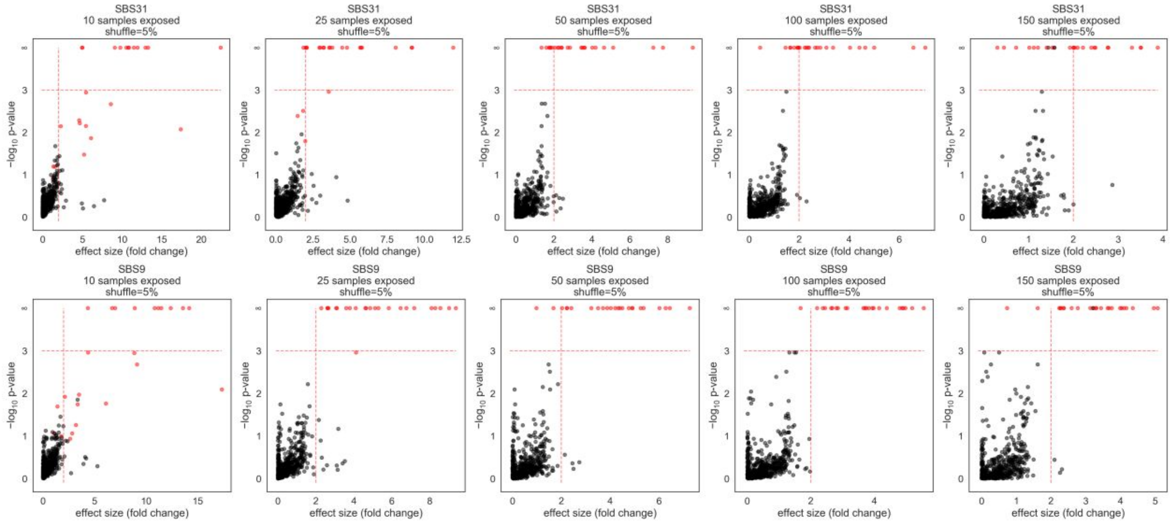

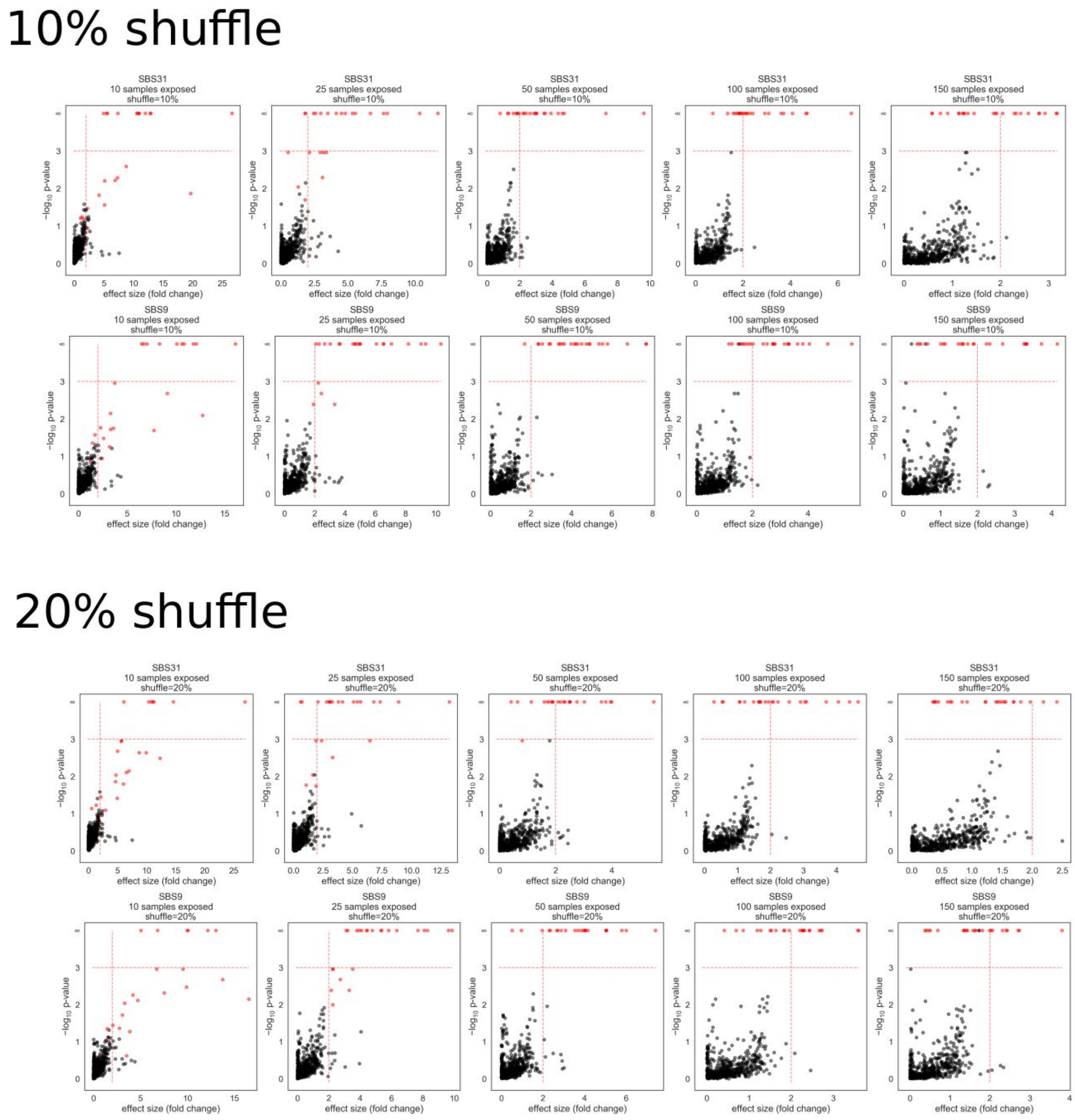


**Fig. 7.** Volcano plots that recapitulate the relationship between significance and effect-size (as in the main Figure 1e) at fitting in which some injected samples have forgotten their label, with rates of 5%, 10% and 20% forgotten labels among the true injected samples. For each number of samples exposed we provide a different plot.

### 5. Co-treatment analysis: injected signal against placebo

Following our previous analysis, it is possible that one signature turns out to be significantly associated to two or more treatments simply because one of them is causing the signature exposure and the rest, even if they do not produce any effect, substantially overlaps the true causal label -- in other words, we can witness hitchhiking effects. This is the case of many anti-cancer treatment regimens, which are often concomitantly administered.

In order to identify the most likely cause of the observed footprint among a set of concomitantly administered treatments with significant association, we described a method (see Methods) to assign with maximum parsimony an average efficiency to each competing factor (cotreatment).

We devised a series of tests with our synthetic datasets to clarify the limits of this methodology to detect the real causing factor of the observed footprint. In each test we allocate new concomitant treatment annotations to the samples (placebos) and check whether our method can correctly infer the relative efficiency of the injected exposure compared to the placebo.

For each catalogue, we can randomly allocate a new treatment --referred to as "placebo"-- to the samples, requiring that the true labels of the placebo overlaps a given percentage (R) of the samples with injected exposure. For each test, we annotated 10 different placebo treatments in this way, in addition to the treatment annotation related to the injected exposure.

For the sake of testing different data configurations, we conducted the placebo allocation in two ways: i) randomly allocating as many true placebos in the non-injected as in the injected samples (symmetric); ii) randomly allocating as many true placebos as the total number of samples with injected exposure (asymmetric).

The cotreatment analysis described in Methods assigns a mean efficiency value $E(t)$ to each treatment $t$ among a set of competing treatments that may be causing a footprint (of signature $S$). With each data configuration described above, we tested two ways of computing the relative efficiency between the true associated treatment (injection) and the placebos, alone or in combination:

- In the *one-to-one* comparison we make each placebo compete with the injected signal (true synthetic etiology) one at a time.
- In the *many-to-one* comparison we incrementally add one placebo to the previous ones and examine the average efficiency attributable to all the placebos in combination and the etiology.

In summary, we create 10 new placebo label assignments in either symmetric or asymmetric way, for 3 different overlap values $R = 25,\ 50,\ 75$ and we estimate the relative efficiency of the injected exposure compared to the placebo in either one-to-one or many-to-one setting.

#### Remark:

Note that each test will produce a value representing the log relative efficiency between the true causal treatment and the placebos (log fold change between exposure explained by true causal treatment vs exposure explained by placebos). Therefore, relative efficiencies above zero indicate that the method correctly recovers the true etiology. As we are conducting several tests for each specific setting, we represent the distribution of relative efficiencies as vertical violin-shaped plots.

#### Analyses:

##### Symmetric one-to-one setting

We examine the fold change between the mean efficiency values of the etiology and the placebo assignments (Figs. 8a and b).

##### Symmetric many-to-one setting

We examine the fold change between the mean efficiency values of the etiology and the placebo assignments (Figs. 9a and b).

##### Asymmetric one-to-one setting

We examine the fold change between the mean efficiency values of the etiology and the placebo assignments (Figs. 10a and b).

##### Asymmetric many-to-one setting

We examine the fold change between the mean efficiency values of the etiology and the placebo assignments (Figs. 11a and b).

##### Conclusions

The method for assessing relative efficiencies of the true etiology versus randomly generated placebos with predefined overlaps proved to yield sound results. Regarding the comparison between symmetric and asymmetric overlap settings, disentangling injection from the asymmetric setting proved more challenging -- leading to false negatives.

Among the two variants of the method we tested --first by comparing the true injection with one placebo at a time (one-to-one), second by comparing the true injection with all placebos at once, i.e., including them all in the design matrix (many-to-one) -- the many-to-one produced a large proportion of false negatives, particularly in the asymmetric setting. Interestingly, while the many-to-one approach can in principle cope with more complex situations in which mixtures of etiologies can compete with each other, in practice the lack of sensitivity completely undermines any potential advantages. On the other hand, the one-to-one approach has higher sensitivity and it also proves more robust in terms of the overlap.

**Figure 8**

**a) Breast**

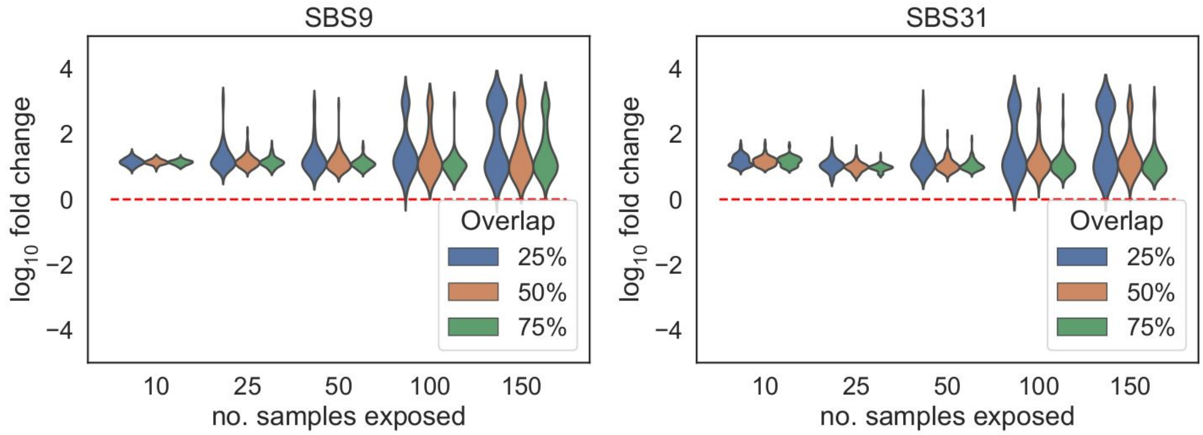


**b) Breast-Lung-Colorectal**

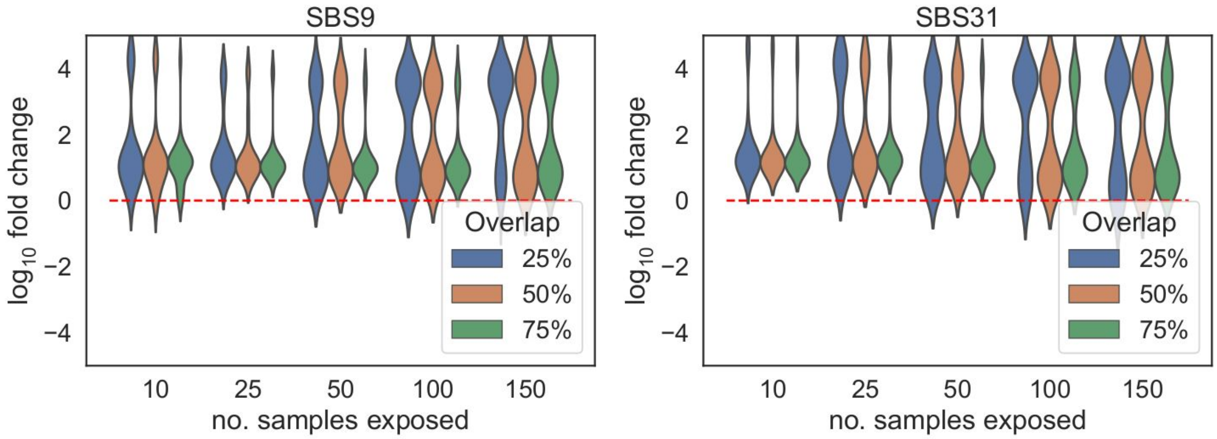

**Figure 9**

**a) Breast**

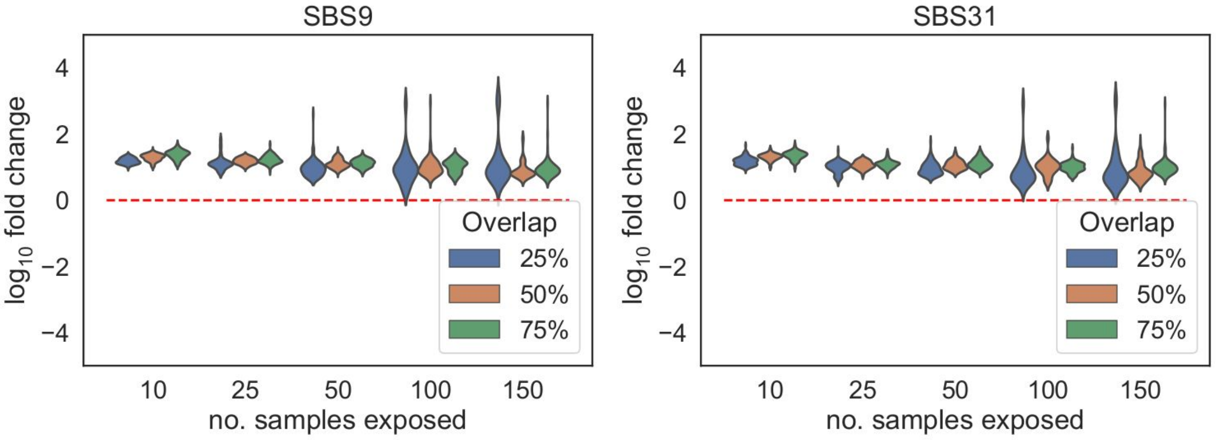


**b) Breast-Lung-Colorectal**

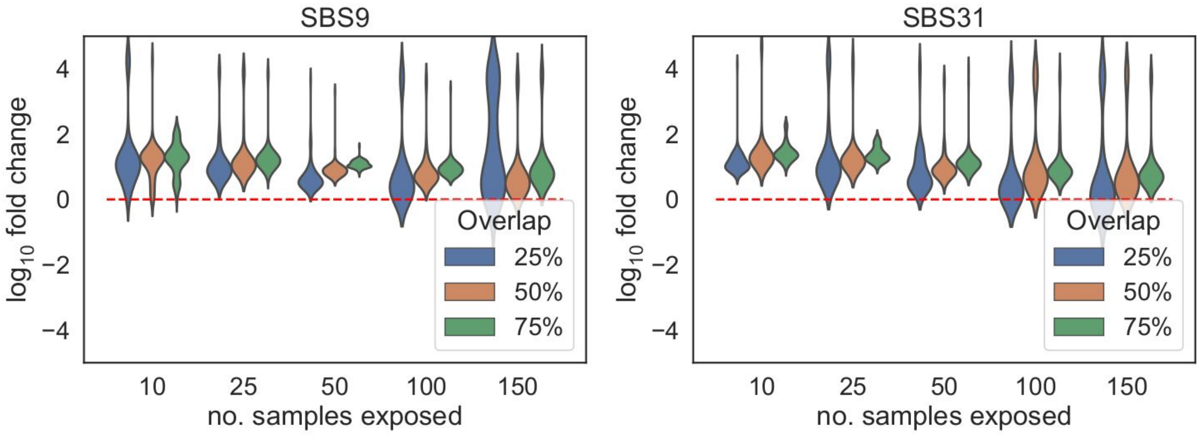

**Figure 10**

**a) Breast**

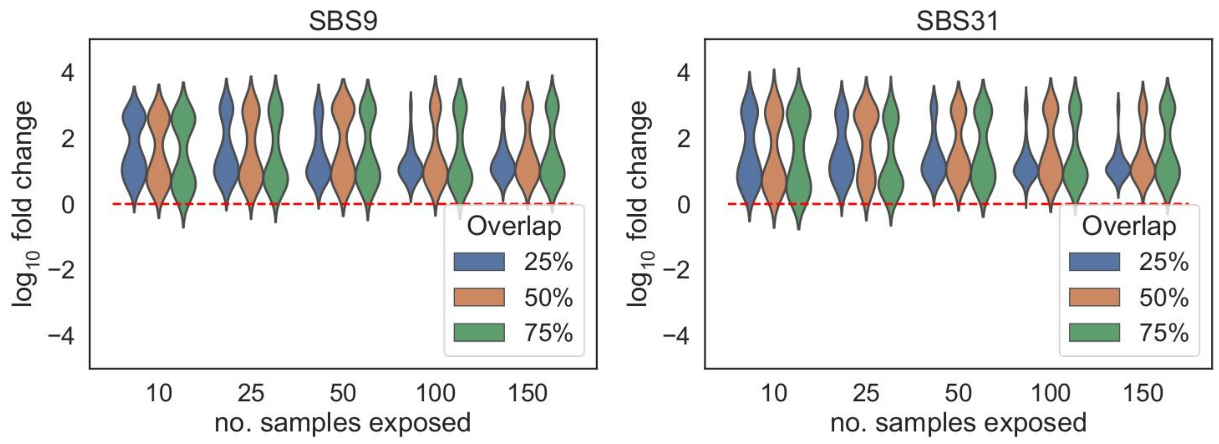


**b) Breast-Lung-Colorectal**

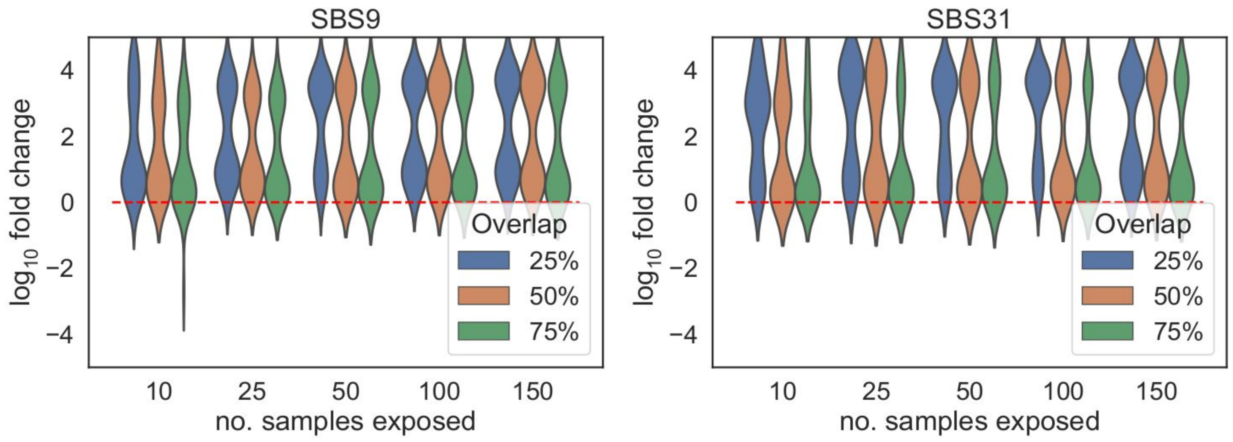

**Figure 11**

**a) Breast**

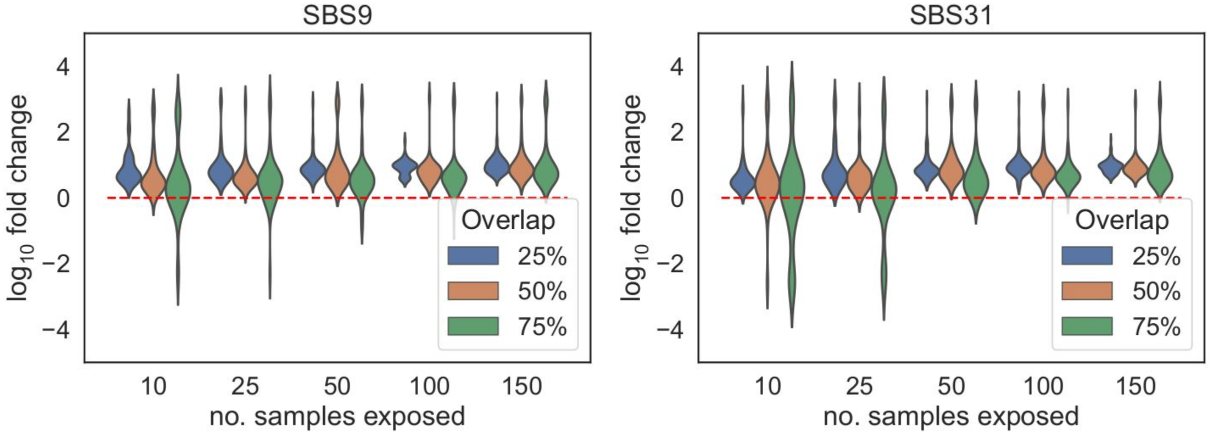


**b) Breast-Lung-Colorectal**

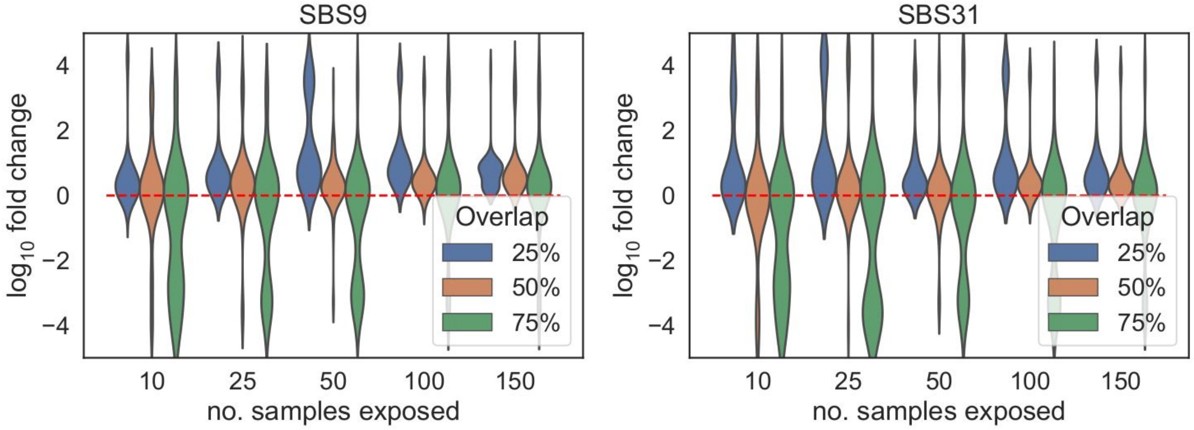


**Figs. 8, 9,10, 11.** Distribution of fold changes for all cotreatment tests with random placebos that overlap the injected labels with a given rate. According to the strategy to assign placebo labels (symmetric/asymmetric) and to infer relative efficiencies (one-to-one/many-to-one) the figures can be tagged as: symmetric/one-to-one (Fig. 8); symmetric/many-to-one (Fig. 9); asymmetric/one-to-one (Fig. 10); asymmetric/many-to-one (Fig. 11).

#### 6. Recovery of the activity of injected signatures

Next we compared the activity of the reconstructed injected signals against the true number of mutations injected to each sample. We provide scatter plots (Figs. 12 and 13) illustrating the relationship between the reconstructed activity and the number of mutations injected within a window of 5000 mutations for each specific configuration.

For each sample, we can also measure the discrepancy between the mutation burden contributed by the injected signature and the exposure of the reconstructed signature using relative residuals:

$$\varepsilon_r(s) = \frac{I(s) - R(s)}{I(s)}$$

where $I(s)$ is the injected number of mutations and $R(s)$ is the reconstructed number of mutations for the extracted signature in sample $s$. In a sample with a relative residual close to 0, the estimation of the activity of the injected signature carried out as part of the extraction is very close to the actual number of injected mutations. For each synthetic dataset we represented the values $\varepsilon_r(s)$ for all the injected samples $s$ (Figs. 14a and b), the median median $\varepsilon_r$ across all samples for each replicate (Figs. 15a and b) and finally the concordance correlation coefficient [3] (Figs. 16a and b), which captures the overall statistical reproducibility of the injected burden by the reconstructed exposures at the injected samples -- values close to 1 mean high reproducibility.

##### Conclusions

The computed mean relative error across synthetic datasets demonstrates that when the injected signature is active in more than 50 samples, the extraction method estimates the number of injected mutations with a reasonable accuracy. Specifically, the relative residuals are well below 25% for most of the samples. The reproducibility of exposure is generally very high, although some random replicates still produce low concordance correlation values (Fig. 16b). This fluctuations are likely driven by the dispersion of the number of mutations per sample in the log scale, which underscores the impact of the cohort composition when it comes to signature extraction. This provides support for the estimation of the mutation burden contributed by treatments carried out.

SBS 9

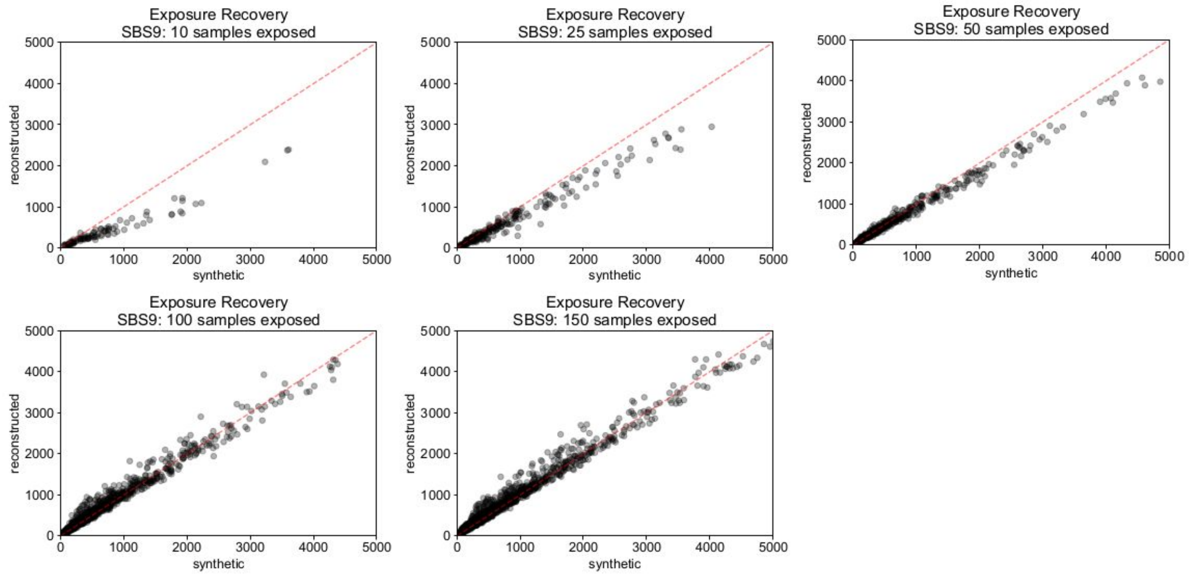


SBS 31

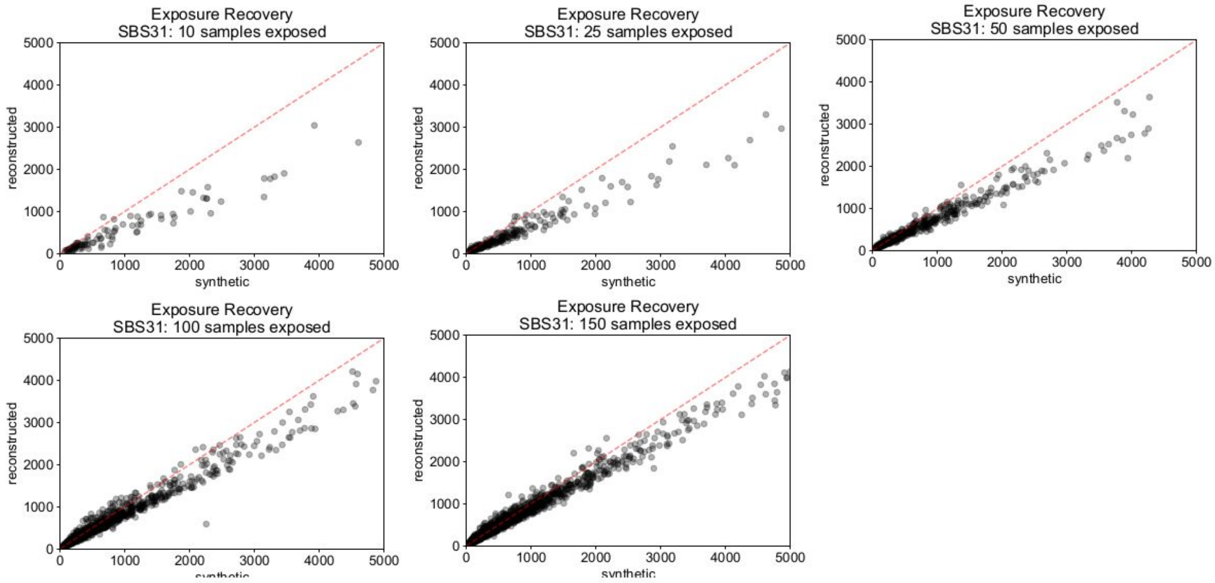


**Fig. 12.** Breast. Scatter plots of injected burden versus reconstructed exposures per sample across replicates.

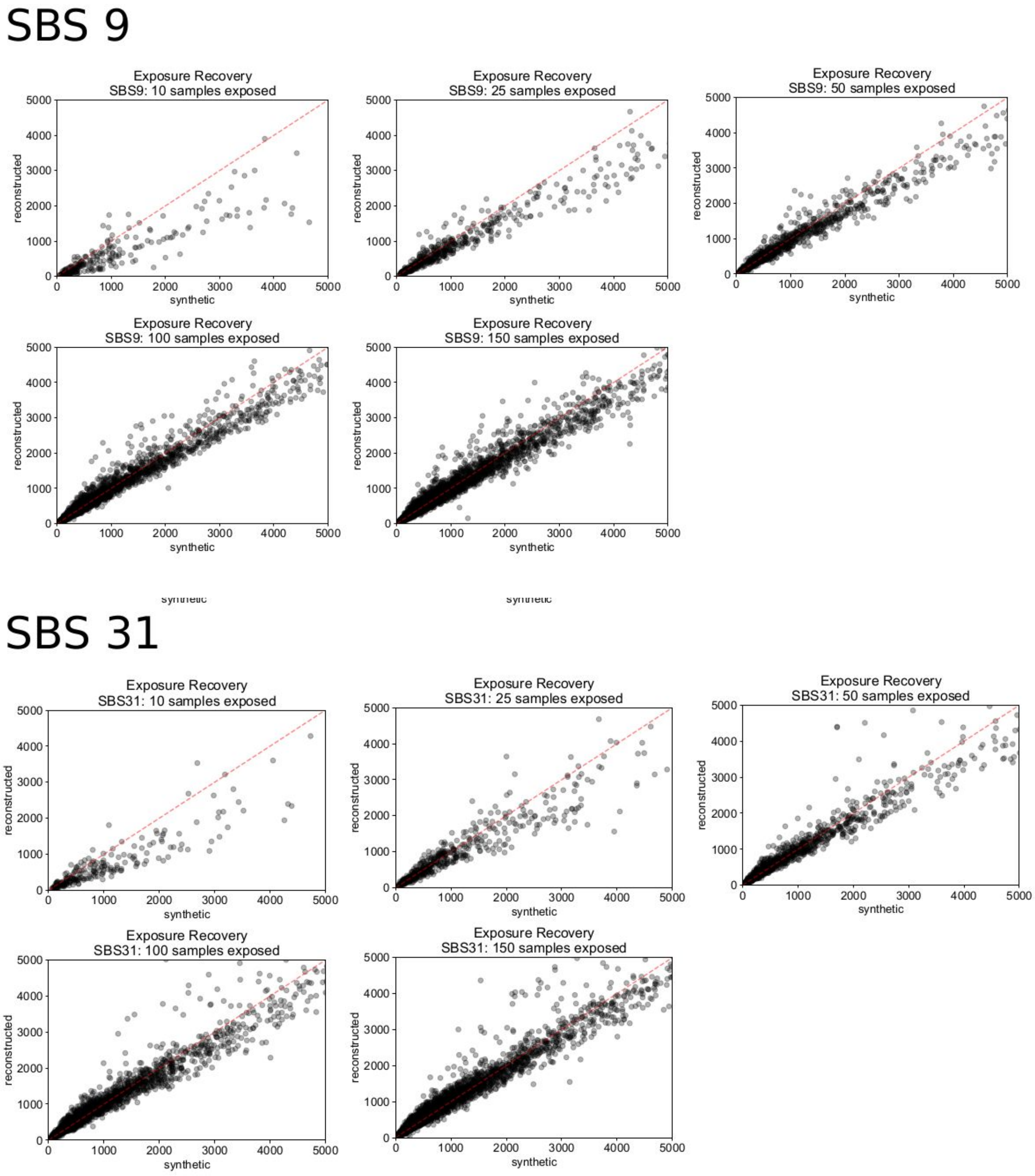


**Fig. 13.** Breast-Lung-Colorectal. Scatter plots of injected (synthetic) burden versus reconstructed exposures per sample across replicates.

**a) Breast**

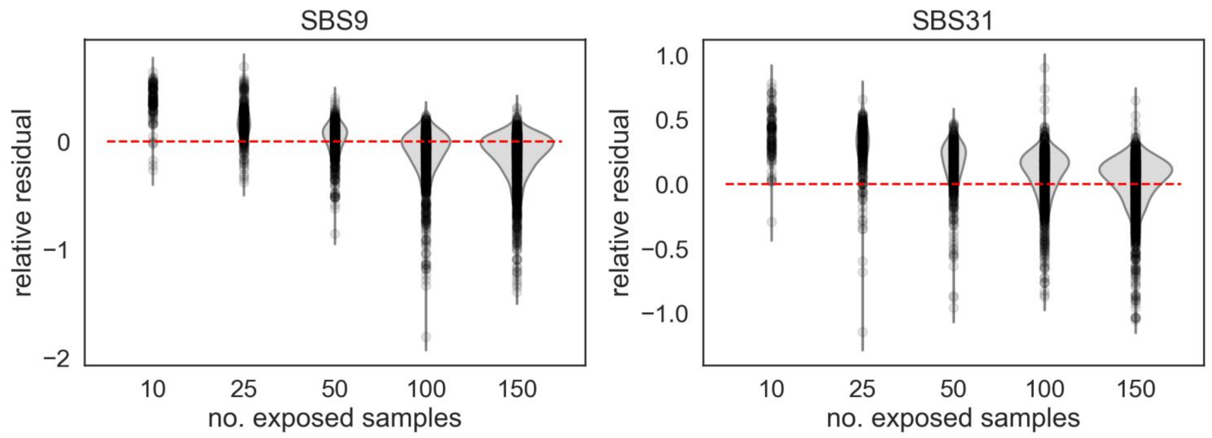


**b) Breast-Lung-Colorectal**

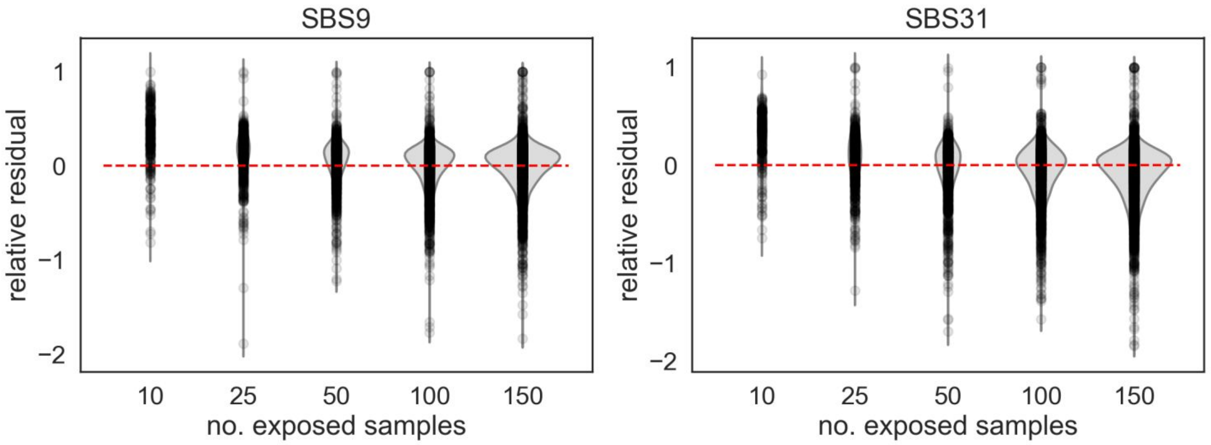


**Fig. 14.** Distribution of relative residuals (signed) for all the samples injected and all the replicates per number of samples exposed. When relative residuals are close to 0, the reconstructed exposure in similar to the injected burden. Relative residuals > 0 (resp. < 0) indicate that the true injected burden is higher (resp. lower) than the reconstructed exposure. As the number of samples exposed increases, the relative residuals concentrate more about 0.

**a) Breast**

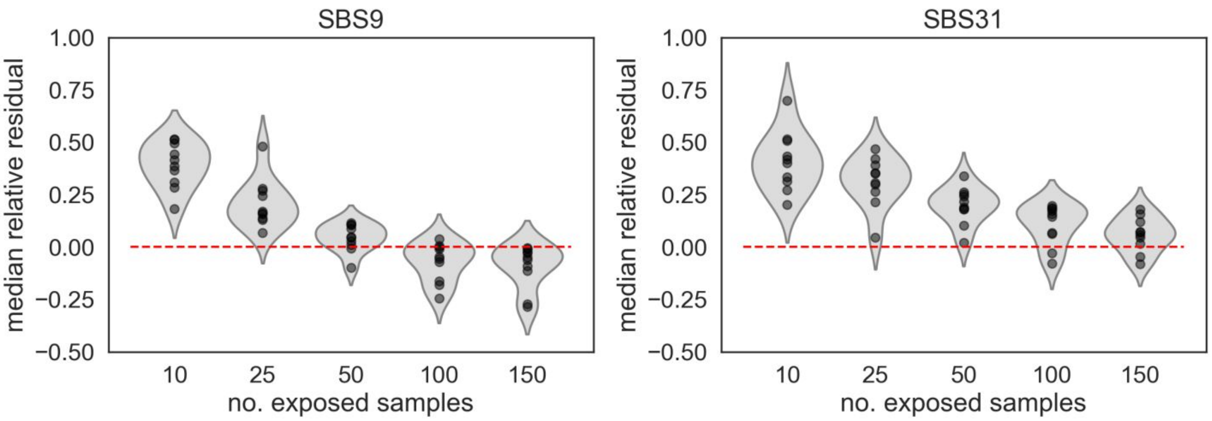


**b) Breast-Lung-Colorectal**

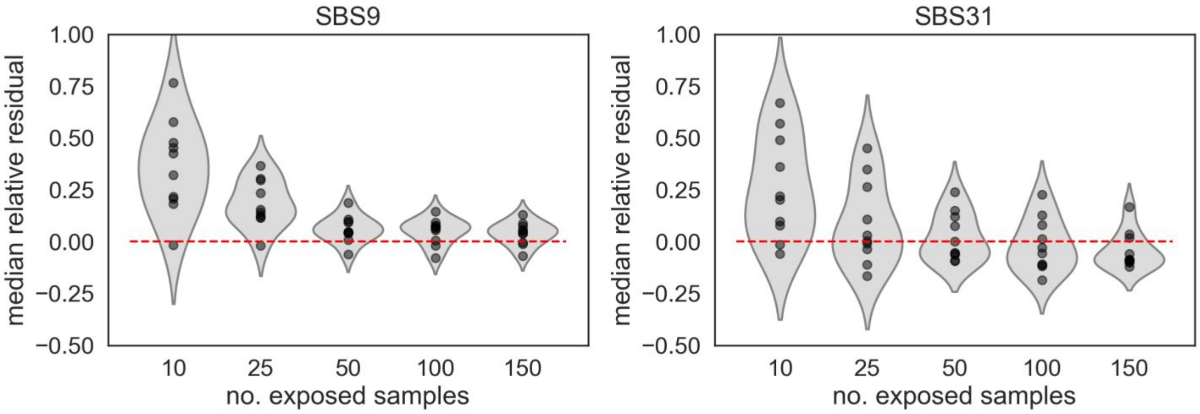


**Fig. 15.** Mean relative residuals per replicate. When mean relative residuals are close to 0, the reconstructed exposure in similar to the injected burden.

**a) Breast**

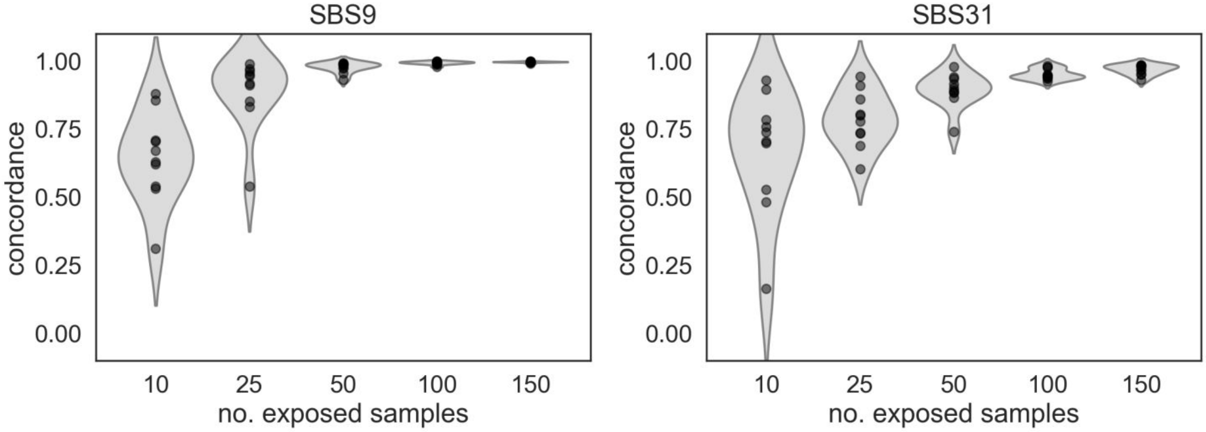


**b) Breast-Lung-Colorectal**

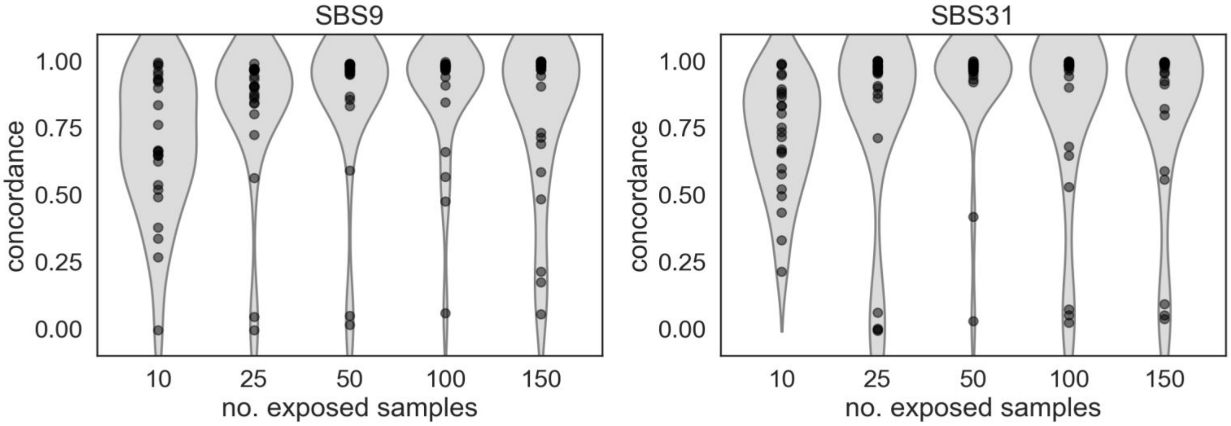


**Fig. 16.** Concordance correlation coefficient per replicate. Values close to 1 indicate high concordance between the injected burden per sample and the reconstructed exposure per sample.

#### 7. Signature activity in non-treated samples

We were also interested in assessing the appearance of activity of a signature in samples in which no mutations of that signature were injected. This can be due to the activity of the signature in the primary tumor or due to “bleeding”, i.e., the artifact whereby some signatures may gain or lose mutations to other processes as part of the NMF deconstruction [1].

We thus computed the ratio between the average reconstructed activity across non-injected samples and the average injected number of mutations across samples exposed to injection (average exposure over average injected burden in Figs. 17a and b). A value of 1 of this metric means that on average the same activity of the injected signature is detected in injected and non-injected samples. Therefore, the closer to 0 this value is, the smaller the effect of “bleeding” of the injected signature into non-injected samples.

##### Conclusions

While some fluctuations arise at some replicates, the amount of mutations attributed to the injected signal in non-injected samples is generally low and stable, with average fold change (bleeding) $< 0.1$ (Figs. 17a and b).

**a) Breast**

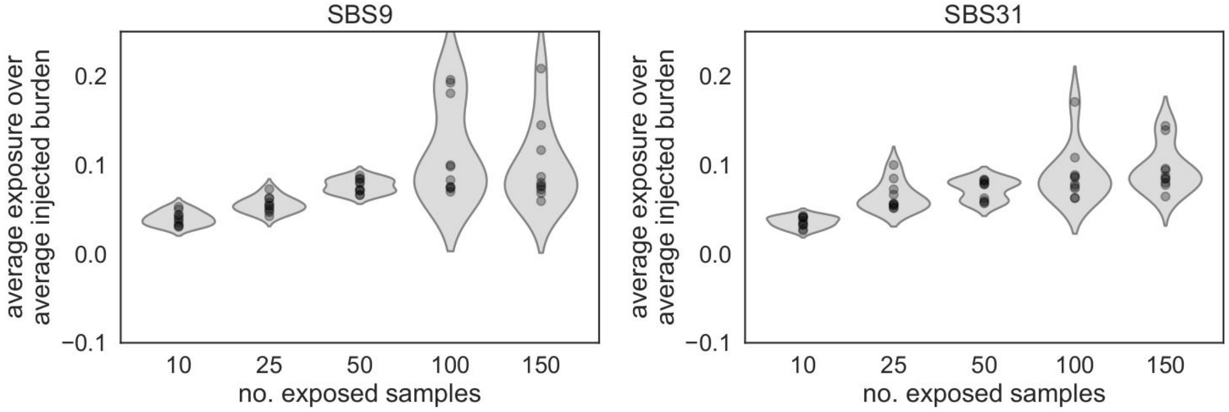


**b) Breast-Lung-Colorectal**

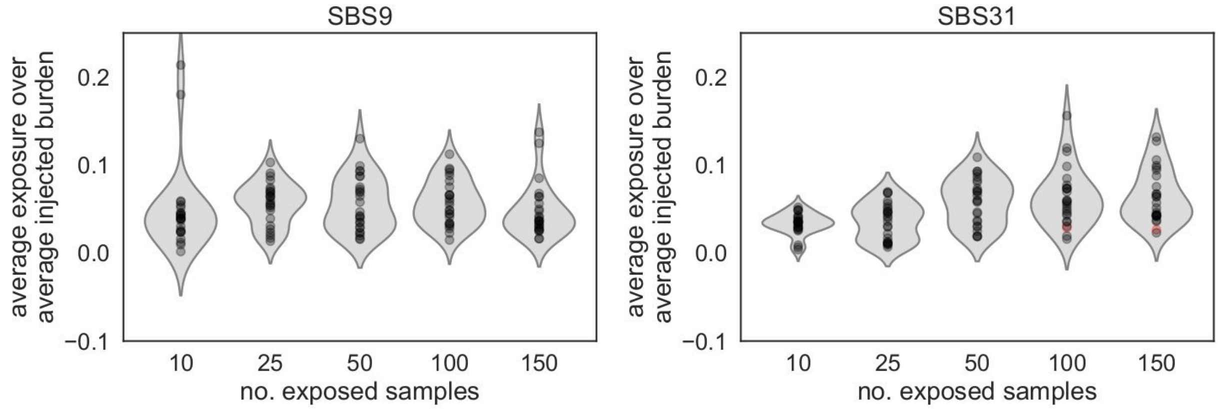


**Fig. 17.** Burden of the injected signature on non-injected samples given terms of the injected burden.
